# Experimental evolution to thermal stress indicates climate resilience in a cosmopolitan arthropod

**DOI:** 10.64898/2026.01.16.699875

**Authors:** Gaoke Lei, Huiling Zhou, Zongyao Ma, Yating Duan, Yanting Chen, Fengluan Yao, Minsheng You, Liette Vasseur, Geoff M. Gurr, Shijun You

**Affiliations:** State Key Laboratory of Agriculture and Forestry Biosecurity, Institute of Applied Ecology, Fujian Agriculture and Forestry University; Institute of Plant Protection, Fujian Academy of Agricultural Sciences, Fuzhou, China; International Joint Research Laboratory of Ecological Pest Control, Ministry of Education, Fujian Agriculture and Forestry University, Fuzhou, China; Ministerial and Provincial Joint Innovation Centre for Safety Production of Cross-Strait Crops, Fujian Agriculture and Forestry University, Fuzhou, China; Department of Biological Sciences, Brock University, St. Catharines, Canada; Gulbali Institute, Charles Sturt University, Orange, NSW, Australia

**Author notes:** **For correspondence:** (SY). These authors contributed equally to this work (Gaoke Lei, Huiling Zhou).

**Keywords:** Climate change, Temperature tolerance, Multi-omic analysis, Genetic adaptation

## Abstract

Adaptive evolution enables species to survive and thrive under changing environmental conditions. In the face of accelerating global climate change, thermal stress represents a major challenge to the persistence of terrestrial arthropods. Understanding the genetic mechanisms underlying thermal adaptation is therefore critical for predicting species’ evolutionary potential and future success. Here, we combine experimental evolution, phenotypic assays, and multi-omics analyses to investigate the adaptive responses of the diamondback moth (*Plutella xylostella*), a globally destructive pest of cruciferous crops, to contrasting thermal environments. Populations evolved under hot (32°C/27°C) and cold (15°C/10°C) regimes exhibited distinct life history and fitness traits relative to those maintained under favorable conditions (26°C). The hot strain showed accelerated development, higher fecundity, and increased survival under extreme heat, while the cold strain exhibited lower supercooling and freezing points, indicating enhanced cold hardiness. Integrated transcriptomic and metabolomic analyses revealed extensive transcriptional reprogramming and convergent metabolic adjustments, notably a reduction in lipid metabolism to conserve energy under thermal stress. Crucially, non-synonymous mutations in *PxSODC* enhance superoxide scavenging efficiency, enabling effective oxidative stress management at lower gene expression levels. Furthermore, we identified epigenetic regulation via DNA methylation as a key mediator of this thermal tolerance. Together, these coordinated mutational, epigenetic, and metabolic insights highlight this arthropod’s capacity for global dispersal and likely persistence under climate change, establishing a framework for understanding equivalent effects in other species.

## Introduction

Human-induced climate change, particularly the continued change in temperature and precipitation patterns (***IPCC, 2023***), is altering the geographical distribution of insect pests, allowing those previously confined by temperature barriers to spread to new areas and posing growing threats to crop production and food security (***Deutsch et al., 2008*; *Outhwaite et al., 2022*; *Lawlor et al., 2024***). Such range expansion requires adaptation not only to warmer conditions in existing habitats but also to cold extremes encountered during colonization of higher latitudes or elevations (***Harvey et al., 2020***). As poikilothermic organisms with high surface area to volume ratios, insects are particularly responsive to temperature change (***Wang et al., 2022***), and species with higher genetic diversity and adaptive potential may be better equipped to cope with novel thermal environments, giving them an advantage in expanding into new habitats.

Adaptive evolution is a crucial mechanism for insect pests to expand their geographical ranges under climate change (***McCulloch and Waters, 2022; Burc et al., 2025***). Understanding the genetic basis of thermal adaptation is therefore essential for predicting how and ultimately where pest species will colonize new regions as temperature barriers shift (***Gibert et al., 2019***). Genetic mutations, particularly non-synonymous mutations, can alter protein structure or function and increase thermal tolerance (***Belfield et al., 2018***). For example, nonsynonymous mutations in the *AcVIAAT* gene of the eastern honeybee, *Apis cerana* (e.g., P42L substitution), are associated with enhanced thermal adaptation (***Li et al., 2024c***). and the alpine ground beetle, *Nebria vandykei,* achieve survival in extreme thermal environments through adaptive selection of mutations in key genes (*TREH*, *EIF3A*, *LRPPRC*, etc.) coupled with immediate responses from heat shock proteins (***Schoville et al., 2024***). Although significant progress has been made in identifying such mutations in non-model organisms, we do not yet know how long-term thermal selection drives coordinated changes across gene function, metabolic networks, and life history traits to enable thermal adaptation and range expansion in pest species.

The diamondback moth (DBM), *Plutella xylostella* (Lepidoptera: Plutellidae), is a globally distributed pest of cruciferous crops thriving across a wide range of climatic conditions (***Furlong et al., 2013***). Genome-wide SNP analysis of field populations from 114 locations revealed climate-adaptive genetic variability, suggesting that *P. xylostella* can tolerate projected future climates in most regions (***Chen et al., 2021***). These features, together with the availability of complete genome sequences and extensive SNP datasets (***You et al., 2013*; *You et al., 2020***), make it an ideal model for studying the genetic basis of thermal adaptation through integrated multi-omics approaches. Here, we investigate the mechanisms of *P. xylostella*’s genetic adaptation and evolutionary responses to different thermal environments. Specifically, we employ thermal regime patterns of 12h/12h in hot (32°C/27°C) or cold (15°C/10°C) environments over the course of three years (∼75 and ∼15 generations for the hot and cold strains, respectively), compared to the favorable constant condition at 26°C, to investigate the adaptive evolution of *P*. *xylostella* in climate-controlled chambers.

Age-stage, two sex life tables (a demographic method that simultaneously incorporates age, developmental stage, and both sexes; Chi, 1988) of *P*. *xylostella* measured the life history variation of the three *P*. *xylostella* strains evolved in the favorable (ancestral), hot and cold environments. Using metabolomic and transcriptomic analyses, we identified the key genes that could facilitate the adaptation of *P*. *xylostella* to thermal extremes. Our results showed that a large number of differentially expressed genes and metabolites were produced in populations adapted to high and low temperatures through multi-generational selection. We find the mutant of a key gene, *PxSODC*, which can alter the superoxide dismutase activity and increase the ability to scavenge superoxide anions. Since extreme temperatures elevate intracellular reactive oxygen species that damage cellular structures, this enhanced scavenging capacity helps maintain cellular homeostasis, thereby significantly affecting the adaptability under high and low temperature environments. CRISPR-Cas9 was used to functionally validate the role of *PxSODC* in facilitating adaptive evolution and influencing regulatory networks. These findings demonstrate that long-term thermal selection drives coordinated transcriptomic, metabolic, and life history divergence in *P. xylostella*, and identify non-synonymous mutations in *PxSODC* that enhance superoxide scavenging efficiency as a key genetic mechanism underlying thermal adaptation, providing a framework for predicting its population dynamics under global climate change.

## Results

### Life history trait divergence among temperature-adapted strains

Following three years of evolution under contrasting thermal regimes (∼75 and ∼15 generations for the hot and cold strains, respectively), the hot strain (HS), cold strain (CS), and ancestral strain (AS) exhibited divergent life history traits. The hot strain exhibited an accelerated life cycle and increased fecundity, while the cold strain had extended male longevity. Specifically, both hot and cold strains had a significantly shorter preadult duration than the ancestral strain. The hot strain also had significantly shorter female longevity, and oviposition days, while the cold strain had a longer male longevity when compared to the ancestral strain (***Supplementary File 1***). The female fecundity, population intrinsic rate of increase (*r*) and finite rate of increase (λ) were all significantly higher in the hot strain than the cold strain which was not significantly different than the ancestral strain (***Supplementary File 1***). Detailed age-stage survival and fecundity curves are provided in Appendix 1.

To assess the evolved thermal tolerance of the temperature-adapted strains, we further examined the stage-specific survival rates of the hot and ancestral strains under extremely high temperatures, as well as the supercooling and freezing points of the cold and ancestral strains at pupae stage. The survival rates of eggs, 3^rd^-instar larvae and adults in the hot strain were higher than those of the ancestral strain at 42°C (e.g., 3^rd^-instar larvae at 120 min: HS 26.67% ± 3.57% vs. AS 13.33% ± 2.47%), and the survival rate of pupae in the hot strain was higher than that in the ancestral strain at 43°C and 44°C (***Figure 1A***). The supercooling and freezing points of pupae in the cold strain (supercooling: -23.99 ± 0.18°C; freezing: -14.24 ± 0.61°C) were significantly lower than those in the ancestral strain (supercooling: -23.09 ± 0.26°C; freezing: -11.58 ± 0.52°C), with differences of 0.90°C and 2.66°C, respectively (***Figure 1B***). The variation in survival rates and the supercooling/freezing points at extreme temperatures suggest that the hot and cold strains of *P*. *xylostella* have undergone profound adaptive adjustments.

**Figure 1.**
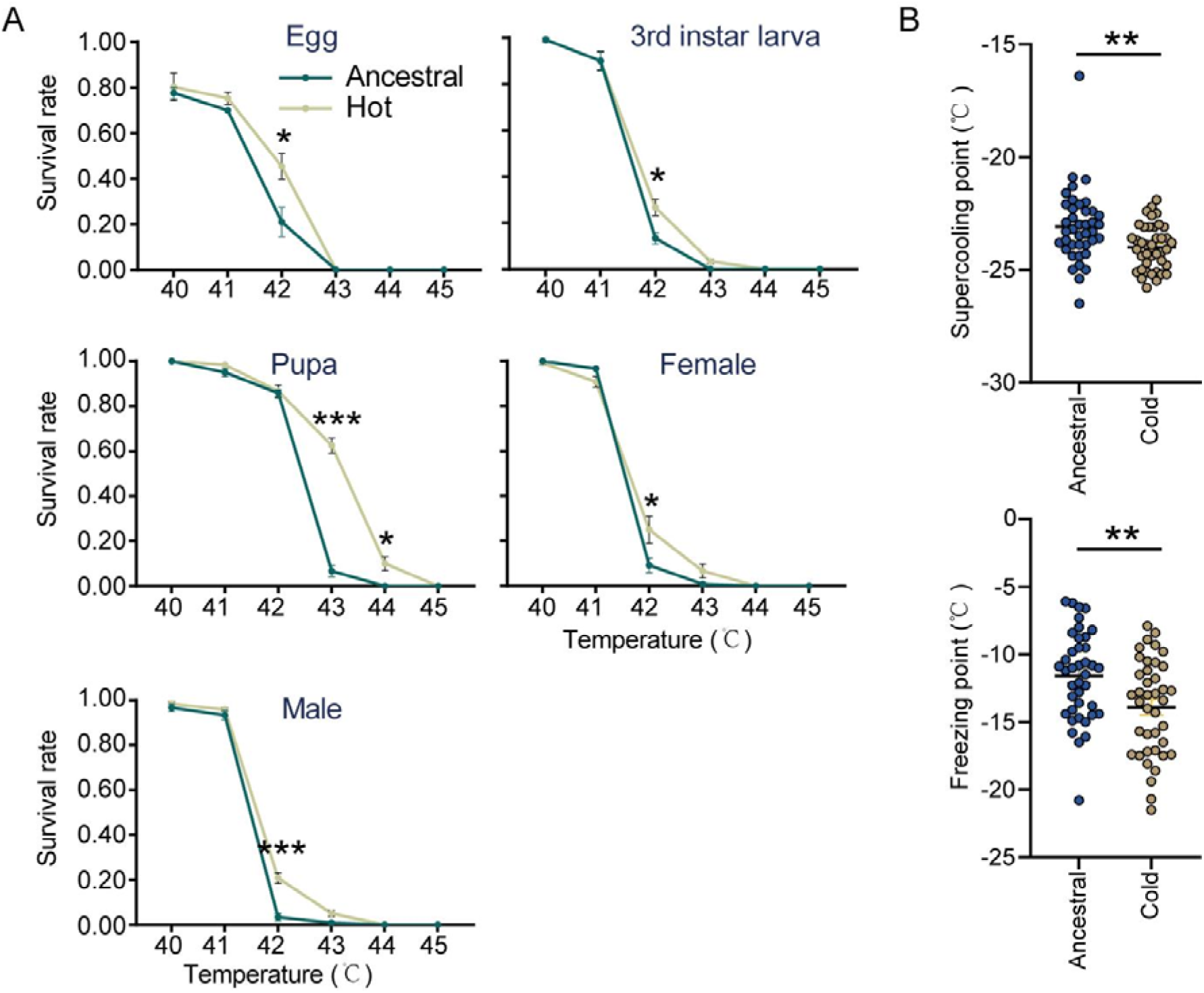
Evolved phenotypic changes in temperature-adapted strains (A) Stage-specific thermal tolerance responses (survival rates) of the ancestral and hot strains, with 20 individuals used in each of the six replicates for every treatment. (B) Supercooling and freezing points of pupae for the ancestral and cold strains, with 40 biologically independent samples used in each treatment. Data are presented as mean ± SEM. Statistical analyses are performed using t-tests with significant levels indicated by asterisks (\**p* < 0.05, \*\**p* < 0.01, \*\*\**p* < 0.001). **Source data 1.** Raw data for stage-specific thermal tolerance and pupal supercooling/freezing points of temperature-adapted strains.

### Omics-based evidence for adaptive evolution

Our previous studies have identified metabolites such as trehalose and very long chain fatty acids that play a role in adaptation of *P*. *xylostella* to both high and low temperatures (***Zhou et al., 2022; Lei et al., 2023***). We performed a broad analysis of targeted metabolites of 3^rd^-instar larvae of each strain using high-throughput UPLC-MS/MS. A total of 781 metabolites were identified, including 199 amino acids and their metabolites, 146 lipids, 90 organic acids and their derivatives, 78 nucleotides and their metabolites, 61 heterocyclic compounds, 45 benzene and substituted derivatives, 42 alcohols and amines, 37 carboxylic acids and derivatives, 21 coenzymes and vitamins, and 62 other metabolites (***Figure 2A***). Principal component analysis (PCA) and inter-sample correlation heat maps revealed significant metabolic changes in the 3^rd^-instar larvae from the ancestral strain to the hot and cold strains (***Figure 2B; Figure 2–figure supplement 1A***). These comprised 77 differential metabolites (34 up-regulated, 43 down-regulated compared to the ancestral strain) in the hot strain, and 37 differential metabolites (13 up-regulated, 24 down-regulated compared to the ancestral strain) in the cold strain (***Figure 2C***).

**Figure 2.**
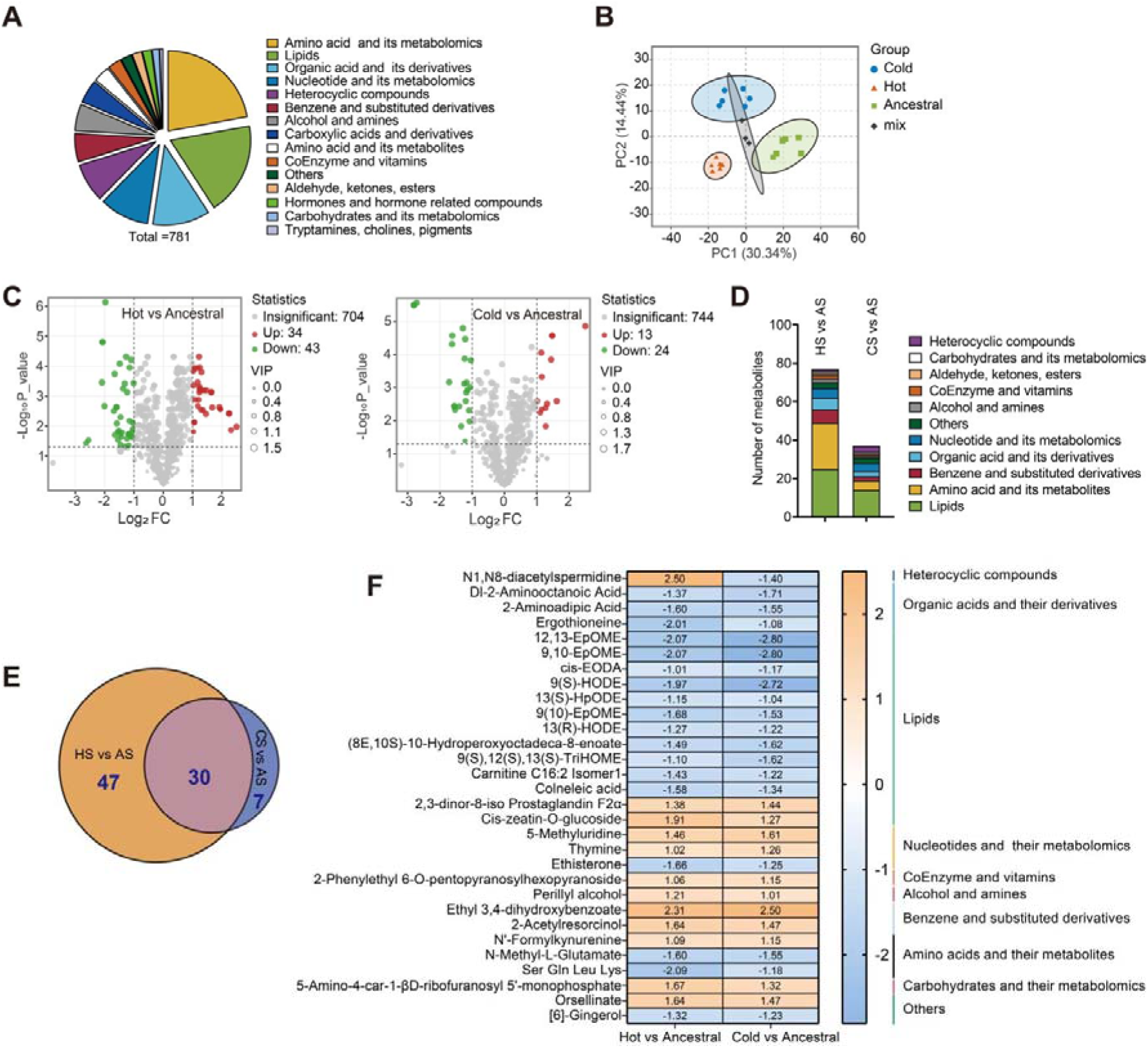
Metabolomic analysis of 3^rd^-instar larvae across ancestral, hot and cold strains (AS, HS and CS). (A) Classification of metabolites, with a total of 781 metabolites being identified in different strains. (B) Principal component analysis (PCA) of the 781 metabolites across different strains. PC1 and PC2 represent the first and second principal components, respectively. (C) Volcano plot showing the down-regulated (green dots) and up-regulated (red dots) metabolites based on comparison between HS/CS and AS. (D) Classification of differential metabolites between HS/CS and AS. (E) Venn diagram showing the common and unique differential metabolites in HS and CS as compared to AS. (F) Fold changes and classifications of the common differential metabolites in HS and CS as compared to AS.

Compared to the ancestral strain, the common differential metabolites of the hot and cold strains included lipids, amino acids and their metabolites, organic acids and their derivatives, nucleotides and their metabolites, and benzene and substituted derivatives (***Figure 2D; Figure 2–figure supplement 1B***). Inter-replicate analysis of differential metabolites showed a low correlation between the ancestral strain and hot/cold strains, but a high correlation between the hot and cold strains (***Figure 2–figure supplement 1C***). Notably, 30 common metabolites were identified across the differential sets based on comparison of the hot/cold strains to the ancestral strain (***Figure 2E***). These metabolites, except for N1, N8-diacetylpiperidine, exhibited similar fold changes when comparing the hot/cold strains to the ancestral strain (***Figure 2F***). These results indicate that *P*. *xylostella* responds to different environmental stresses by regulating similar metabolic pathways. Further analysis revealed a reduction in most of the lipid metabolites in both HS and CS compared to AS (***Figure 2F***).

We then profiled and compared the transcriptomes of the three strains to identify the key genes involved in adaptation of *P*. *xylostella* to temperature extremes. This revealed significant variation in gene expression among strains (***Figure 3A; Figure 3–figure supplement 1A***), with 1364 (825 up-regulated, 539 down-regulated) and 2029 (1205 up-regulated, 824 down-regulated) genes differentially expressed in the hot and cold strains, compared to the ancestral strain (***Figure 3B***). Pearson correlation showed, in contrast to the metabolomics data, a lack of strong correlation between the differentially expressed genes of the hot/cold strains and the ancestral strain (***Figure 3C***), with 498 common differentially expressed genes (***Figure 3D***). However, KEGG analysis revealed that the differentially expressed genes between the hot/cold strains and the ancestral strain were enriched in a substantial overlap of similar pathways, such as transport and catabolism, signal transduction, and lipid metabolism (***Figure 3E***), indicating that while multiple genes are involved in the adaptation of *P*. *xylostella* to high and low temperatures, a relatively limited range of biological functions might be affected.

**Figure 3.**
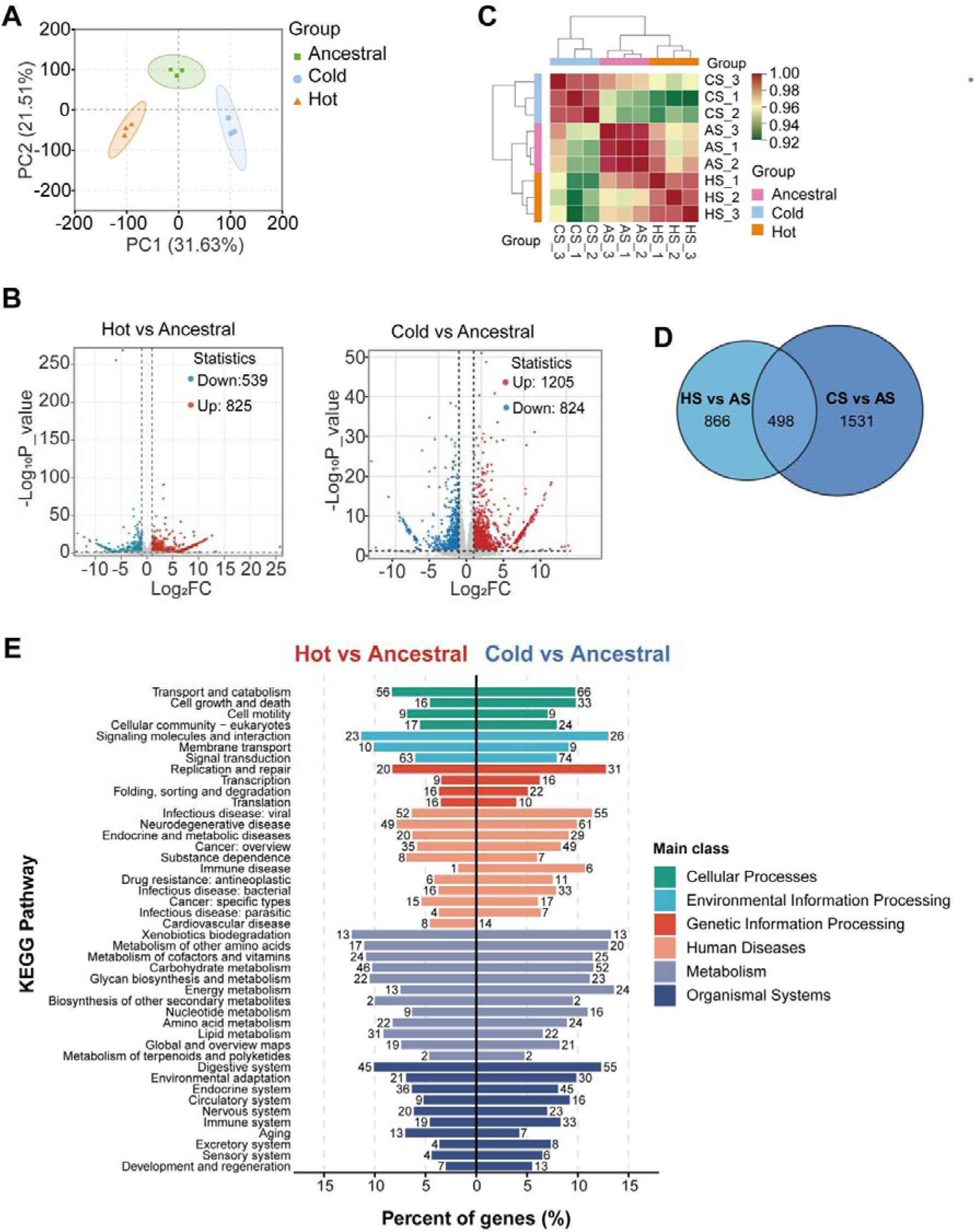
Transcriptomic analysis of the 3^rd^-instar larvae across the ancestral, hot and cold strains. (A) Principal component analysis (PCA) of genes across different strains. PC1 and PC2 represent the first and second principal components, respectively. (B) Volcano plots of differential gene expression, showing significantly up-regulated (red dots) and down-regulated (green dots) genes between HS/CS and AS (FDR < 0.05, fold change > 2). (C) Cluster analysis of the transcriptome. The colors represent the Pearson correlation coefficients between samples, indicating transcriptomic similarity. (D) The number of common or unique differentially expressed genes between HS/CS and AS. (E) KEGG function classification of differentially expressed genes between HS/CS and AS.

The gene expression-based clustering tree using a weighted gene co-expression network analysis (WGCNA) was divided into 29 modules as shown with alphanumeric identifiers (M1-M29) (***Figure 4A; Figure 3–figure supplement 1B***). Module M4 contained the most genes (3463), while module M29 contained the fewest (31) (***Figure 4B***). Selecting the common differential metabolites (30 in total) shared between the hot and cold strains as compared to the ancestral strain, we performed a correlation analysis with co-expressed networks and found that multiple modules, including modules M13, M19, and M24, showed strong correlations with shared differential metabolites. Module M13 was selected for further analysis as it had the highest number of significantly correlated metabolites (28 of 30) (***Figure 4C; Figure 3–figure supplement 1C***). Further analysis revealed that 79 genes within module M13 were differentially expressed in the hot and cold strains when compared with the ancestral strain (***Figure 4D***). These results suggest that genes in module M13 may be candidates involved in the adaptation of *P*. *xylostella* to extreme temperatures.

**Figure 4.**
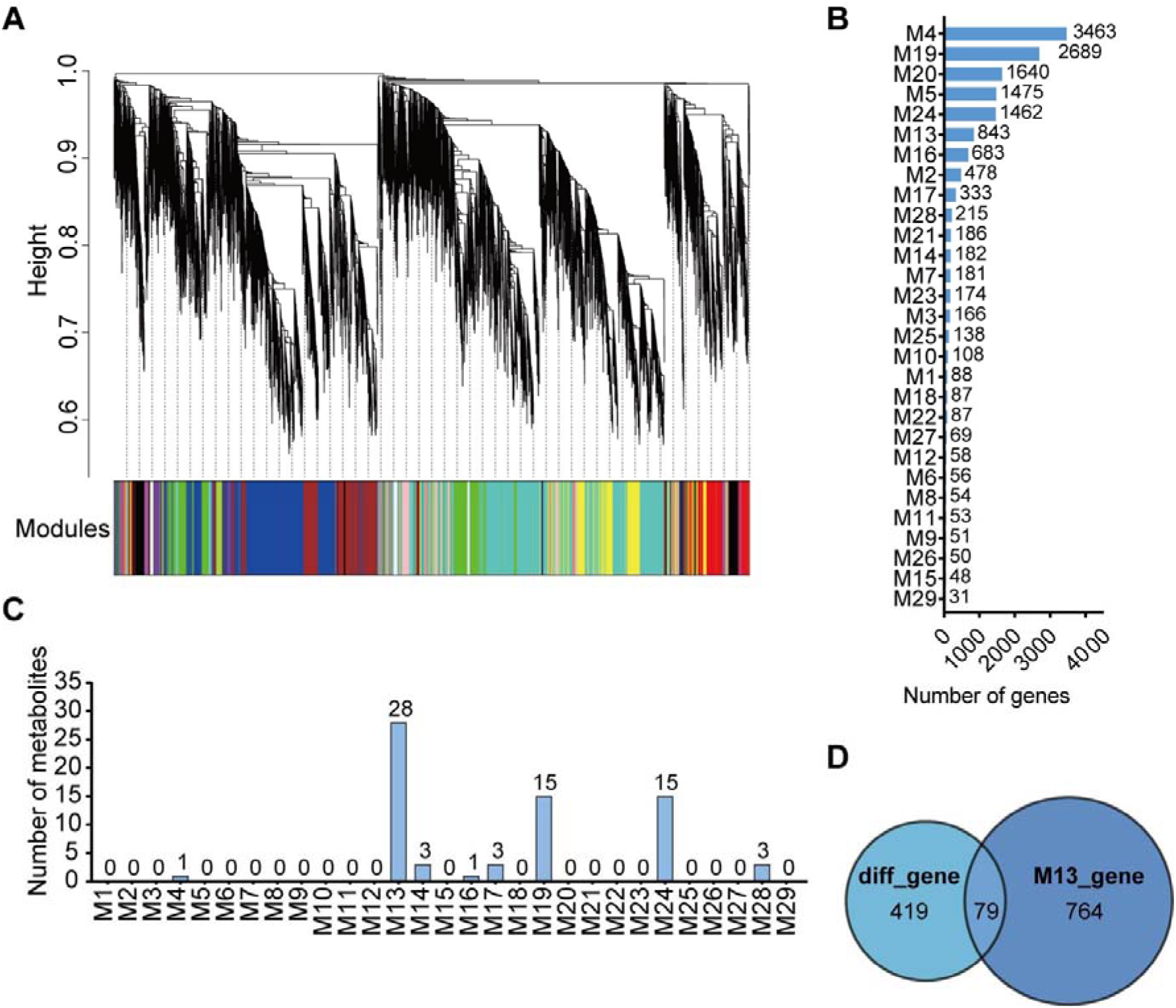
Weighted gene co-expression network analysis (WGCNA) of transcriptomes for the 3^rd^-instar larvae across the ancestral, hot and cold strains (HS, CS and AS) of *P*. *xylostella*. (A) Hierarchical cluster tree illustrating 29 modules identified by WGCNA. (B) Numerical distribution of genes of different modules as identified by WGCNA clustering. (C) In WGCNA, module M13 shows the highest number of metabolites strongly correlated with genes. (D) Overlap of genes in module M13 from WGCNA with common differentially expressed genes between HS/CS and AS.

### Genetic basis of temperature adaptation

To further elucidate the genetic basis of *P*. *xylostella* adaptation to hot and cold environments, from the 79 candidate genes identified above, we selected 15 that were annotated in the genome and had high expression levels (FPKM > 10) for further analysis (***Figure 4D***) and identified 11 genes being successfully amplified. Comparative results revealed eight genes with nonsynonymous mutations and one with a synonymous mutation in both the hot and cold strains (***Supplementary File 2***). Among these genes, we focused on the role of *Px04C00666* (*PxSODC*) in temperature adaptation of *P*. *xylostella* because the deletion of superoxide dismutase (SOD) genes can alter the response of insects to abiotic stresses including temperature (***Bittner et al., 2019; Quan et al., 2024***).

The NCBI database predicted that *PxSODC* contained three exons and two introns, with a conserved domain belonging to the copper-zinc superoxide dismutase superfamily (***Figure 5–figure supplement 1A***) which plays an antioxidative role in cellular defense systems, protecting cells from damage caused by reactive oxygen species (***Fridovich, 1995***). The open reading frame of *PxSODC* was 633 bp and encodes 210 amino acids. Expasy predicted that the molecular weight of the PxSODC protein was 22,168.24 Da, with the isoelectric point being 6.29. According to PSIPRED predictions, its secondary structure consisted of approximately 55.24% random coils, 16.19% alpha helices, and 28.57% extended strands (***Figure 5–figure supplement 1B***). Evolutionary analysis using a Maximum Likelihood approach showed *PxSODC* of *P*. *xylostella* clustered with that of other Lepidoptera insects such as *Operophtera brumata*, *Vanessa cardui*, and *Cydia fagiglandana*, indicating its conserved evolution within this taxonomic order (***Figure 5–figure supplement 1C***). The coding region of *PxSODC* in the hot and cold strains had 23 SNP sites, including 20 synonymous and three non-synonymous mutations (Leu9-Val9, Lys25-Gln25, Leu194-Met194) (***Figure 5A-B***). Leu9-Val9 and Leu194-Met194 mutations were involved in the substitution of hydrophobic amino acids. Based on sequencing of 10 individuals per strain, the Leu194-Met194 mutation was present at a frequency of 70% in HS, 90% in CS, and 30% in AS (***Figure 5B***). The expression of *PxSODC* at different developmental stages of the hot and cold strains was significantly lower than that of the ancestral strain (***Figure 5–figure supplement 2A***). After 2 h exposure of the 3^rd^-instar larvae to the stress of high (32°, 34°, 36°, 38° and 40°C) or low (12°, 10°, 8°, 6° and 4°C) temperature environments, the expression of *PxSODC* in the hot and cold strains was significantly lower than in the ancestral strain (***Figure 5–figure supplement 2B-C***). This suggests that the expression of the *PxSODC* gene is regulated by temperature triggers, and its altered function contributes to the temperature-adaptive evolution in *P*. *xylostella*.

**Figure 5.**
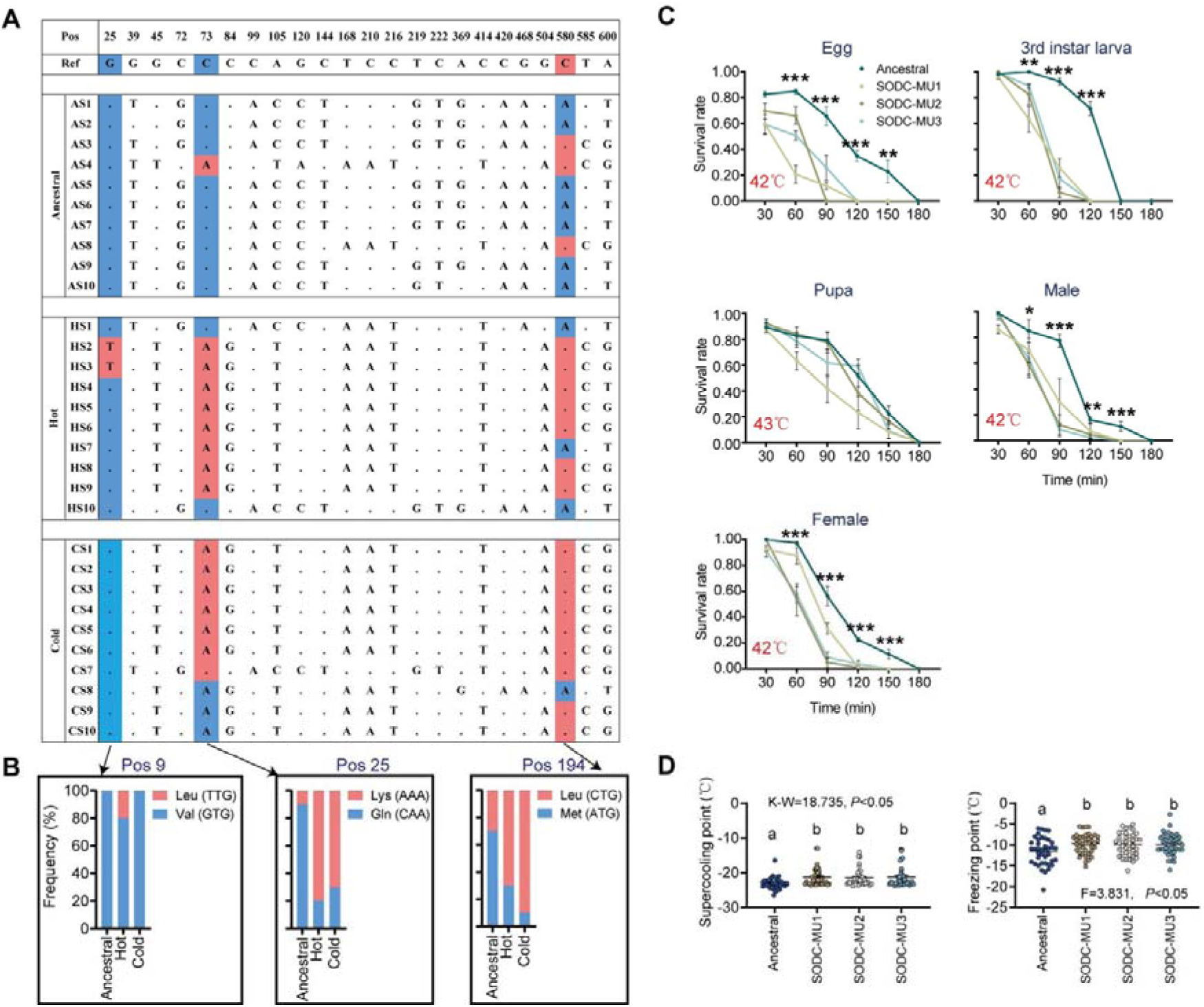
Role of *PxSODC* in temperature adaptation of *P*. *xylostella*. (A) Allele frequencies of SNPs in the *PxSODC* gene amplified by PCR from the ancestral, hot and cold strains (AS, HS and CS). The analysis involves ten 4^th^-instar larvae from each of the strains; the dot (·) indicates identity with the reference base. (B) Frequency of amino acid translations from non-synonymous codon mutations in the *PxSODC* gene in different strains. (C) Stage-specific survival rates of the ancestral and mutant strains (AS, SODC-MU1, SODC-MU2 and SODC-MU3) under extreme heat conditions. (D) Supercooling and freezing points of the pupae from different strains (AS, SODC-MU1, SODC-MU2 and SODC-MU3). Data are presented as mean±SEM, one-way ANOVA with Tukey’s test was used for comparison.Six biologically independent samples were used in (C) and significant levels between groups with the same stress duration are indicated by asterisks (\**p* < 0.05, \*\**p* < 0.01, \*\*\**p* < 0.001). A total of 40 biologically independent samples were used in (D) and statistical significance is indicated by different letters (*p* < 0.05). **Source data 1.** Raw data for thermal tolerance phenotypes of *PxSODC* knockout strains.

To elucidate the structural mechanism by which these non-synonymous mutations enhance protein function under thermal stress, we performed 100 ns MD simulations on AlphaFold-generated models of the wild-type (WT) and mutant (MU) PxSODC at 15°C, 26°C (favorable baseline), and 32°C (***Figure 5–figure supplement 3***). RMSD analysis revealed that at the 26°C baseline, both WT and MU exhibited comparable structural stability (1.62 ± 0.21 vs. 1.59 ± 0.27) (***Figure 5–figure supplement 3B***). However, under heat stress (32°C), WT underwent severe conformational drift (RMSD surged to 2.49 ± 0.35, an increase of 0.87 from baseline), while MU remained remarkably stable (1.66 ± 0.26, an increase of only 0.07) (***Figure 5–figure supplement 3C***). SASA analysis showed that MU possessed an inherently more compact structure than WT, with lower values at both 15°C (118.39 ± 7.57 vs. 127.29 ± 6.12 nm^2^) and 26°C (113.82 ± 7.40 vs. 125.61 ± 6.76 nm^2^), indicating optimized hydrophobic core packing (***Figure 5–figure supplement 3D***). Analysis of intramolecular hydrogen bonds further revealed that the MU network exhibited dual stress resistance: under cold stress (15°C), MU actively increased hydrogen bonds from its 26°C baseline (113→119), demonstrating a compensatory response, while WT suffered bond loss (117→112); under heat stress (32°C), MU fully maintained its hydrogen bond count (113→113), whereas WT showed a slight decrease (117→116) (***Figure 5–figure supplement 3E***). Collectively, these simulations demonstrate that the non-synonymous mutations confer enhanced global structural rigidity and dual-directional thermal resilience to PxSODC through a more compact hydrophobic core and a more resilient intramolecular hydrogen bond network, providing a direct structural basis for its increased catalytic efficiency at lower expression levels.

The above analyses revealed naturally occurring non-synonymous mutations in *PxSODC* that are enriched in the hot and cold strains. To directly test whether *PxSODC* is functionally required for thermal adaptation, we generated loss-of-function mutants by disrupting *PxSODC* in the ancestral strain using CRISPR/Cas9-mediated mutagenesis. Of 162 eggs treated with CRISPR/Cas9, 75 successfully developed into adults. We confirmed three mutant strains in the G0 generation of *P*. *xylostella*: +1 bp (SODC-MU1), +2 bp (SODC-MU2) and -1 bp (SODC-MU3). Self-crossing continued, and three types of homozygous mutation were obtained in the G5 generation (***Figure 5–figure supplement 4***). Life table analysis showed that the three SODC-MU strains had prolonged development, lower survival rates, and reduced fecundity and population fitness compared to the ancestral strain, particularly under hot/cold environments (***Appendix 2***; ***Figure 5–figure supplement 5*; *Supplementary File 3-5***).

We then assessed the stage-specific survival of the ancestral strain and all three SODC-MU mutant strains (SODC-MU1, SODC-MU2 and SODC-MU3) under extreme heat stress to determine whether *PxSODC* loss consistently impairs thermal tolerance. At 42°C, the survival rates of eggs, 3^rd^-instar larvae, female adults and male adults of the mutant strains were significantly lower from those of the ancestral strain at several time points (***Figure 5C***). However, survival rates of the mutant pupae exposed to the high temperature (43°C) were not significantly different from those of the ancestral strain at different time points (***Figure 5C***). The pupal stage appeared more tolerant to high temperature than other life stages, as the *PxSODC* knockout did not significantly reduce pupal survival at 43°C while it significantly reduced survival of eggs, larvae, and adults at 42°C (***Figure 5C***). This may be due to pupal-specific heat resistance mechanisms, such as protective chrysalis and U-shaped metabolism (***Kaiser et al., 2010; Barros-Cordeiro et al., 2014***). In addition, supercooling and freezing points of the mutant strains (MU1: -21.32±0.41 and -9.75±0.38; MU2: -21.50±0.38 and -9.93±0.43; MU3: -21.23±0.48 and -9.94±0.41) were significantly higher than those of the ancestral strain (-23.09±0.26 and -11.58±0.52) at pupal stage (***Figure 5D***), indicating a key role of the *PxSODC* gene in the adaptability and tolerance of *P*. *xylostella* to extreme temperatures.

To further investigate the effect of *PxSODC* gene mutations on the temperature adaptability of *P*. *xylostella*, we identified five genes from the same SOD family in transcriptomes of the 3^rd^-instar larvae from the three tested strains. We found that *Px04C00505* and *Px13C00423* showed SNP mutations in the hot and cold strains, whereas *Px20C00248*, *Px15C00224* and *Px15C00223* were not mutated (***Supplementary File 6***). Further comparison of gene expressions across different strains revealed that, relative to the ancestral strain, the expression levels of *PxSODC*, *Px04C00505*, and *Px13C00423* were significantly reduced in the hot and cold strains, while the remaining genes maintained stable or increased expression levels (***Figure 6A***). Concurrently, SOD activity decreased in the hot and cold strains, along with a reduction in O_2_^-^ levels (***Figure 6B-C***). When SOD expression and activity, as well as O_2_^-^ levels, were compared under different temperature conditions between the ancestral and hot/cold strains, similar patterns were observed (***Figure 6D-F***). These results suggest that non-synonymous mutations in the hot and cold strains may alter SOD protein conformation, increasing catalytic efficiency per molecule and enabling effective O_2_ ^-^ scavenging at lower expression levels. This energy-efficient strategy is beneficial under thermal stress, where conserving metabolic resources for development and reproduction is critical for survival. Compared to the ancestral strain, expression of the mutated *SODC* genes (*Px04C00505* and *Px13C00423*) was increased in the male adults in SODC-MU1 and SODC-MU2. The expression levels of the non-mutated genes *Px20C00248*, *Px15C00224* and *Px15C00223* were also increased (***Figure 6G***). Further, SOD activity decreased, while O_2_^-^levels increased in the two mutant strains (***Figure 6H-I***), which were unable to fully compensate for the effects caused by the deletion of the *PxSODC* gene, implying that the SOD protein encoded by *PxSODC* plays a crucial role in O_2_^-^ scavenging.

**Figure 6.**
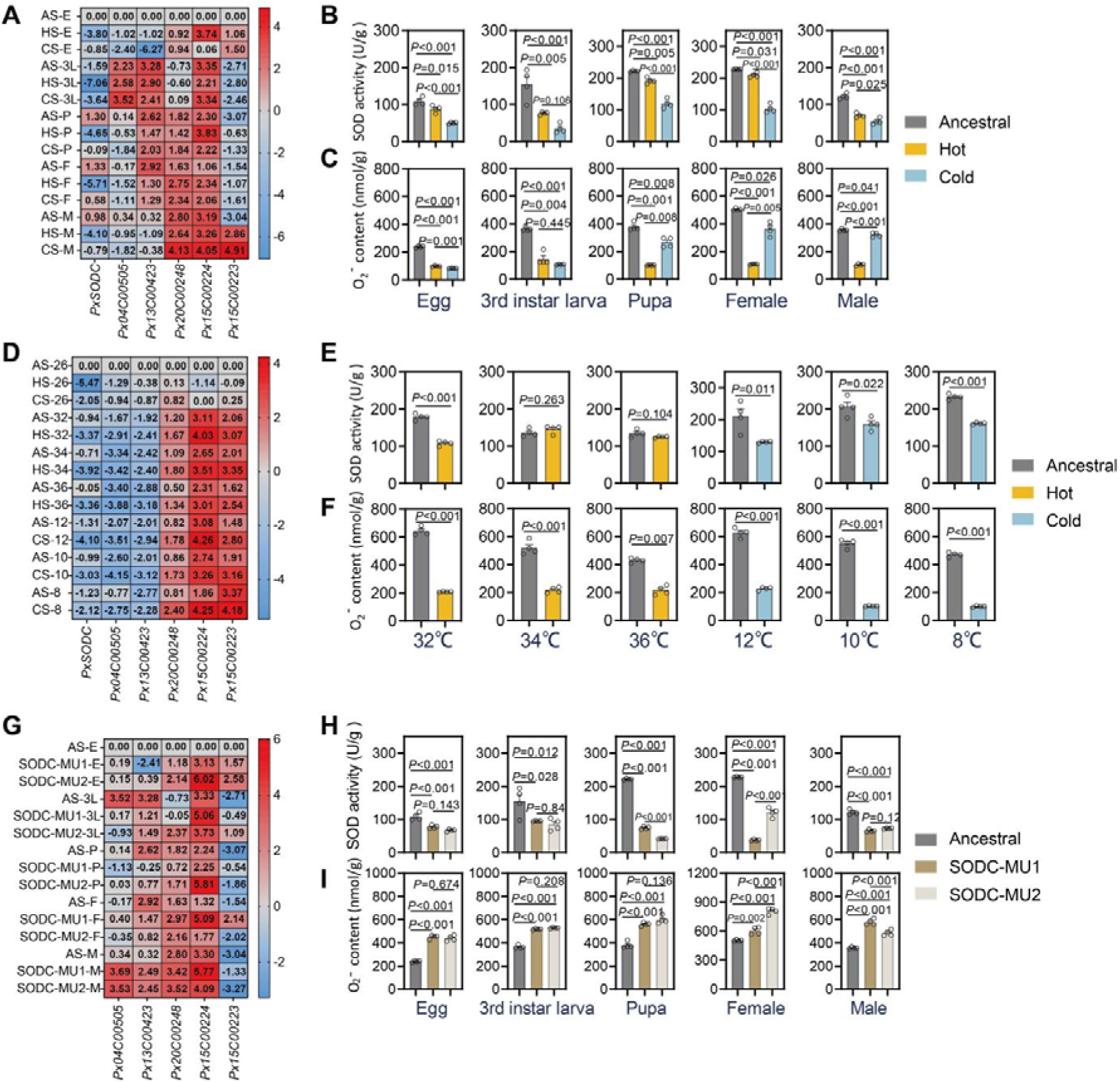
SOD expression and activity and superoxide anion (O_2_^-^) levels across developmental stages and temperature environments in different strains of *P*. *xylostella*. (A) Expression levels of the *SOD* genes at different developmental stages of AS, HS, and CS in the favorable temperature environment (26°C). (B) SOD enzyme activity at different developmental stages of the ancestral, hot and cold strains (AS, HS, and CS) in the favorable temperature environment. (C) O_2_ ^-^ levels at different developmental stages of AS, HS, and CS in the favorable temperature environment. (D) Expression levels of the genes from the *SOD* family in the 3^rd^-instar larvae of AS, HS, and CS in the hot (32°C, 34°C, 36°C) and cold (12°C, 10°C, 8°C) environments. (E) SOD enzyme activity in the 3^rd^-instar larvae of AS, HS, and CS in the extreme temperature environments. (F) O_2_^-^ levels in the 3^rd^-instar larvae of AS, HS, and CS in the hot and cold environments. (G) Expression levels of *SOD* family (excluding the *PxSODC* gene) at different developmental stages of the ancestral and SODC-MU strains in the favorable temperature environment. (H) SOD enzyme activity at different developmental stages of the ancestral and SODC-MU strains in the favorable temperature environment. (I) O_2_ ^-^ levels at different developmental stages of the ancestral and SODC-MU strains in the favorable temperature environment. n = 3 biologically independent samples in (A), (D), (G); within each of the boxes, the numerical value represents log_2_-fold change of the gene expression level in the treated samples with respect to the control. n = 4 biologically independent samples in (B), (C), (E), (F), (H), with data being presented as mean±SEM. One-way ANOVA with Tukey’s test was used for comparison in (A), (B), (C) and (G), (H), (I) (*p* < 0.05). t-test was used for comparison in (D), (E), (F) (*p* < 0.05). **Source data 1.** Raw data for SOD enzyme activity and superoxide anion (OCC) levels across strains, developmental stages, and temperature conditions.

### *PxSODC*-allied metabolic networks

Untargeted metabolomic analysis of the ancestral and SODC-MU strains across developmental stages revealed broad metabolic adjustments involving lipids, nucleotides, carbohydrates, and amino acids following *PxSODC* deletion (***Appendix 3***; ***Figure 7-figure supplement 1 and 2***). In the metabolome, the abundance of three metabolites, namely 5-hydroxymethyluracil, 2-methylcitric acid, and 5’-deoxyadenosine, differed between the mutant strains (SODC-MU1 and SODC-MU2) and the ancestral strain across all developmental stages (***Figure 7A***). With the exception of 5’-deoxyadenosine, which was significantly higher in the 3^rd^-instar larvae, these three metabolites were significantly lower in other developmental stages of the mutant strains (***Figure 7B***). This suggested a possible direct association with *PxSODC*, and may represent a key biological regulatory response in *P. xylostella*’s adaptation to different environmental conditions. Importantly, this specific metabolic signature provided a compelling, data-driven hypothesis linking *PxSODC*-mediated oxidative stress management directly to epigenetic regulation: 5-hydroxymethyluracil is directly involved in dynamic DNA demethylation, and 5’-deoxyadenosine is a precursor to S-adenosylmethionine, the primary methyl donor for DNA methylation (***Bhutani et al., 2011; McKean et al., 2019***). Their consistent alteration in the SODC-MU strains suggested a potential link between *PxSODC* and DNA methylation. To test this hypothesis, we measured the expression levels and enzymatic activities of DNA methyltransferase 1 (*PxDnmt1*) in the 3^rd^-instar larvae of different strains. The results showed that both the expression and enzymatic activity of *PxDnmt1* were significantly reduced in the hot and cold strains compared to the ancestral strain (***Figure 7C-D***). Using RNA interference (RNAi) technology, we specifically silenced the expression of *PxDnmt1* in the ancestral strain of *P. xylostella* (***Figure 7E***) and observed significantly reduced levels of 5-methylcytosine (5-mC, a marker of DNA methylation) in both pupae and female adults (***Figure 7F***) (***Ni et al., 2023***). Further, we found that silencing of *PxDnmt1* could significantly decrease the critical thermal maximum (CTmax) of female adults and increase the supercooling and freezing points of pupae (***Figure 7G-H***). These results suggest that DNA methylation may play a role in the thermal tolerance of *P. xylostella*, though further work is needed to establish a direct mechanistic link between *PxSODC*, methylation-related metabolites, and epigenetic regulation of thermal adaptation.

**Figure 7.**
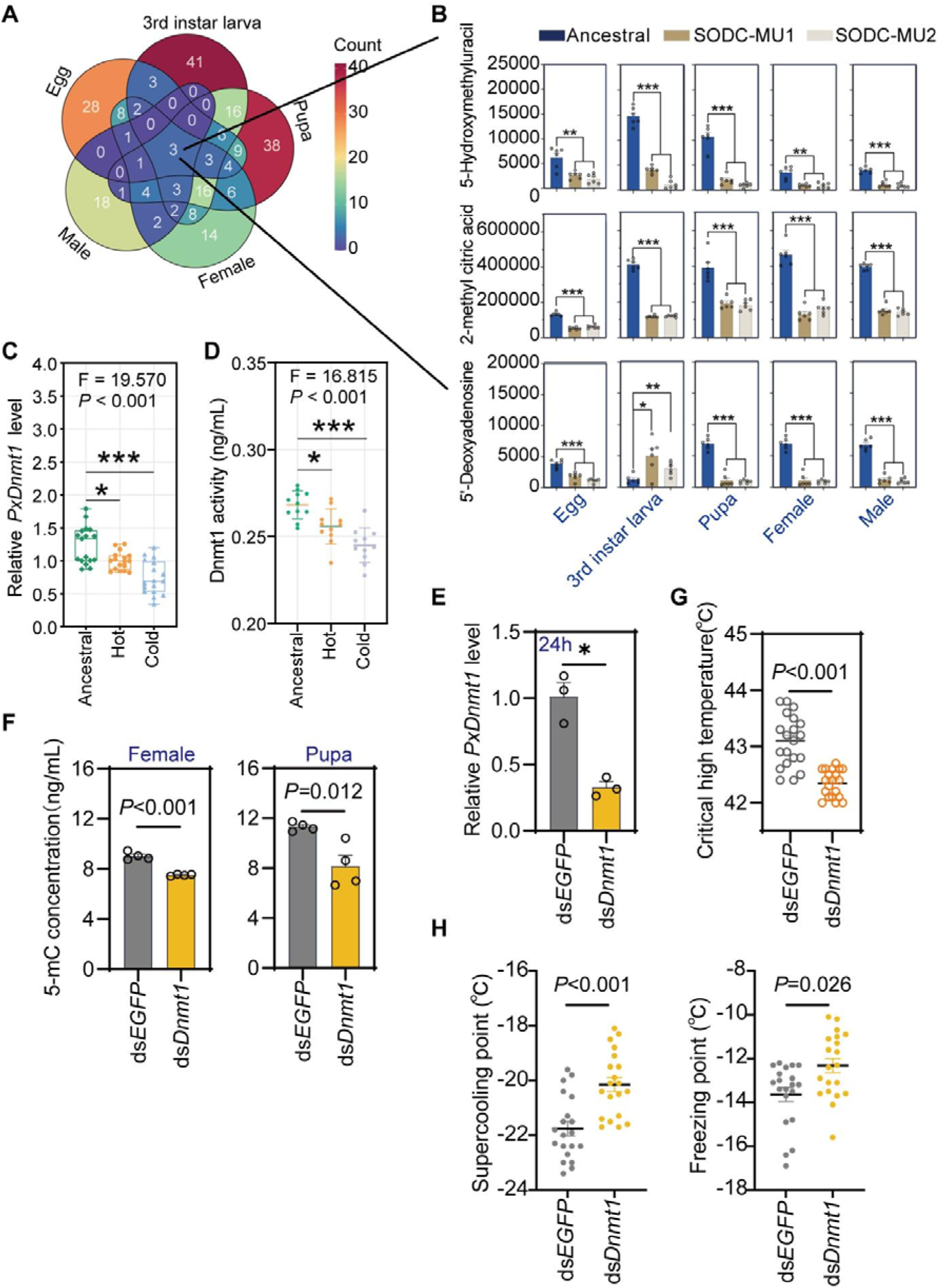
Comparison of metabolites and DNA methylation across different strains of *P*. *xylostella*. (A) A Venn diagram showing the metabolites that are consistently different between the ancestral and mutant strains across different developmental stages. (B) Three metabolites with persistent discrepancy across different developmental stages in the ancestral and mutant and strains. (C) Expression level of the DNA methyltransferase 1 gene (*PxDnmt1*) in the ancestral, hot and cold strains. *n* = 17 biologically independent samples. (D) DNA methyltransferase activity in the ancestral, hot and cold strains. *n* = 12 biologically independent samples. Data are presented as mean±SEM, one-way ANOVA with Tukey’s test was used for comparison in (C), (D) (*p* < 0.05). (E) Injection of ds*Dnmt1* significantly reduced the expression level of *PxDnmt1* in the ancestral strain of *P. xylostella*. *n* = 3 biologically independent samples. (F) Silencing of *PxDnmt1* decreased 5-methylcytosine (5-mC) content in the female adults and pupae of *P. xylostella*. *n* = 4 biologically independent samples. (G) Female adults with silenced *PxDnmt1* exhibited a significantly decreased critical thermal maximum (CTMax). *n* = 20 biologically independent samples. (H) Pupae with silenced *PxDnmt1* displayed elevated supercooling and freezing points. *n* = 40 biologically independent samples. Data are presented as mean±SEM, unpaired t-test was used for comparison in (E), (F), (G), (H) (*p* < 0.05). **Source data 1.** Raw data for *PxDnmt1* expression, DNA methyltransferase activity, and effects of *PxDnmt1* silencing on thermal tolerance.

## Discussion

Insects are valuable bio-indicators of the effects of climate change via their phenology, distribution, population dynamics responses (***Bale et al., 2002; Halsch et al., 2021***). The present study demonstrates that *P. xylostella* can undergo rapid genetic adaptation to thermal extremes, providing mechanistic insights into how this globally invasive pest may expand its range under climate change through coordinated transcriptomic, metabolomic, and life history changes. By conducting the life history trait, demography, and fitness assessment of the ancestral, hot and cold strains of *P. xylostella*, we observed that the hot and cold strains had evolved significant genetic differences from the ancestral strain in multiple traits related to thermal tolerance and fitness. In the context of global warming, *P*. *xylostella* may evolve greater flexibility across its range and ecological niche leading the hot-evolved populations to be able to persist in regions with increased temperatures due to climate change (***Chen et al., 2021***). With climate change, cold-adapted *P. xylostella* may be favored during episodes of late frost in the spring or early frost in the autumn in temperate regions. The lower supercooling and freezing points of the cold strain pupae facilitate the survival of insects in cold climates, extending their ecological adaptability to low-temperature environments (***Block, 1997***). The demonstrated capacity of *P. xylostella* to adapt to extreme thermal conditions of both forms implies that *P*. *xylostella* may survive under a broader range of climatic conditions, posing new challenges for the management and control of this worldwide pest.

Our findings reveal a significant metabolic adjustment in *P*. *xylostella* during its adaptive evolution to both high and low temperatures. The convergent reduction of lipid-related metabolites such as octadecenoic acid, epoxystearic acid, and carnitine in both the hot and cold strains (***Figure 2F***) suggests a shared metabolic adjustment during thermal adaptation. Lipids play a key role in energy storage and membrane stability in insects, and downregulation of lipid metabolism may enable *P. xylostella* to conserve metabolic resources and reallocate energy toward development and reproduction (***Rommelaere et al., 2019***; ***Mallard et al., 2018***). As a key mediator of fatty acid transport and energy metabolism, the reduction in carnitine levels may further reflect this energy reallocation strategy (***Bremer, 1983***).

Although gene expression between the hot and cold strains does not show a strong correlation, KEGG analysis indicates that differentially expressed genes in both adapted strains share similarly enriched core pathways (***Figure 3E***). Notably, the convergent reduction in lipid metabolism represents a shared energy reallocation strategy. Freeing up resources to fuel the accelerated development, higher fecundity, and extreme-heat survival in the hot strain, while facilitating the extended male longevity and lower supercooling and freezing points required in the cold strain (***Tigano et al., 2020***; ***Sherpa et al., 2022***; ***Li et al., 2024a***). Relative to the ancestral strain, the cold strain exhibited more differentially expressed genes (2029 *vs.* 1364) but fewer differential metabolites (37 *vs.* 77) than the hot strain. This pattern contrasts with findings in *Drosophila*, where thermal adaptation produced a simpler transcriptomic (***Mallard et al., 2018***). This divergence in omics profiles strictly aligns with their distinct physiological requirements. The cold-adapted strain relies on broader transcriptional reprogramming to structurally maintain basic cellular homeostasis and sustain cold hardiness over its prolonged lifespan, whereas the hot-adapted strain utilizes targeted gene expression changes combined with broader metabolic rewiring to actively sustain rapid energy turnover and its accelerated life cycle.

Mutations in these genes during adaptive evolution in response of thermal adaptation can lead to novel phenotypic traits, such as changes in the life history and population fitness of insects (***Gibson et al., 2019***). The *PxSODC* gene encodes a superoxide dismutase (SOD) that scavenges superoxide anions in cells, maintaining redox balance and protecting cellular structures (***Sheng et al., 2014***). Under extreme temperatures, insects can adjust their survival strategy, allocating more energy to maintain fundamental life functions (***Hoffmann and Sgro, 2011***). Here, the deletion of the *PxSODC* gene led to reduced SOD enzyme activity and increased O_2_^-^ levels, reducing the tolerance of *P*. *xylostella* to extreme temperatures. These findings provide novel insights into how genetic variation translates into phenotypic variation (***You et al., 2024***), and the ways in which *P*. *xylostella* responds – and has responded – to a changing climate (***Chen et al., 2021***).

Our molecular dynamics simulations provide direct structural evidence that the non-synonymous mutations enhance *PxSODC* protein thermostability (***Figure 5–figure supplement 3***). Specifically, the mutant protein maintained structural integrity under both heat and cold stress through a more compact hydrophobic core and a more resilient hydrogen bond network. This enhanced dual-directional thermostability likely increases catalytic efficiency per molecule, enabling effective superoxide scavenging at the lower expression levels observed in the evolved strains and reducing the energetic cost of enzyme production, consistent with previous findings that specific structural mutations can directly increase SOD enzyme activity (***Sheng et al., 2013***). After long-term adaptation, insects may acquire the ability to maintain cellular homeostasis in new thermal environments by reducing basal metabolism and allocating more energy to development and reproduction (***Mallard et al., 2018***). At 34°C and 36°C, the trends in SOD enzyme activity and *PxSODC* gene expression differ between different temperature-evolved strains, suggesting the involvement of additional genes in the regulation of SOD enzyme activity. This hypothesis is supported by our transcriptomic analysis which identified additional *SOD* genes (***Supplementary File 5***).

In the SODC gene family, two *SOD* genes (*Px04C00505* and *Px13C00423*) underwent non-synonymous mutations and showed reduced expression in the hot and cold strains at different developmental stages, while the expression of three *SOD* genes without mutations remained relatively stable or increased at most developmental stages. The evolution of protein functions is driven by mutations, which in some cases can switch directly from one function to another through single amino acid changes (***Nobeli et al., 2009; Patsch et al., 2024***). Under extreme temperatures, the expression of these genes in the 3^rd^-instar *P*. *xylostella* larvae of the hot and cold strains trends to be similar to those of the ancestral strain. This indicates that adverse environmental conditions increase intracellular oxidative stress, which requires regulation of SOD expression and enzyme activity to scavenge superoxide anions (***Islam et al., 2022***). Maintaining high levels of SOD enzyme activity requires additional energy, placing a strain on cellular energy metabolism and resource allocation (***Emre et al., 2013***). *SOD* genes with non-synonymous mutations, like *PxSODC*, can lead to the change in protein structure or function, affecting enzyme activity and allowing for faster O_2_^-^ scavenging at lower transcript levels, reducing resource requirements. The three unmutated *SOD* genes, if mutated, might adversely affect the moth or have other functions, such as involvement in cellular signaling pathways (***Mondola et al., 2016***).

Metabolomic analysis of different developmental stages in the ancestral and mutant strains revealed that after the loss of *PxSODC* gene, the metabolism of *P*. *xylostella* underwent temperature-adaptive adjustments involving lipids, nucleotides, carbohydrates, coenzymes and vitamins, and amino acids. This study also showed that DNA methylation plays a key role in the temperature adaptation of *P*. *xylostella*. While DNA methylation may be associated with gene activation, its main function remains the inhibition of gene expression (***Stroud et al., 2015; Wang et al., 2018***). Transcriptomic analysis also showed that more genes were up-regulated in the hot and cold strains compared to the ancestral strain, highlighting the role of DNA methylation in regulating gene expression to help *P*. *xylostella* maintain physiological functions and survive. In addition to directly regulating the expression of temperature responsive genes, epigenetic effects can also indirectly affect the response of insects to temperature challenges (***Reynolds, 2017***). By adding the molecular data of epigenetic markers, underlying mechanisms on the adaptive adaptation can be more easily elucidated. Therefore, further work is required to better understand the impact of non-genetic effects on adaptation to future climates including how they interact with genetic adaptive capacity. While our experimental evolution approach successfully uncovered potential genetic mechanisms of thermal adaptation, its ecological relevance warrants consideration. Exposing the insects to extreme 12 h thermal bounds was necessary to impose strong directional selection and observe adaptive evolution over a relatively short timeframe (∼3 years) in the laboratory. However, we acknowledge that comparing fluctuating stressful environments to a constant favorable environment (26°C) may conflate adaptation to absolute temperature extremes with adaptation to thermal fluctuation itself. Future field-based studies across natural gradients, or using fluctuating optimal control regimes, will be valuable to validate the ecological impact of these mutational and epigenetic pathways in wild habitats.

This study elucidates the molecular mechanisms underlying adaptation of *Plutella xylostella* to both high- and low-temperature environments and functionally validates differentially expressed genes identified in ancestral, hot-evolved, and cold-evolved strains. Nevertheless, thermal adaptation in arthropods may engage distinct, temperature-specific biological pathways; accordingly, future work will prioritize the characterization of strain-specific differentially expressed genes. Beyond functional validation of the canonical stress-associated gene *PxSODC*, additional genes harboring nonsynonymous mutations warrant detailed investigation to clarify their roles within the broader regulatory network. Importantly, our findings also underscore a critical role for DNA methylation in thermal adaptation in *P. xylostella*. More broadly, future research should integrate genetic, epigenetic, and metabolic approaches across diverse taxa to determine whether adaptive mechanisms such as SOD-mediated oxidative stress regulation and DNA methylation represent general strategies by which arthropod pests adapt to novel thermal environments and expand their ranges under climate change.

## Materials and methods

### Key resources table

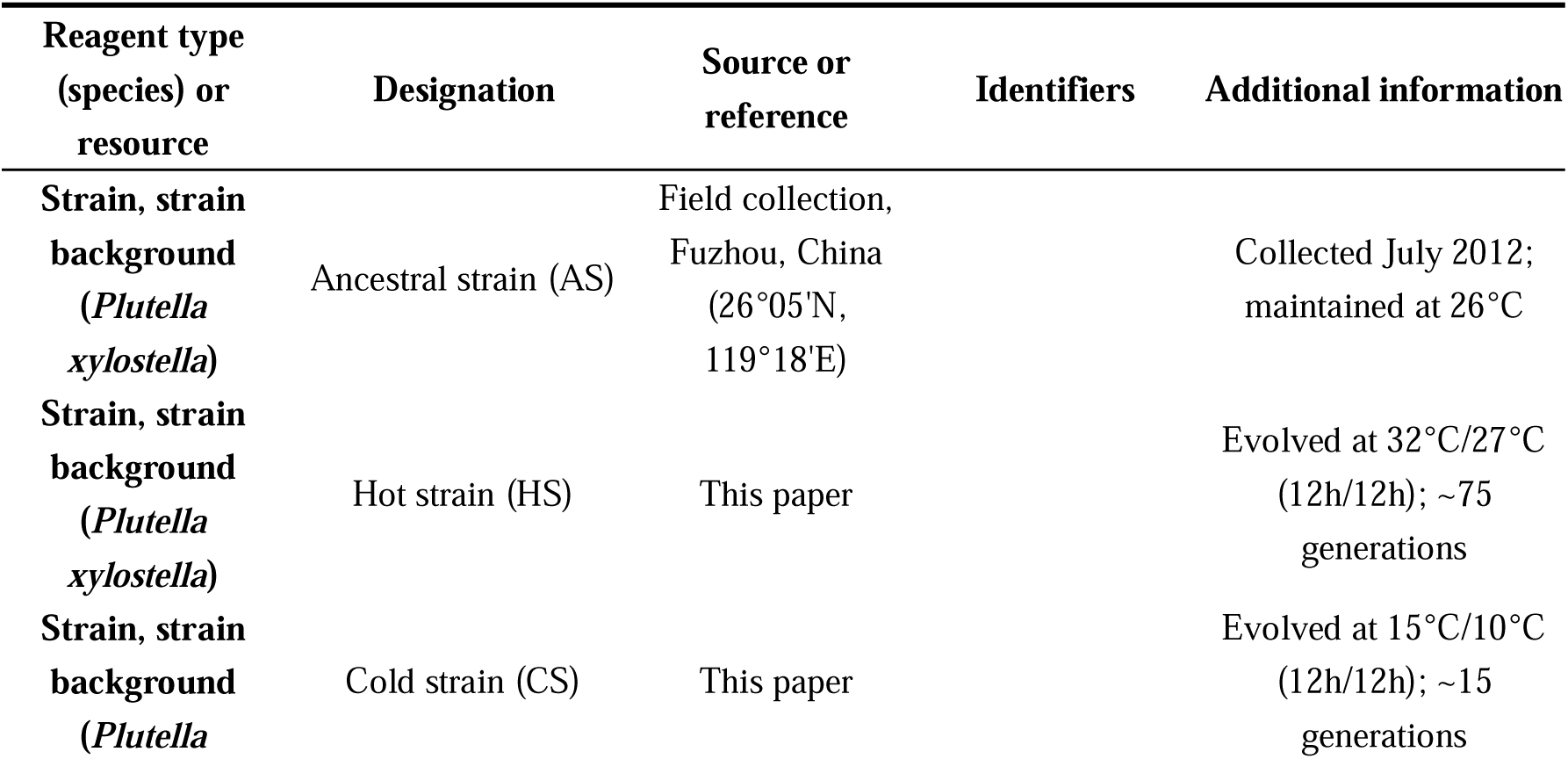
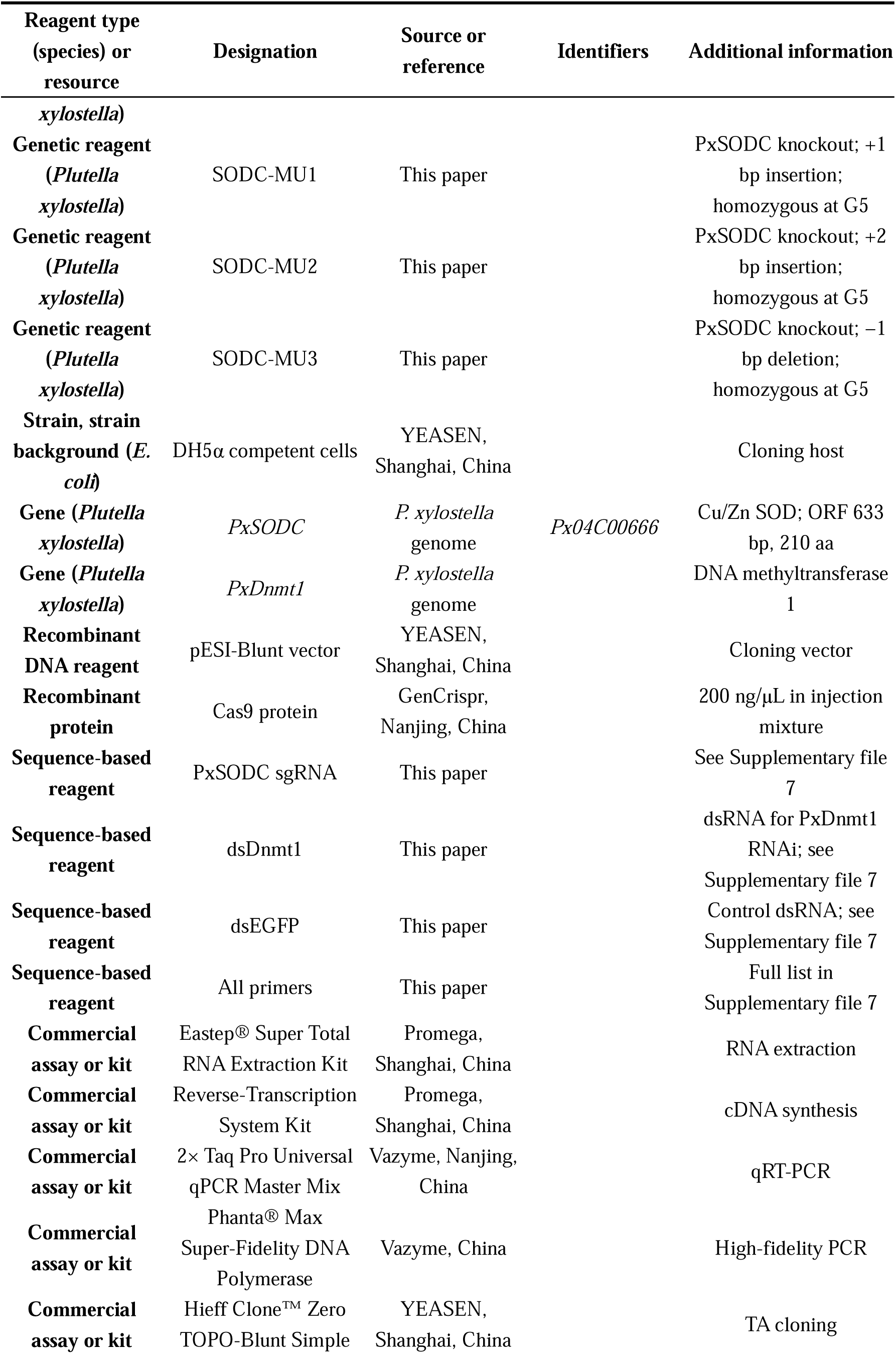
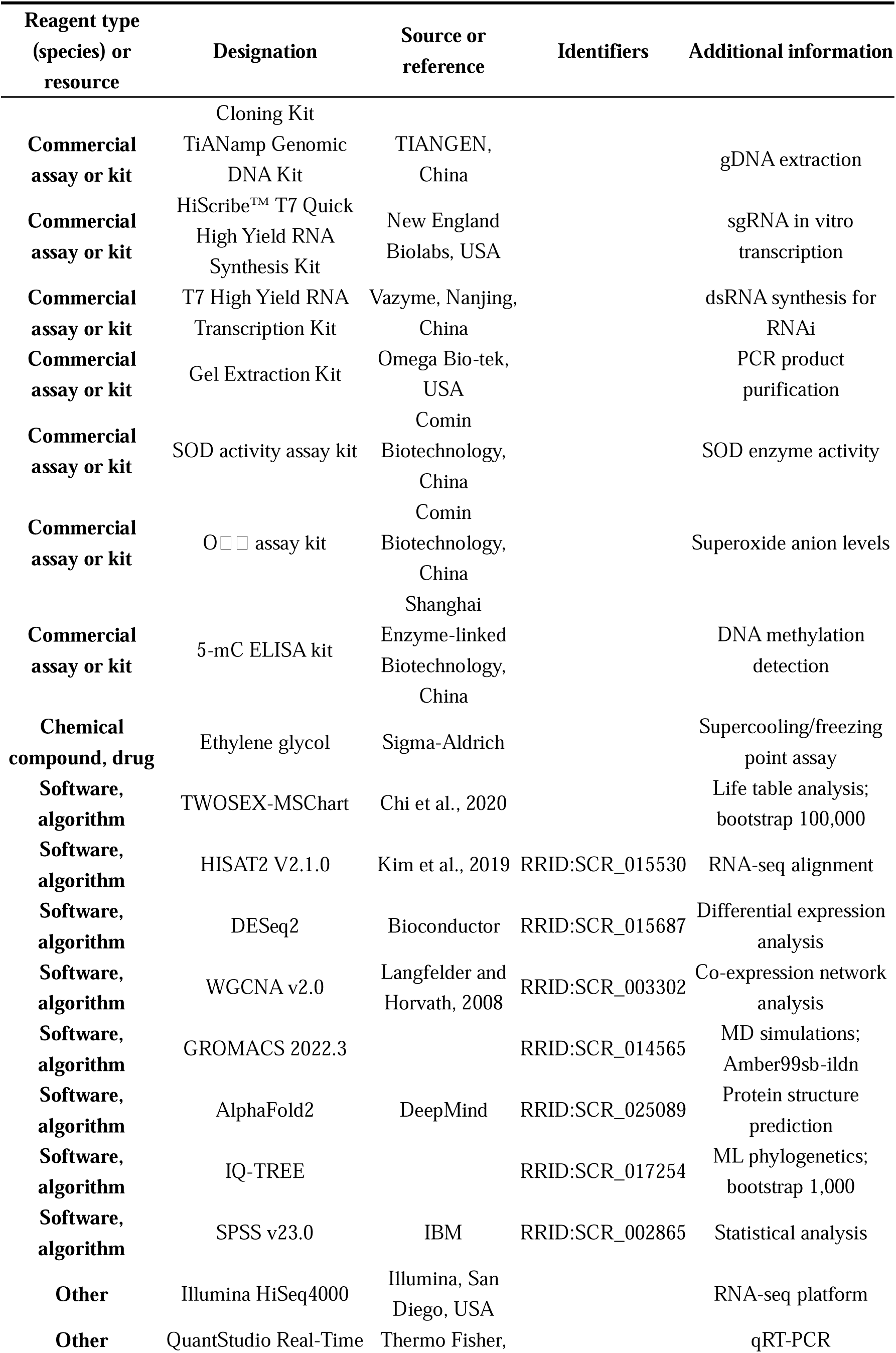
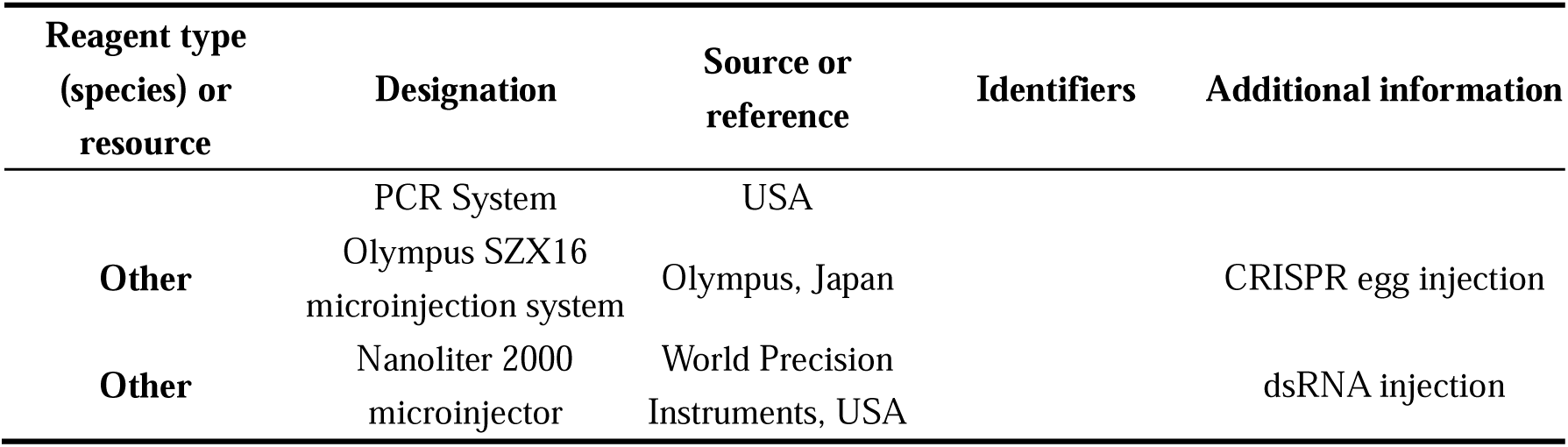

### Insects

The founding population of *P. xylostella* was established from field-collected larvae on cabbage (*Brassica oleracea* var. *capitata*) in organic farms in Fuzhou, Fujian Province, China (26°05’N, 119°18’E) in July 2012. Fuzhou experiences a subtropical monsoon climate, where summer high temperatures exceed 32°C and winter lows can drop below 10°C, making our experimental selection temperatures ecologically relevant extremes for this population. The collection site was confirmed through farmer interviews and local agricultural records to have no history of insecticide application for at least five years. Approximately 300-500 larvae were collected from multiple host plants distributed across a 2-hectare area to maximize genetic diversity and minimize founder effects. The field-collected population was reared in the laboratory under controlled conditions of 26°C and 60% relative humidity with a 12-h light/12-h dark cycle, without exposure to insecticides. This setup was referred to as the favorable temperature environment based on a previous study on the relationship between temperature and the *P. xylostella* development (***Liu et al., 2002***). Population size was maintained at >500 individuals per generation to minimize inbreeding and genetic drift. This laboratory-acclimated population was designated as the ancestral strain and served as the baseline for all subsequent experiments. Eggs and larvae were reared in sterile plastic disposable Petri dishes (90 mm) with an artificial diet containing 68 g yeast powder, 20.4 g agar, 127.5 g raw wheat germ, 3.4 g potassium sorbate, 3.4 g methyl paraben, 34 g sucrose, 10.2 g powder of radish seed, 1.7 g vitamin premix, 3.4 g ascorbic acid, 3.4 mL cola oil, 0.34 mL linoleic acid and 850 mL water The adults were allowed to mate and lay eggs in disposable paper cups, and were fed with a 10% honey solution.

The previous study on the relationship between temperature and developmental rate shows that *P*. *xylostella* can survive and develop at the temperatures 32°C at the maximum, 26°C as the optimum, and 10°C for the minimum (***Liu et al., 2002***). To generate thermally adapted populations, we established 18 replicate populations from the ancestral strain and randomly assigned them to three thermal regimes: (1) a hot-evolved treatment with cycling temperatures of 32°C/27°C (12 h light/12 h dark), (2) a cold-evolved treatment with cycling temperatures of 15°C/10°C (12 h light/12 h dark), and (3) a control treatment maintained at constant 26°C. The hot and cold regimes used cycling temperatures to simulate diurnal fluctuations experienced in natural environments, while the control was kept at the constant optimal developmental temperature for *P. xylostella* (***Liu et al., 2002***). All other environmental conditions (humidity, photoperiod, diet) remained identical across treatments. The populations were maintained with non-overlapping generations and a census population size of approximately 500 individuals per replicate.

Prior to the start of the selection experiment, the ancestral population was maintained in the laboratory at 26°C for approximately ∼170 generations. The thermal selection experiment was then conducted for approximately three years, encompassing ∼75 and ∼15 generations for the hot and cold strains, respectively. To minimize maternal effects and ensure that observed differences were due to genetic adaptation rather than developmental plasticity, all populations were reared for two additional generations under common garden conditions (26°C) prior to phenotypic and molecular assays. All six replicate populations per treatment were used for downstream experiments, resulting in 18 experimental cohorts in total.

### Development of age-stage-specific two-sex life tables

To study the life history and population fitness of the three different *P*. *xylostella* strains, age-stage-specific life tables were developed. A total of 90 randomly selected newly laid eggs from each strain were transferred to three 90 mm diameter Petri dishes (30 eggs per dish) and maintained at the favorable temperature (26°C) (***Chi et al., 2020***). The stage-specific number of individuals was recorded daily (food replaced every two days). During the pupal stage, each pupa was placed in a perforated 1.5 mL centrifuge tube. After eclosion, adults were transferred to 50 mL plastic cups for mating and oviposition and fed with a 10% honey solution. We continuously monitored and recorded daily oviposition and the number of surviving adults within the population until all individuals died.

For the individual life tables, 120 randomly selected newly laid eggs from each strain (i.e., 120 eggs/strain/temperature) were placed under different temperature conditions (favorable, hot, and cold environments). The eggs were individually transferred to 30 mm diameter petri dishes (artificial diet replaced every two days). The number of surviving larvae and pupae was recorded daily, and each pupa was placed individually in a perforated 1.5 ml centrifuge tube. One newly emerged adult female and one newly emerged adult male were placed in a 25 mL plastic cup and fed with a 10% honey solution for mating and oviposition. The number of eggs laid by each adult female was recorded daily until death. The longevity of both males and females was recorded.

Life history and population fitness parameters, including egg duration, larval duration, pupal duration, preadult duration, female and male longevity, oviposition days, fecundity, intrinsic rate of increase (*r*), finite rate of increase (λ), net reproductive rate (*R_0_*), and mean generation time (*T*), were calculated based on the recorded data of age-stage-specific two-sex life tables of different strains. The numerical computation was done using the TWOSEX-MSChart software (https://www.faas.cn/cms/sitemanage/index.shtml?siteId=810640925913080000) and the bootstrap method with 100,000 replications to obtain standard errors of the fitness parameters. The paired bootstrap (BT) method with 100,000 replications was used to assess pairwise differences among all strains, including comparisons among HS, AS, and CS, as well as among the ancestral and SODC-MU mutant strains under each thermal regime. A *P* value of less than 0.05 indicates a statistically significant difference (***Chi, 1988***).

### Metabolomic and transcriptomic profiling

All samples for omics profiling were collected after two generations of rearing under common conditions (26°C in climate-controlled chambers) to capture constitutive, genetically based differences between the strains, avoiding confounding effects of immediate physiological plasticity that would occur under thermal stress. The 3^rd^-instar larval stage was selected as the most actively feeding and rapidly growing stage, where metabolic demands and energetic trade-offs critical for adaptation are most pronounced. Samples collected for metabolomic profiling included: (1) the 3^rd^-instar larvae from different strains (AS, HS and CS) maintained at the favorable temperature (26°C); and (2) the 2-day-old eggs, 1-day-old 3^rd^-instar larvae, 2-day-old pupae and newly emerged adults from the ancestral and SODC-MU (MU1 and MU2) strains (see section Deletion of the targeted gene using the CRISPR/Cas9 genome editing) at the favorable temperature. Each biological replicate consisted of pooled individuals at the same developmental stage. Six independent biological replicates per strain were used for metabolomic profiling and three for transcriptomic profiling. The same sample collection strategy was applied to both analyses, but replicates were collected independently.

### Metabolomic profiling

To identify metabolic changes associated with thermal adaptation, we performed targeted metabolomic profiling using UPLC-MS/MS. Stored samples were thawed on ice and weighed (50 ± 2 mg) into 1.5 mL centrifuge tubes, to which three pre-cooled steel balls (3 mm) and 500 µL of pre-cooled 70% methanol (Merck, Germany) were added. Each of the samples was homogenized in a pre-cooled tissue homogenizer (25 HZ, 5 min) (Tissuelyser, Qiagen), and then the homogenized sample was left on ice for 15 min, then centrifuged at 12,000 rpm at 4°C for 10 min. The supernatant was transferred to a new 1.5 mL centrifuge tube and stored overnight at -20°C. The following day, the samples were centrifuged at 12,000 rpm for 3 minutes at 4°C. The supernatant was then collected with a sterile syringe and filtered through a 0.22 µm filter (Waters, USA) into an HPLC sample vial. The instrumental system for data acquisitions mainly used ultra-high performance liquid chromatography and tandem mass spectrometry (multiple reaction monitoring mode). The chromatography and tandem mass spectrometry conditions were as described by Li et al (***2021***). For all metabolomic comparisons (HS *vs*. AS, CS *vs*. AS, and SODC-MU *vs*. AS), differential metabolites were identified through pairwise comparisons using Student’s t-test with false discovery rate (FDR) correction. A multi-criteria threshold of |log_2_Fold Change| ≥ 1, VIP (variable importance in projection) ≥ 1, and FDR < 0.05 was applied. All differential metabolites were assigned to different pathways by KEGG analysis.

### RNA extraction and cDNA synthesis

To obtain templates for gene cloning and qRT-PCR analysis, total RNA was extracted using the Eastep® Super Total RNA Extraction Kit (Promega, Shanghai) according to the manufacturer’s instructions. RNA integrity and quality were assessed using a NanoDrop 2000 spectrophotometer (GE Healthcare, USA) and 2% agarose gel electrophoresis. Total RNA (2000 ng) was reverse transcribed into cDNA using the Reverse-Transcription System Kit (Promega, Shanghai).

### Transcriptomic profiling

To identify gene expression changes associated with thermal adaptation, mRNA libraries were constructed for each of the samples and sequenced on the Illumina HiSeq4000 platform (Illumina, San Diego). Raw reads obtained from sequencing were filtered, low quality reads were removed using adapters, and clean reads were obtained for subsequent information analysis. Clean reads were aligned to the *P*. *xylostella* genome using HISAT2 (V2.1.0) (http://121.37.197.72:5010), with sequence alignment performed using the software’s default parameters. Gene expression levels were measured using FPKM (fragments per kilobase of transcript per million fragments mapped). The *P*-value was corrected for multiple hypothesis testing following Benjamini and Hochberg (***1995***). Differential expression analysis between samples was performed using DESeq2. Differentially expressed genes were identified using the criteria of |log_2_Fold Change| ≥ 1 and FDR (false discovery rate) < 0.05. All differentially expressed genes were classified into different pathways by KEGG analysis.

### Weighted gene co-expression network analysis (WGCNA)

To identify gene modules correlated with differential metabolites between thermally adapted and ancestral strains, we performed weighted gene co-expression network analysis (WGCNA). Genes with an FPKM value below 0.1 were filtered out. The WGCNA package in R (v2.0) was used to calculate weight values and a soft threshold was determined based on the scale-free network principle. A gene clustering tree was constructed based on gene expression correlations, and gene modules were identified based on these clustering relationships. The minimum number of genes per module was set to 30, and the cut height threshold was set to 0.25 to merge potentially similar modules, with other parameters set to default values. Differential metabolites shared between the hot and cold strains relative to the ancestral strain were used as trait data for correlation analysis, as metabolites represent intermediate molecular phenotypes that bridge gene expression and organismal-level traits, enabling more direct identification of functionally relevant gene modules. Pearson correlations were calculated between each module eigengene and each of the 30 common differential metabolites (29 modules × 30 metabolites = 870 tests). Following standard WGCNA practice, a stringent dual threshold of |correlation coefficient| > 0.8 and *P* < 0.05 was applied to identify significant module-metabolite associations, effectively controlling for false positives (***Langfelder and Horvath, 2008***). Genes within specific modules were compared with differentially expressed genes and those common to both were considered as candidate genes.

### Gene cloning

To identify non-synonymous mutations in candidate genes across the different strains, the following gene sequences were amplified: (1) differentially expressed genes identified from specific modules; and (2) genes from the *PxSODC* gene family. Reference sequences for these genes were obtained from the *P*. *xylostella* genome database and full-length primers were designed using Primer Premier 6.0 (***Supplementary File 7***). PCR amplifications were performed using cDNA from the ancestral, hot and cold strains of *P*. *xylostella* (at least six samples) as templates, using Phanta® Max Super-Fidelity DNA Polymerase (Vazyme, China). The 50 µL PCR reaction mixture contained 2× reaction buffer (25.0 µL), dNTP mix (1.0 µL), 20 µM upstream primer F (2.0 µL), 20 µM downstream primer R (2.0 µL), DNA polymerase (1.0 µL), nuclease-free water (15.0 µL) and cDNA (4.0 µL). Amplification conditions were as follows 95°C for 3 minutes, followed by 35 cycles of 95°C for 15 seconds, 58°C for 15 seconds and 72°C for a gene-specific extension time, with a final extension at 72°C for 5 minutes. The integrity of the PCR products was verified by 2% agarose gel electrophoresis and purified using the Omega gel extraction kit (USA). The purified fragments were cloned into the pESI-Blunt vector using the Hieff CloneTM Zero TOPO-Blunt Simple Cloning Kit (YEASEN, Shanghai, China) and transformed into competent DH5α cells (YEASEN, Shanghai). The ligated product was transferred to DH5α competent cells and plated on LB+100 μg/ml ampicillin plates and incubated overnight at 37°C. A single colony was picked and placed in 500 µL of liquid LB with 100 µg/mL ampicillin. A positive clone was sent to Sangon Biotech (Shanghai, China) for sequencing. Sequence alignments of candidate genes from different strains were performed using Snap Gene software.

### Sequence analysis and phylogenetic tree construction

To determine the exons and introns of the target genes, their sequences were aligned with gDNA from the *P*. *xylostella* genome database. Protein sequences were analyzed using a protein sequence analysis and classification tool (InterPro, http://www.ebi.ac.uk/interpro/). The relative molecular masses and isoelectric points of the proteins were predicted using Expasy (http://expasy.org/tools/dna.html). The secondary structures of the proteins were predicted using PSIPRED (http://bioinf.cs.ucl.ac.uk/psipred). A phylogenetic tree was inferred using the model-based Maximum Likelihood method implemented in IQ-TREE, and the robustness of the tree was verified by bootstrap analysis (bootstrap = 1000 replicates). In the absence of a valid outgroup sequence, the resulting gene tree was presented as unrooted

### Molecular dynamics (MD) simulations

To investigate the structural consequences of the identified non-synonymous mutations in *PxSODC* under thermal stress, molecular dynamics (MD) simulations were performed using GROMACS 2022.3. The 3D structures of the wild-type (WT) and mutant (MU) PxSODC proteins were predicted using AlphaFold2. The Amber99sb-ildn force field was applied to the protein systems. Each system was solvated in a cubic box with TIP3P water molecules and neutralized by adding appropriate NaC ions. Energy minimization was performed using the steepest descent algorithm. Subsequently, 100 ps of NVT (constant volume and temperature) equilibration followed by 100 ps of NPT (constant pressure and temperature) equilibration were conducted with a coupling constant of 0.1 ps. Production MD simulations were then run for 100 ns (5,000,000 steps with a 2 fs timestep) at three temperatures: 15°C (288.15 K, cold stress), 26°C (299.15 K, favorable baseline), and 32°C (305.15 K, heat stress). Structural stability and dynamic properties were evaluated using built-in GROMACS analysis tools, including root mean square deviation (RMSD), solvent accessible surface area (SASA), and intramolecular hydrogen bond number. The 26°C simulation served as the physiological baseline for interpreting stress-induced structural changes.

### qRT-PCR analysis

To validate transcriptomic data and quantify expression levels of candidate genes across strains and temperature conditions, we collected samples as follows: (1) we randomly selected 22 genes from the transcriptomes of the 3^rd^-instar larvae of ancestral, hot, and cold strains to validate the transcriptome data; (2) we collected samples from the eggs, 3^rd^-instar larvae, pupae and adult males and females of the ancestral, hot, cold and SODC-MU (MU1 and MU2) strains to assess the transcription levels of genes including *PxSODC*, *Px04C00505*, *Px13C00423*, *Px20C00248*, *Px15C00224* and *Px15C00223*; (3) we collected the 3^rd^-instar larvae of the ancestral and hot strains exposed to different high temperature treatments (32°C, 34°C, 36°C) to analyze the transcription levels of the above-mentioned genes; and (4) we collected the 3^rd^-instar larvae of the ancestral and cold strains exposed to different low temperature treatments (12°C, 10°C, 8°C) for similar assessments.

qRT-PCR primers were designed using Primer Premier 6.0, with *PxRPL^32^*as the reference gene (***Supplementary File 7***). The reaction mixture of 20 μL contained 10 μL 2× Taq Pro Universal qPCR Master Mix (Vazyme, Nanjing, China), 0.4 μL of each primer, 7.15 μL nuclease-free water and 2.0 μL cDNA. The QuantStudio Real-Time PCR System (Thermo, USA) protocol was as follows: 95°C for 30 s; 40 cycles of 95°C for 10 s, 60°C for 30 s; followed by melting curve analysis of 95°C for 15 s, 60°C for 1 min, 95°C for 15 s. Each sample contained 3 biological replicates and 3 technical replicates, and the relative expression of genes was calculated using the 2^-ΔΔCt^ method.

### Deletion of the targeted gene using the CRISPR/Cas9 genome editing

To functionally validate the role of *PxSODC* in thermal adaptation, we generated stable homozygous mutant strains of *P*. *xylostella* with the *PxSODC* gene deleted using the CRISPR/Cas9 system. The target site was designed based on the 5’-N20<u>NGG</u>-3’ motif (underscore indicates PAM sequence), and the potential off-target effect of sgRNAs was predicted using Cas-OFFinder (http://www.rgenome.net/cas-offinder). The in vitro transcription template for sgRNA was generated from a single nucleotide strand under the following conditions: 95°C for 3 min, followed by 35 cycles of 95°C for 15 s, 68°C for 15 s and 72°C for 30 s, with a final extension at 72°C for 5 min. The amplified product was purified by gel extraction. The sgRNA was obtained by in vitro transcription of the gel-purified product using the HiScribe™ T7 Quick High Yield RNA Synthesis Kit (New England Biolabs, USA). The reaction mixture contained 2.5 μL NTP buffer mix, 0.5 μL T7 RNA polymerase mix, 65 ng gel-purified product, made up to 5 μL with nuclease-free water. After overnight incubation at 37°C, 0.5 μL of DNase was added to remove DNA and the product was incubated at 37°C for 20 minutes to yield sgRNA. The sgRNA was purified by phenol-chloroform extraction and stored at -80°C.

We prepared a 10 μL reaction mixture containing 300 ng/μL sgRNA and 200 ng/μL Cas9 protein (GenCrispr, Nanjing), 1 μL 10× reaction buffer and nuclease-free water to make up to 10 μL and incubate at 37°C for 25 minutes. The mixture was injected into freshly laid eggs using the Olympus SZX16 microinjection system (Olympus, Japan) and the entire microinjection was completed within 30 minutes of eggs being laid. After injection, the eggs were placed in a Petri dish and the number of eggs hatched was recorded. Adult gDNA was extracted using the TiANamp Genomic DNA Kit (TIANGEN, China). Specific primers were designed for PCR amplification (***Supplementary File 7***), with conditions as follows: 95°C for 3 min, followed by 34 cycles of 95°C for 15 s, 58°C for 15 s and 72°C for 15 s, with a final extension at 72°C for 5 min. The sequence of the PCR products was checked by Sangon Biotech (Shanghai) Co., Ltd.

The injected eggs were referred to as the G0 generation. These were reared to adulthood, crossed with the ancestral (non-injected) adults and used to extract genomic DNA from G0 adults after oviposition (the resulting progeny representing the G1 generation). PCR products flanking the two sgRNA target sites were amplified as mentioned above to determine genotypes and identify heterozygotes (individuals with double peaks in the sequence chromatogram starting from the sgRNA target site). The G1 generation was self-crossed to produce the G2 generation, and all G1 adults were genotyped based on PCR amplification for individual identification. G2 progeny derived from G1 heterozygotes with the same allelic mutation were selected. The G2 generation was then self-crossed to produce the G3 generation, retaining those with the same type of homozygous mutations to establish homozygous lines. If the G3 generation remained heterozygous, self-crossing continued until homozygous mutations were obtained (***Wang et al., 2020***). By the end, three mutants were obtained and called SODC-MU (MU1, MU2 and MU3) strains.

### RNA interference mediated silencing of target genes

To assess the role of *PxDnmt1* in thermal tolerance, we silenced its expression using RNA interference. Gene-specific primers containing T7 promoter sequences were designed (***Supplementary File 7***), and PCR was performed using total *P. xylostella* cDNA as a template. The PCR products were purified using a gel extraction kit. Double-stranded RNA (dsRNA) was synthesized by in vitro transcription using the T7 High Yield RNA Transcription Kit (Vazyme, Nanjing, China). The dsRNA was diluted to 2 μg/μL using DEPC-treated water (Beyotime, Shanghai, China). A volume of 500 nL diluted dsRNA (ds*EGFP* or ds*Dnmt1*) was injected into pupae using a Nanoliter 2000 microinjector (World Precision Instruments LLC, USA). Total RNA was extracted 24 hours after injection and reverse transcribed to cDNA. Gene knockdown efficiency was analyzed by qPCR using pupae injected with ds*EGFP* as controls. The experiment was performed in three independent biological replicates (***Zhou et al., 2024***).

### Assessing the response to high temperature

To assess the response of different *P*. *xylostella* strains (ancestral strain, hot strain and mutants) to extremely high temperatures, 2-day-old eggs, 1-day-old 3^rd^-instar larvae and 2-day-old pupae were individually placed in 90 mm diameter Petri dishes. Adult females and males were placed individually in perforated 1.5 mL centrifuge tubes. Based on a preliminary trial on the stage-specific temperature tolerance limit of *P*. *xylostella* (eggs, larvae, pupae, and both male and female adults of the ancestral and hot strains were placed in different temperature environments ranging from 40 to 45°C), pupae from the ancestral and mutant strains were exposed to 43°C while eggs, larvae, and adults were exposed to 42°C for periods ranging from 30 to 180 minutes. After treatment, all replicate samples were transferred to an environment maintained at 26°C, where survival was observed and recorded. Survival was defined as the successful development of eggs, larvae and pupae to the next stage, while adults had to show movement of an appendage or mouthparts. Experiments were performed with six biological replicates, with each replicate contained 20 individuals.

We randomly selected 20 female adults injected with dsRNA to determine their critical thermal maximum (CTMax). A thermistor probe (Omega, USA) was inserted into a 1.5 mL centrifuge tube, which was suspended inside a 50 mL centrifuge tube with the opening sealed with cotton. This assembly was then placed in a 2 L glass beaker containing 1000 mL water, with the beaker top sealed with insulating foam board. The entire setup was positioned on a thermostatically controlled magnetic stirrer, where the temperature inside the 1.5 mL tube was increased at a constant rate of 0.5°C/min. When the temperature reached 26°C, female adults were quickly transferred into the 1.5 mL centrifuge tube containing the temperature probe, and their behavioral responses were continuously monitored as temperature increased. The CTMax was recorded when moths exhibited spasms, lost their crawling or flying ability, and remained motionless at the bottom of the tube, typically lying ventral side up (in most cases) or dorsal side up (in fewer instances). Although antennae and limbs might still exhibit slight tremors at this point, the insects typically died within seconds (***Li et al., 2024b***).

### Measurement of the supercooling and freezing points

To investigate the cold hardiness of different *P*. *xylostella* strains (including the ancestral, cold and mutant trains), we randomly selected 40 pupae from each strain to examine their supercooling and freezing points. A thermistor probe from a subcooling point tester (Omega, USA) was attached to a pupa, secured with conductive tape and placed in a centrifuge tube, with the tube mouth sealed by cotton. The centrifuge tubes were then placed in a 50 mL plastic cup filled with ethylene glycol (antifreeze), and the cup was stored in an ultra-low temperature freezer set at -70°C, with the temperature first dropping rapidly and then decreasing at a rate of 0.10°C per second until the supercooling point was reached. By recording temperature changes at intervals of every second, the supercooling and freezing points of pupae were determined based on the inflection point of body temperature. The same experimental approach was also applied to *P. xylostella* injected with dsRNA.

### Detection of oxidative stress indicator

To assess the impact of thermal adaptation and *PxSODC* deletion on oxidative stress, we measured superoxide dismutase (SOD) activity and superoxide anion (O_2_ ^-^) levels. Samples were collected in the following conditions: (1) the eggs, 3^rd^-instar larvae, pupae, and adult males and females of the ancestral, hot, cold, and SODC-MU (MU1 and MU2) strains at the favorable temperature (26°C); (2) the 3^rd^-instar larvae of the ancestral, hot, and SODC-MU (MU1 and MU2) strains after 2 hours of heat stress at 32°C, 34°C, and 36°C; and (3) the 3^rd^-instar larvae of the ancestral, cold, and SODC-MU (MU1 and MU2) strains after 2 hours of cold stress at 12°C, 10°C, and 8°C. The experiment was performed with four independent biological replicates. The levels of SOD and O_2_^-^ were measured using commercial assay kits (Comin, China) according to the manufacturer’s instructions.

### Detection of 5-methylcytosine concentration

To evaluate the effect of *PxDnmt1* silencing on DNA methylation levels, pupae and female adults were collected for detection of 5-methylcytosine (5-mC) concentration after injection with dsRNA. The levels of 5-mC were measured using a commercial insect 5-methylcytosine (5-mC) ELISA detection kit (Shanghai Enzyme-linked Biotechnology Co., Ltd., China) according to the manufacturer’s instructions. The experiment was performed with four independent biological replicates.

### Data analysis

Statistical analyses for life table parameters, metabolomic data, and transcriptomic data are described in their respective sections above. For all other experimental data (qRT-PCR, SOD activity, O_2_^-^ levels, DNA methyltransferase activity, 5-mC concentration, stage-specific survival rates, and supercooling/freezing points), analyses were performed using SPSS version 23.0. The Shapiro-Wilk test was used to assess normality of data distribution. For normally distributed data, two-group comparisons were analyzed using independent samples t-tests, while comparisons involving three or more groups were analyzed using one-way ANOVA followed by Tukey’s test (homogeneous variances) or Tamhane’s T2 test (unequal variances). For non-normally distributed data, the Mann-Whitney test (two groups) or Kruskal-Wallis test (three or more groups) was used. A *P* value of less than 0.05 was considered statistically significant in all cases (***Lei et al., 2024***).

## Acknowledgements

This work was financially supported by the central government-guided local science and technology development project (2022L3087), the Fujian Natural Science Fund for Distinguished Young Scholars (2022J06013), and the research project grants of Fujian Agriculture and Forestry University (KFXH23021, KFB24114A), the State Key Laboratory of Agriculture and Forestry Biosecurity, the International Joint Research Laboratory of Ecological Pest Control, Ministerial and Provincial Joint Innovation Centre for Safety Production of Cross-Strait Crops, and the “111” Program in China.

## Additional information

### Funding

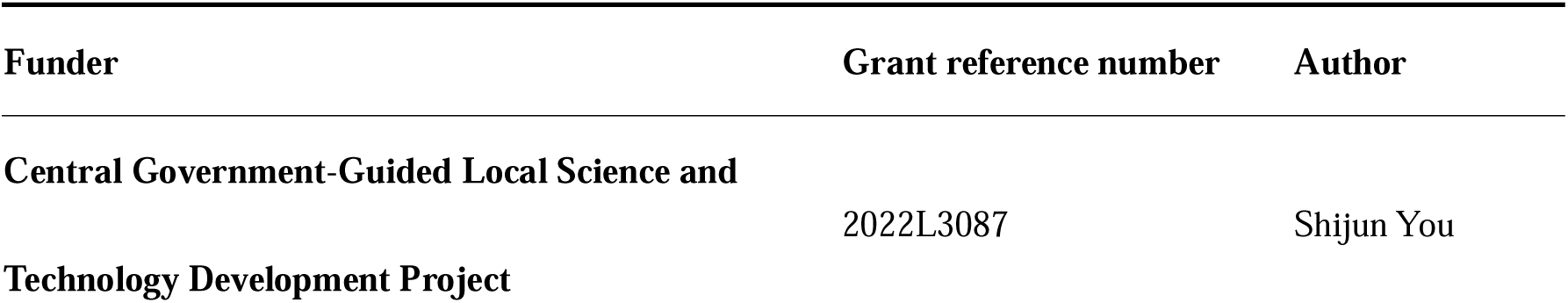
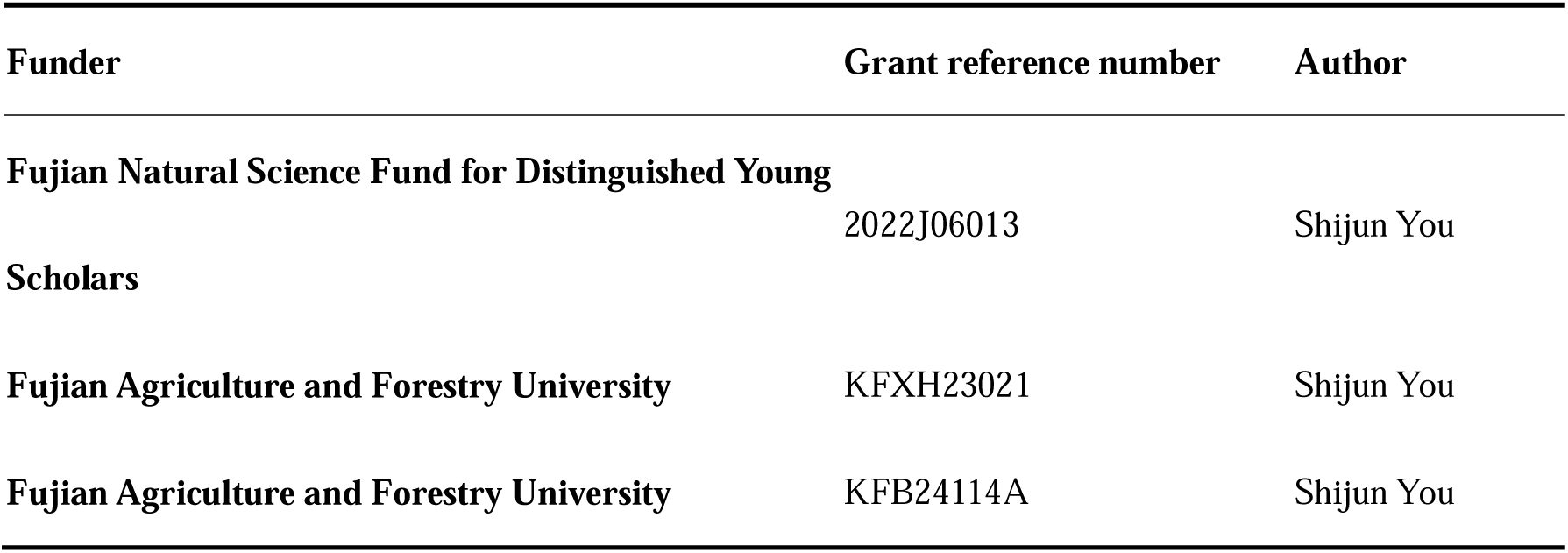

### Author contributions

Gaoke Lei, Conceptualization, Data curation, Methodology, Investigation, Writing – original draft;

Huiling Zhou, Conceptualization, Data curation, Methodology, Investigation, Writing – original draft;

Zongyao Ma, Data curation, Formal analysis, Investigation, Methodology;

Yating Duan, Data curation, Formal analysis, Investigation, Methodology;

Yanting Chen, Conceptualization, Methodology;

Fengluan Yao, Conceptualization, Methodology;

Minsheng You, Conceptualization, Data curation, Methodology, Writing – original draft, Writing – review and editing;

Liette Vasseur, Supervision, Writing – review and editing;

Geoff M. Gurr, Supervision, Writing – review and editing;

Shijun You, Conceptualization, Data curation, Methodology, Project administration, Supervision, Writing – original draft, Writing – review and editing.

## Competing interests

The authors declare that no competing interests exist.

## Additional files

### Supplementary files

Supplementary File 1. Life table parameters of the ancestral, hot and cold strains of *P. xylostella* at the favorable temperature (26°C).

Supplementary File 2. Candidate genes linked to thermal adaptation in DBM.

Supplementary File 3. Population fitness parameters of the ancestral strain (AS) and SODC-mutant strains of DBM under the constant favorable environment (26°C).

Supplementary File 4. Populaion fitness parameters of the ancestral strain (AS) and mutant strains of DBM under the hot environment (32°C/27°C:12 h/12 h).

Supplementary File 5. Population fitness parameters of the ancestral strain (AS) and mutant strains of DBM under the cold environment (15°C/10°C:12 h/12 h).

Supplementary File 6. Information on the *PxSODC* homologous genes identified in the transcriptome.

Supplementary File 7. The primers used in this study.

### Source data files

Figure 1—Source Data 1. Raw data for stage-specific thermal tolerance and pupal supercooling/freezing points of temperature-adapted strains.

Figure 5—Source Data 1. Raw data for thermal tolerance phenotypes of *PxSODC* knockout strains.

Figure 6—Source Data 1. Raw data for SOD enzyme activity and superoxide anion (OCC) levels across strains, developmental stages, and temperature conditions.

Figure 7—Source Data 1. Raw data for *PxDnmt1* expression, DNA methyltransferase activity, and effects of *PxDnmt1* silencing on thermal tolerance.

## MDAR checklist

### Data availability

The raw sequence data generated in this study have been deposited in the Genome Sequence Archive at the National Genomics Data Center, China National Center for Bioinformation / Beijing Institute of Genomics, Chinese Academy of Sciences, under accession number CRA024611. The associated metadata are available in the OMIX database under accession numbers OMIX009807 and OMIX009846. All other data generated or analyzed during this study are included in the manuscript and its supporting files.

The following datasets were generated:

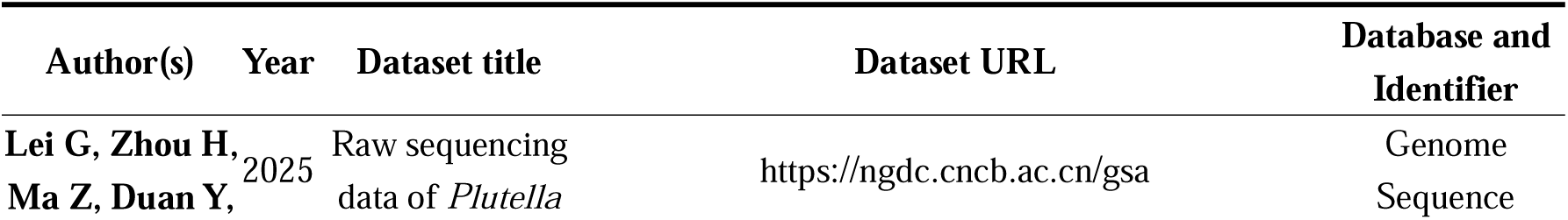
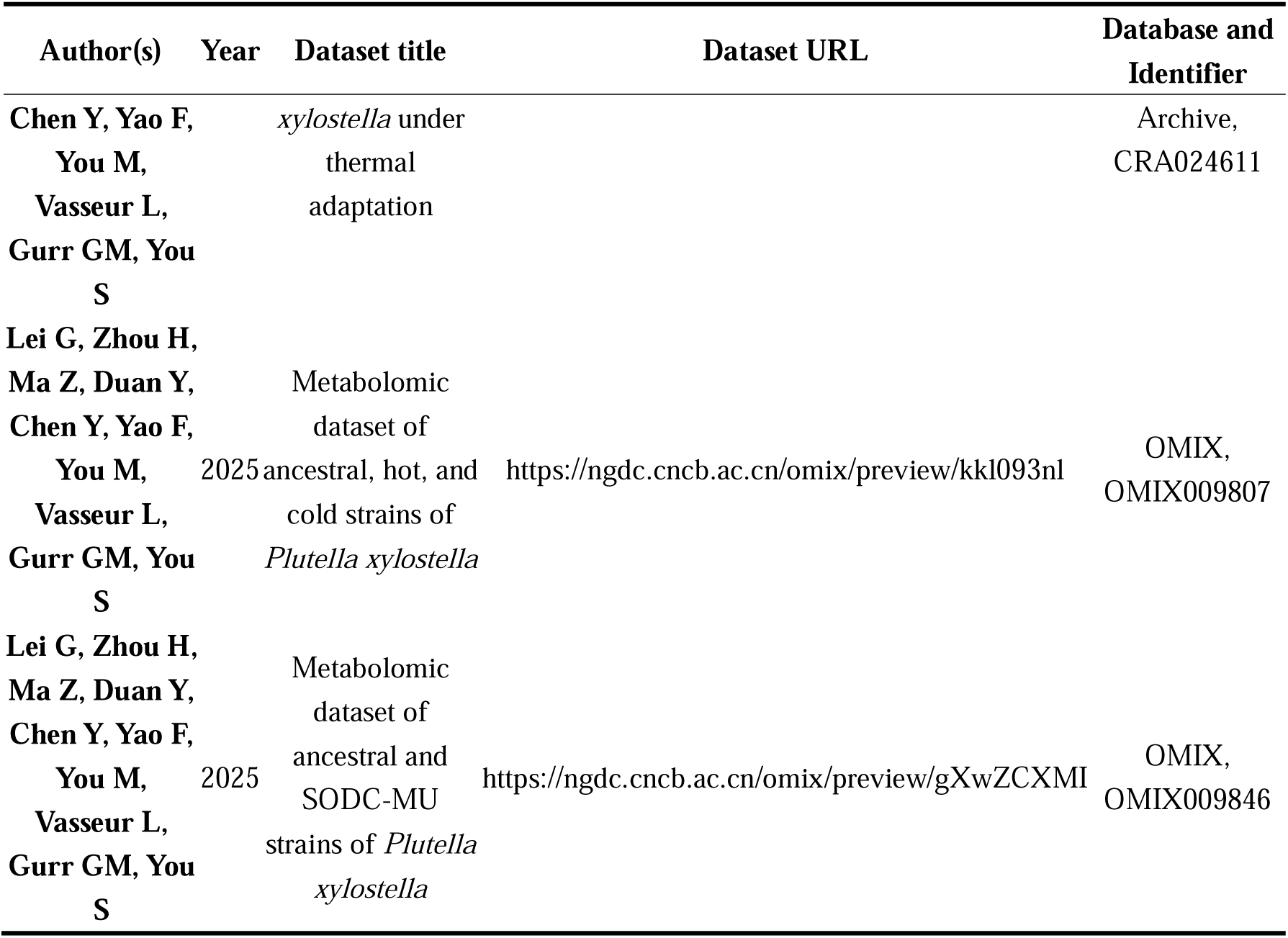

## Appendix

## Appendix 1: “Detailed age-stage survival and fecundity analysis”

To characterize how thermal adaptation affects population dynamics across different life stages, we generated curves of the age-stage specific survival rate (*s_xj_*), age-specific survival rate (*l_x_*), female age-specific fecundity (*f_x_*), and population age-specific fecundity (*m_x_*) from the age-stage, two-sex life tables of the hot strain (HS), cold strain (CS), and ancestral strain (AS) (***Figure 1–figure supplement 1***). These curves capture stage-specific variation in survival and fecundity that summary life table parameters alone may obscure, providing a more detailed view of how thermal selection shapes life history trajectories at each developmental stage. The maximum daily survival rates at the pre-adult stages of HS and AS were significantly higher than that of CS (HS = 78.9%, AS = 76.7%, CS = 66.7%; Supplemental Table S1). The peak daily fecundity of HS and AS was also higher than that of CS (HS = 53.22, AS = 53.00, CS = 45.10 eggs per female).

## Appendix 2: “Life table analysis of *PxSODC* mutant strains”

Age-stage, two-sex life tables were constructed for the ancestral strain and the three SODC-MU mutant (MU1, MU2, MU3) strains under constant favorable (26°C), hot (32°C/27°C) and cold (15°C/10°C) environments to examine their phenotypic traits including development, survival, fecundity and fitness (***Figure 5–figure supplement 5*; *Supplementary File 3-5***). Overall, the three mutant strains had a prolonged development time (*T*), lower survival rates (*s_xj_* and *l_x_*), and reduction in the fecundity (*f_x_* and *m_x_*) and population fitness parameters (*r*, λ, *R_0_*) compared to the ancestral strain. The observed change in these parameters indicated that the loss of *PxSODC* gene function significantly affected the life history and population fitness of *P. xylostella*, particularly in the hot/cold environments. Prolonged developmental time and reduced fecundity suggest that the physiological functions of the mutant strains may be damaged when exposed to both high and low temperature conditions.

## Appendix 3: “Metabolomic profiling of *PxSODC* mutant strains”

To explore the metabolic networks associated with the *PxSODC* gene in *P. xylostella* and better understand the biological functions underlying their complex relationships, we performed an untargeted metabolomic analysis on the ancestral and mutant (SODC-MU1 and SODC-MU2) strains at different developmental stages. We detected 608 metabolites across the strains, including 167 organic acids and their derivatives, 102 nucleotides and their metabolites, 102 lipids, 92 amino acids and their metabolites, 33 heterocyclic compounds, 24 carbohydrates and their metabolites, and 88 other metabolites (***Figure 7–figure supplement 1A***). Principal component analysis (PCA) revealed a distinction between the ancestral and mutant strains along the first component (PC1). However, a trend of clustering was observed in the 3^rd^-instar larvae of ancestral and SODC-MU1 strains and the male adults of ancestral and SODC-MU2 strains, suggesting that the difference at these developmental stages may be driven by a few key metabolites (***Figure 7–figure supplement 1B***). Compared to ancestral strain, different metabolites were identified in the eggs, 3^rd^-instar larvae, pupae, female adults and male adults of mutant strains (***Figure 7–figure supplement 1C***). The mutant strains had 68, 103, 110, 77 and 35 common differential metabolites compared to the ancestral strain at different developmental stages of *P. xylostella* (***Figure 7–figure supplement 1D***). In addition, the lipid content significantly decreased in the 3^rd^-instar larvae, pupae and female adults, while the content of nucleotides and their metabolites increased in the eggs, 3^rd^-instar larvae and pupae following deletion of the *PxSODC* gene in *P. xylostella* (***Figure 7–figure supplement 2A***). The up-regulation and down-regulation of various metabolites may be associated with the antioxidant function of *PxSODC*. We observed that the deletion of *PxSODC* gene could lead to elevated levels of the O ^-^ within *P. xylostella* (***Figure 6I***).

We then performed enrichment analysis of differential metabolite KEGG pathways for the SODC-MU (MU1 and MU2) eggs, 3^rd^-instar larvae, pupae, female adults and male adults resulting in these differential metabolites wre distributed in 23, 28, 28, 22, and 18 pathways, respectively. They were mainly involving lipid metabolism, nucleotide metabolism, carbohydrate metabolism, cofactor and vitamin metabolism, and amino acid metabolism (***Figure 7–figure supplement 2B*)**. These pathways were related to the *PxSODC* gene and might have contributed to the temperature-adaptive capacity in *P. xylostella* through the regulation of biological functions/processes in different temperature environments.

---

**Figure 1-figure supplement 1.**
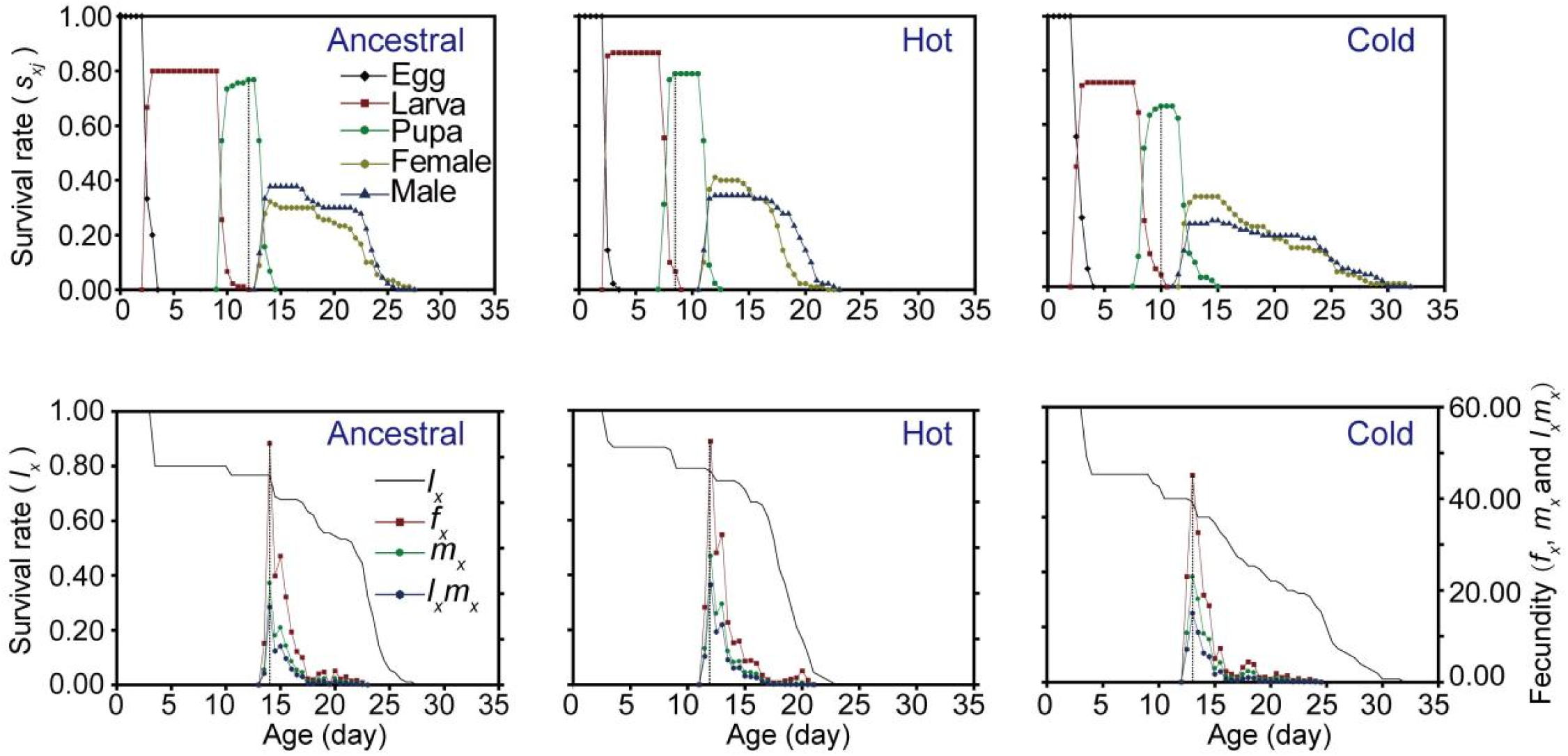
Phenotypic fitness variation in the ancestral, hot and cold strains of *P*. *xylostella*. The curves show the age-stage survival rate (*s_xj_*), age-specific survival rate (*l_x_*), female age-specific fecundity (*f_xj_*), and population age-specific fecundity (*m_x_*) of the ancestral, hot, and cold strains.

**Figure 2-figure supplement 1.**
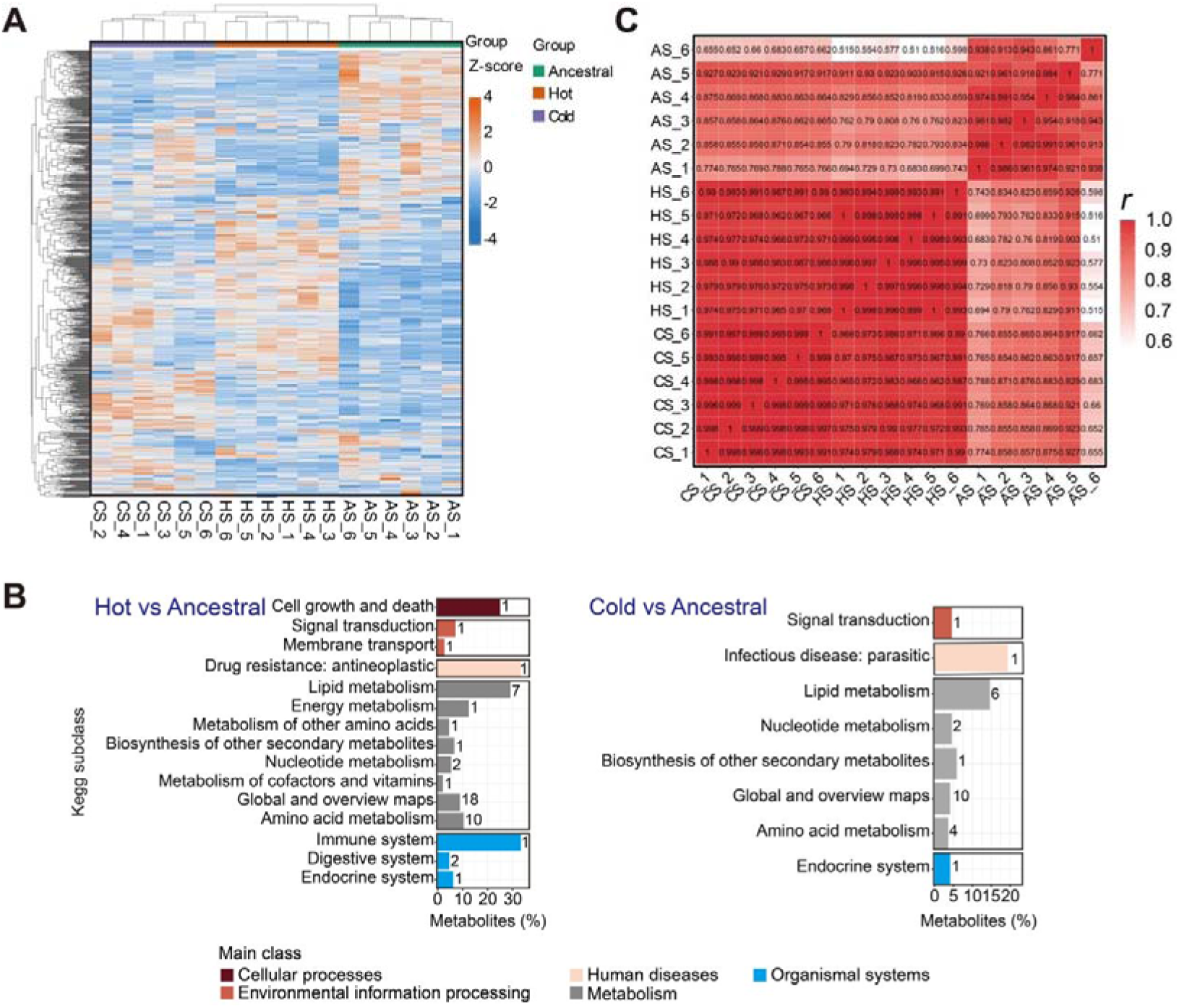
Metabolomic analysis of the 3^rd^-instar larvae across the ancestral, hot and cold strains of *P*. *xylostella*. (A) Inter-sample correlation heat map of the metabolites. Z-score standardized values for each of the metabolites were used in clustering analysis. The color bar indicates an increase in the content of each metabolite, scaling from blue to red. (B) KEGG functional classification of differential metabolites between the hot/cold and ancestral strains. (C) Correlation analysis of differential metabolites between the hot/cold and ancestral strains. Pearson’s correlation coefficient (*r*) was used to evaluate the biological correlation between different replicates.

**Figure 3-figure supplement 1.**
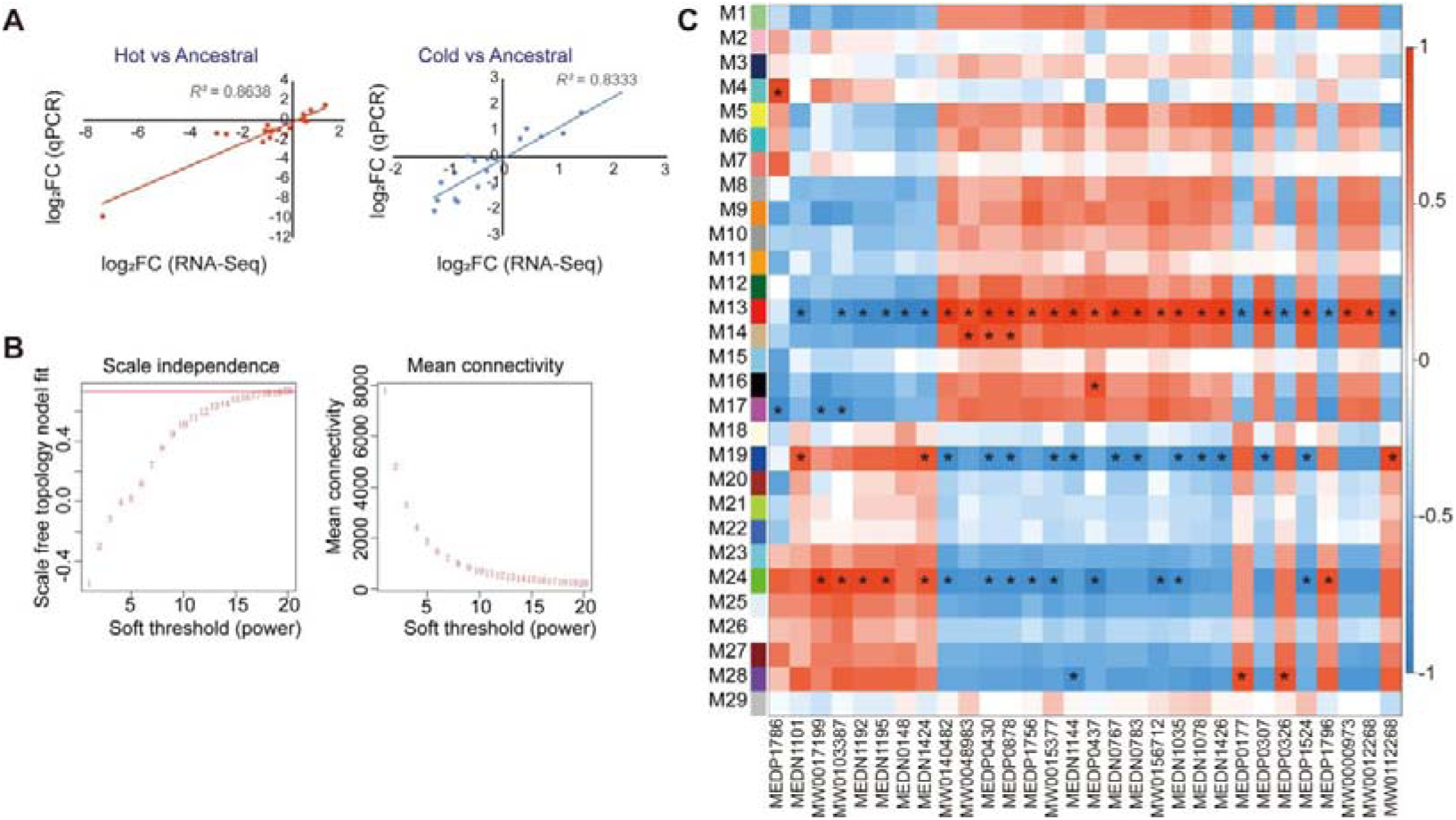
Transcriptomic analysis of the 3^rd^-instar larvae across the ancestral, hot and cold strains of *P*. *xylostella*. (A) qRT-PCR validation of transcriptome data. (B) Soft threshold selection in WGCNA. Left: The soft threshold selection graph. Right: The mean connectivity of genes under different soft thresholds. (C) Correlation heat map showing the association of the 29 modules with common differential metabolites between the hot/cold and ancestral strains.

**Figure 5-figure supplement 1.**
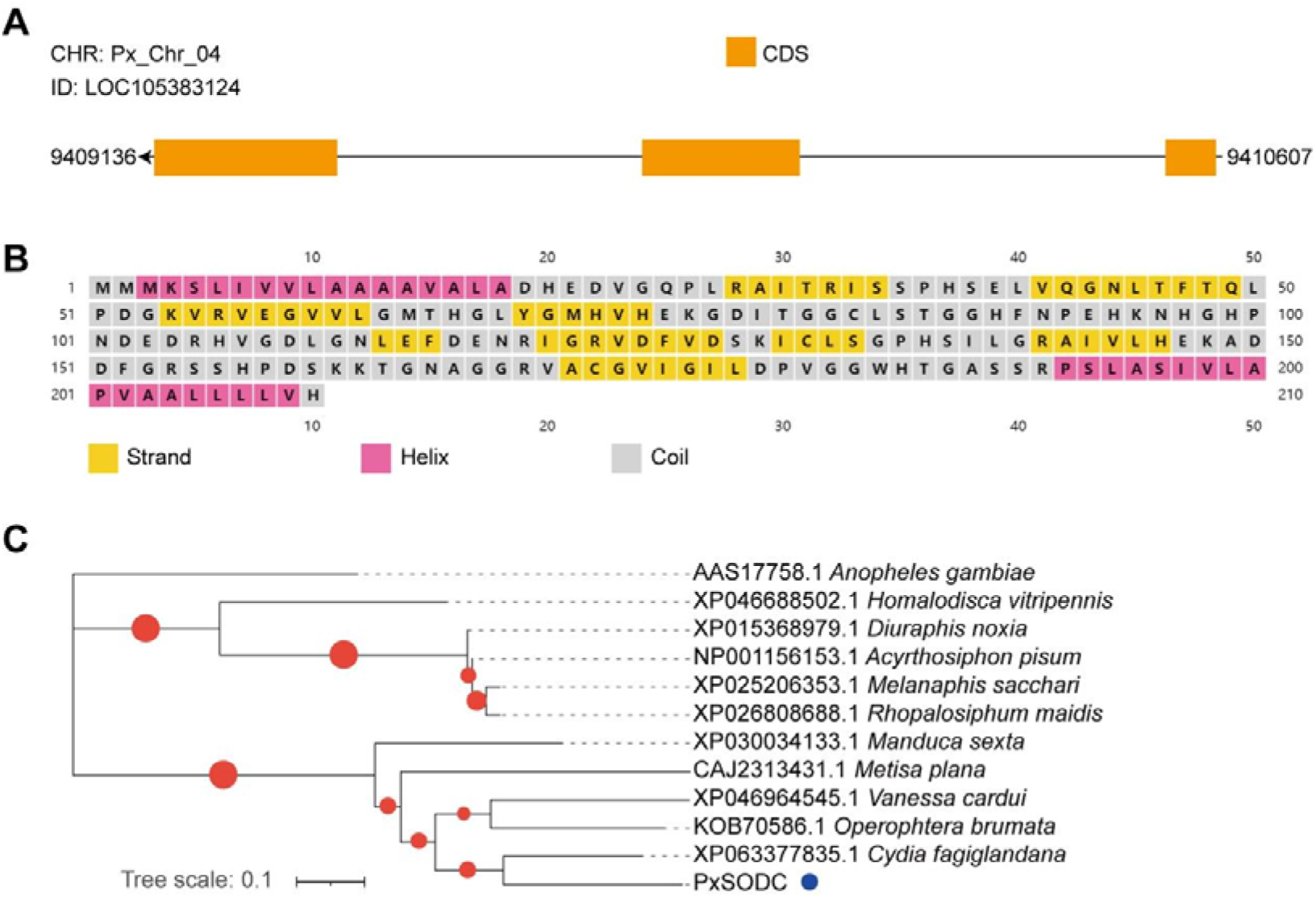
Bioinformatic prediction and analysis of the *PxSODC* gene sequence. (A) The structure of the *PxSODC* gene, consisting of three exons and two introns. (B) Secondary structure prediction of PxSODC. Yellow represents strand, pink represents helix, and grey represents coil. (C) Unrooted Maximum Likelihood phylogenetic tree of PxSODC based on amino acid sequences. The tree was inferred using IQ-TREE with 1000 bootstrap replicates, with a blue dot marking the position of *P*. *xylostella*.

**Figure 5-figure supplement 2.**
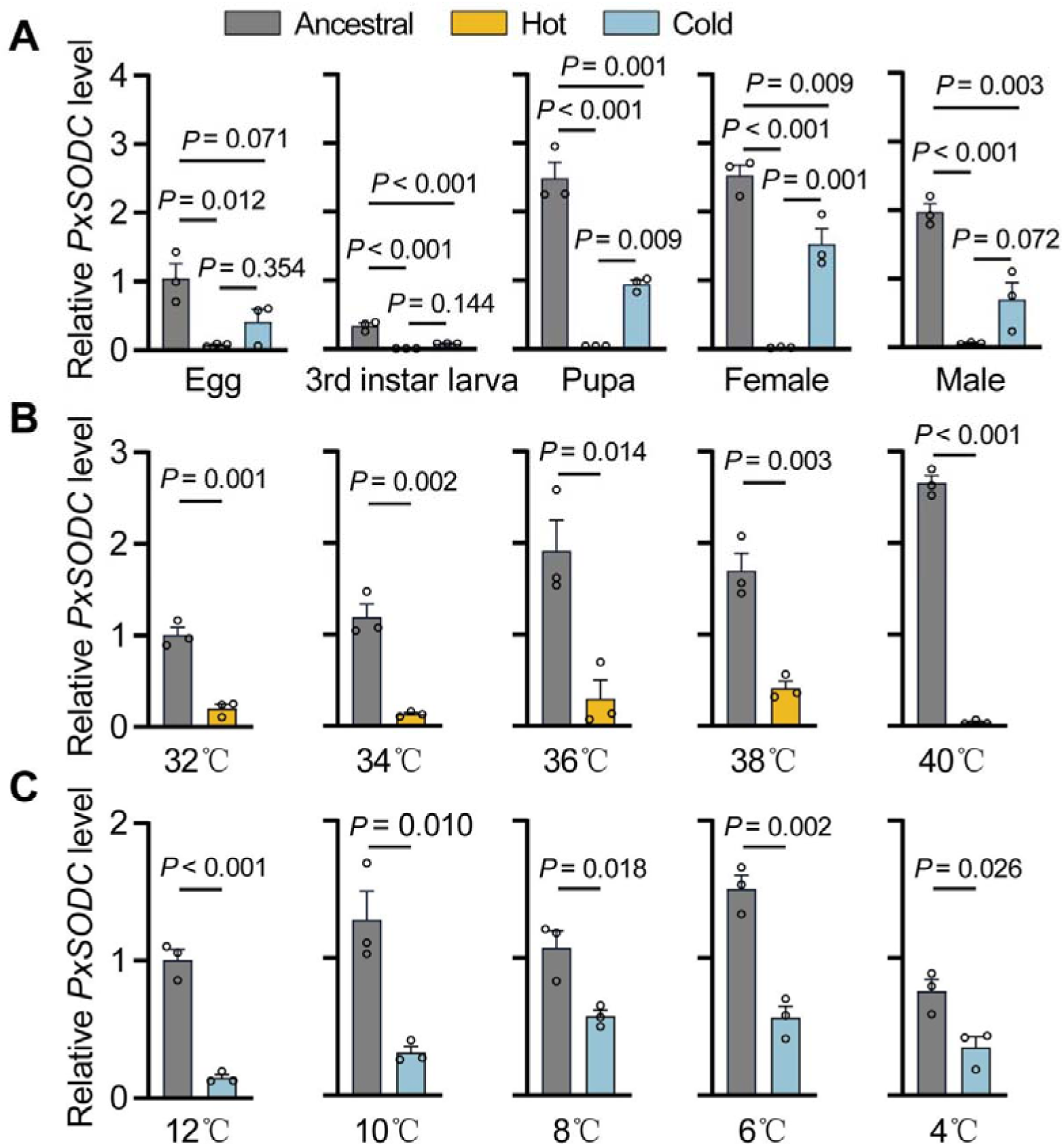
Spatio-temporal expression patterns of the *PxSODC* gene in the ancestral, hot and cold strains of *P*. *xylostella*. (A) Stage-specific profiling of the *PxSODC* expression patterns among different strains in the favorable temperature environment (26°C), and One-way ANOVA with Tukey’s test was used for comparison (*p* < 0.05). (B) Expression levels of *PxSODC* in the 3^rd^-instar larvae of the ancestral and hot strains after 2 h exposure to different high temperature environments (32°, 34°, 36°, 38° and 40°C), and t-test was used for comparison (*p* < 0.05). (C) Expression levels of *PxSODC* in the 3^rd^-instar larvae of the ancestral and cold strains after 2 h exposure to different low temperature environments (12°, 10°, 8°, 6° and 4°C), and t-test was used for comparison (*p* < 0.05). Data are presented as mean±SEM, with *n* = 3 biologically independent samples for each data point.

**Figure 5-figure supplement 3.**
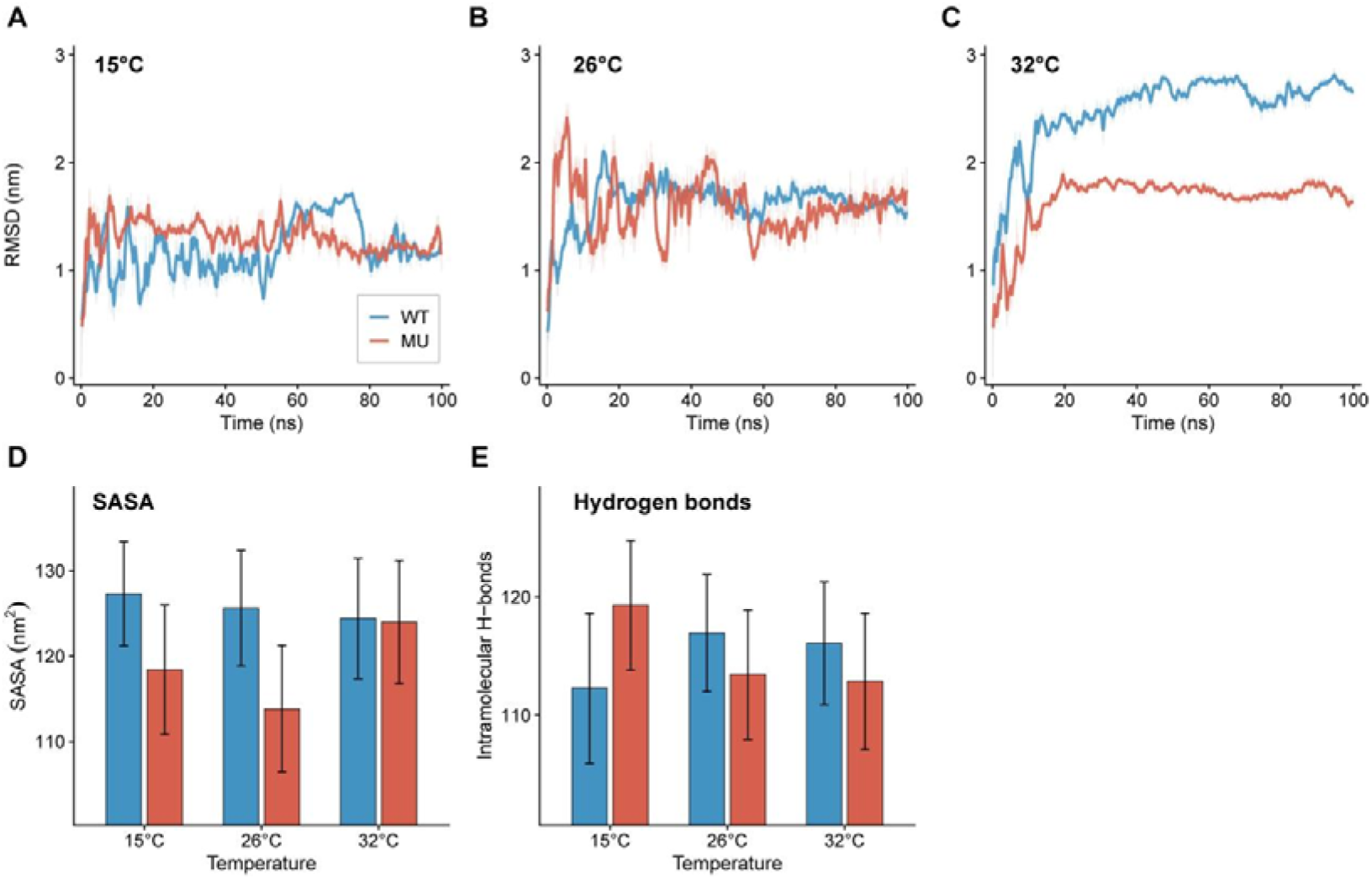
Molecular dynamics simulations of wild-type (WT) and mutant (MU) PxSODC proteins at 15°C, 26°C, and 32°C. The favorable temperature (26°C) served as the physiological baseline. (A-C) RMSD time-series trajectories over 100 ns at 15°C (A), 26°C (B), and 32°C (C). (D) Solvent accessible surface area (SASA) across three temperatures, demonstrating the more compact structure of MU at 15°C and 26°C. (E) Number of intramolecular hydrogen bonds across three temperatures. Under cold stress (15°C), MU actively increased hydrogen bonds compared to the 26°C baseline, whereas WT lost bonds. Under heat stress (32°C), MU fully maintained its bond count. Data in (D) and (E) are presented as mean ± SE from 100 ns simulations.

**Figure 5-figure supplement 4.**
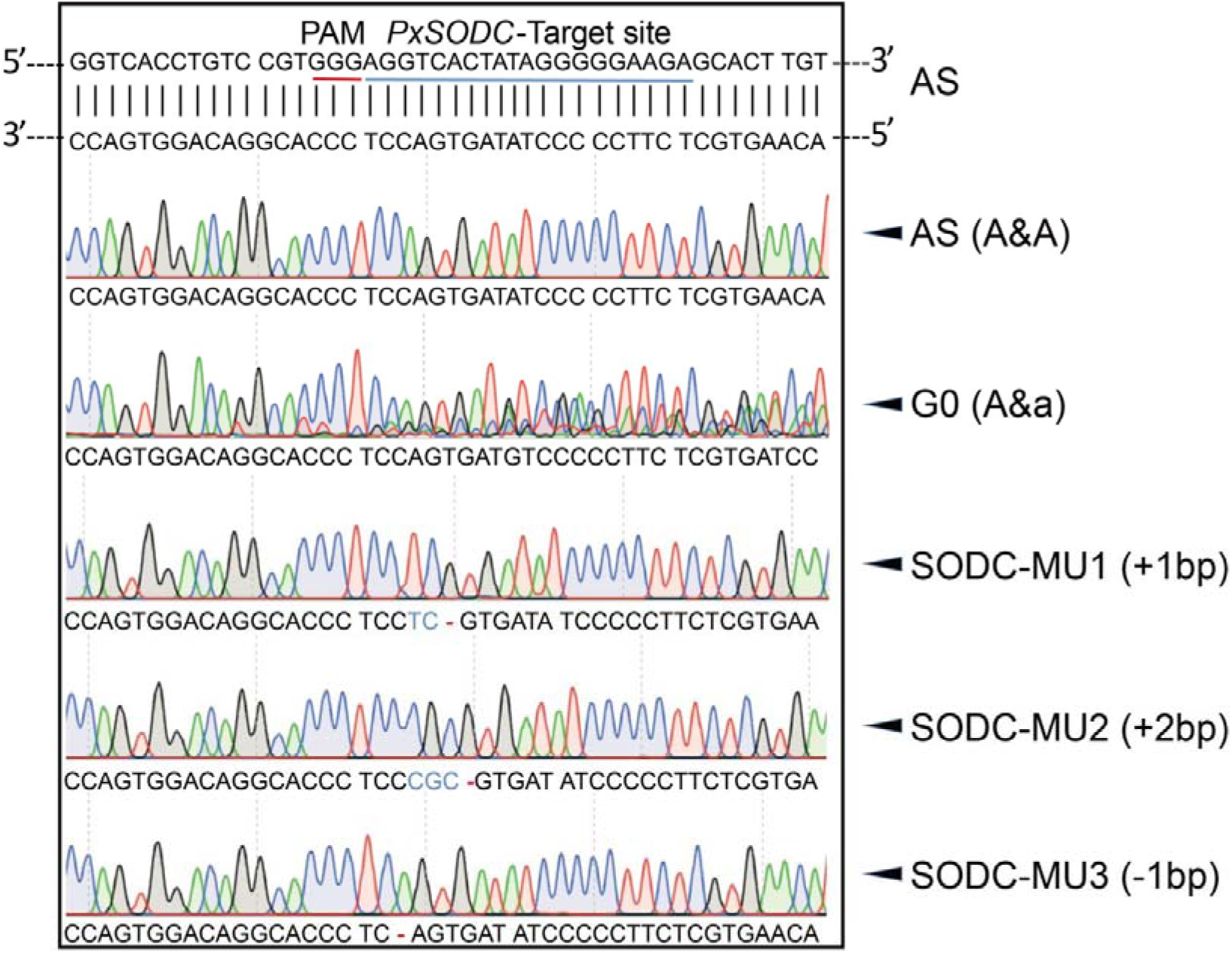
CRISPR/Cas9-mediated mutagenesis of the *PxSODC* gene. Black underlines indicate the sgRNA target sequence in the second exon, yellow underlines highlight the PAM recognition site, dashes indicate deleted base sequences and blue letters indicate inserted base sequences.

**Figure 5-figure supplement 5.**
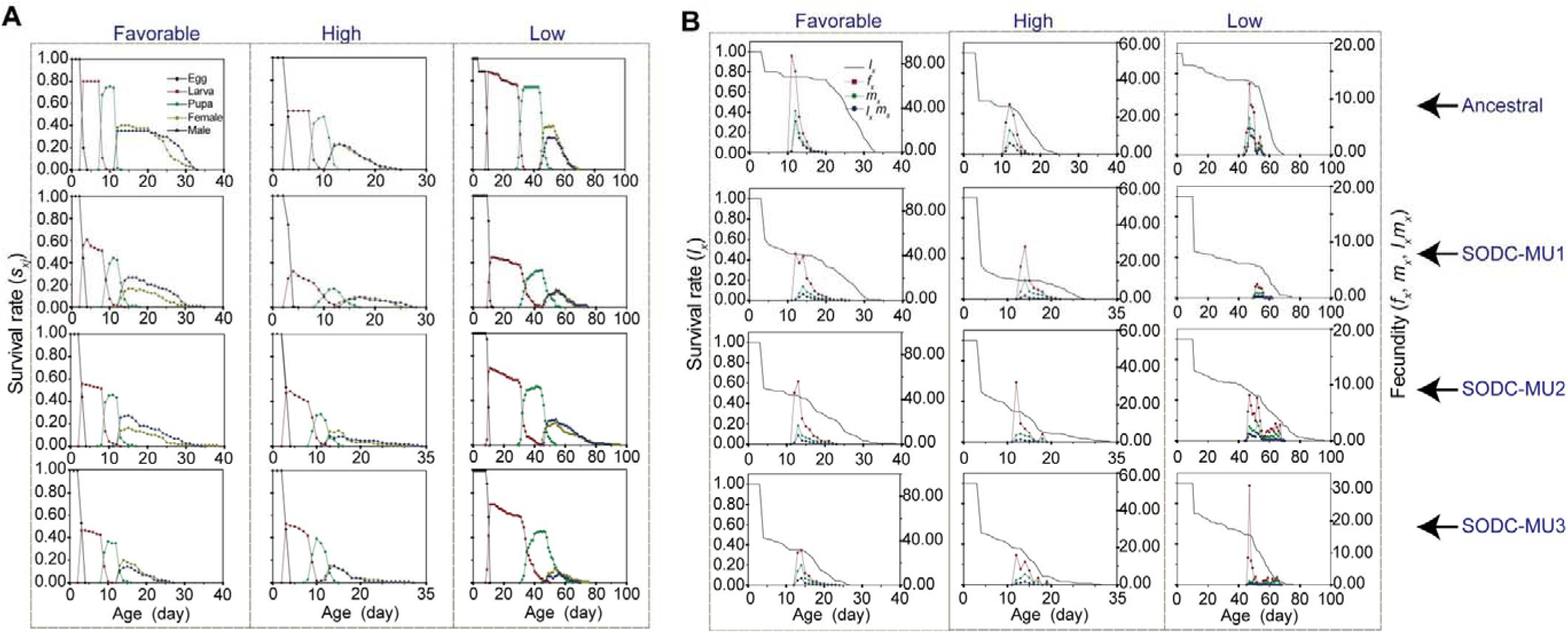
Comparison of the survival rates (*s_xj_* and *l_x_*) and female fecundity (*f_x_* and *m_x_*) of *P*. *xylostella* strains, showing the role of *PxSODC* in the temperature adaptation. (A) Age-stage survival rates (*s_xj_*) of the ancestral and mutant strains (AS, SODC-MU1, SODC-MU2 and SODC-MU3) in different temperature environments. (B) Age-specific survival rates (*l_x_*), female age-specific fecundity (*f_x_*), and population age-specific fecundity (*m_x_*) of the ancestral and mutant strains (AS, SODC-MU1, SODC-MU2 and SODC-MU3) in different temperature environments.

**Figure 7-figure supplement 1.**
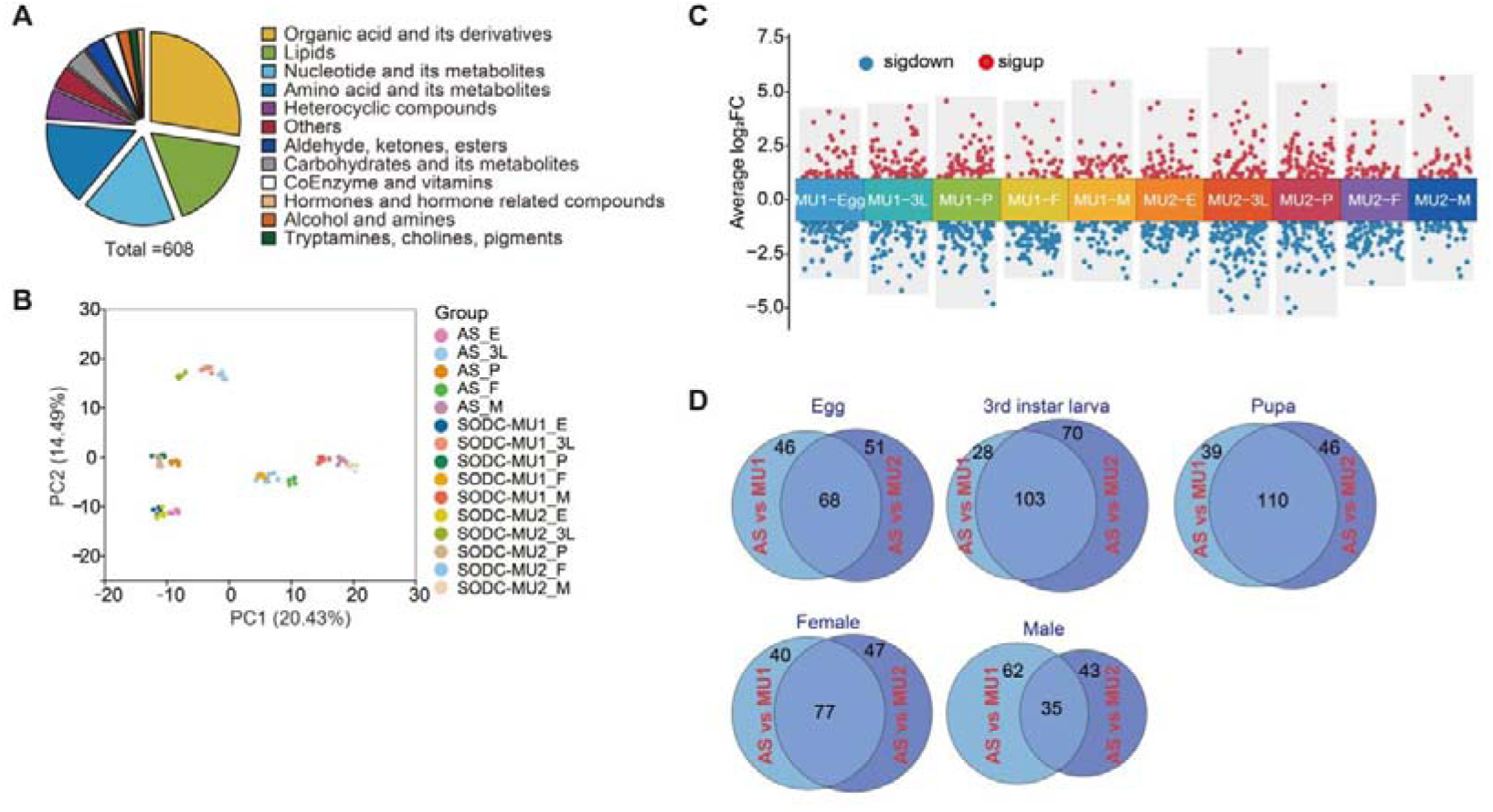
Metabolomic analysis of different developmental stages in the ancestral and mutant strains of *P*. *xylostella*. (A) Classification of metabolites. A total of 608 metabolites were detected in the ancestral and mutant strains (AS and SODC-MU1/SODC-MU2). (B) Principal component analysis (PCA) of the 608 metabolites in the ancestral and mutant strains (AS and SODC-MU1/SODC-MU2). PC1 and PC2 represent the first and second principal components, respectively. (C) Volcano plot showing down-regulated (blue dots) and up-regulated (red dots) metabolites identified in the mutant strains compared to the ancestral strain). (D) Venn diagram showing common and unique differential metabolites between the ancestral and mutant strains at different developmental stages.

**Figure 7-figure supplement 2.**
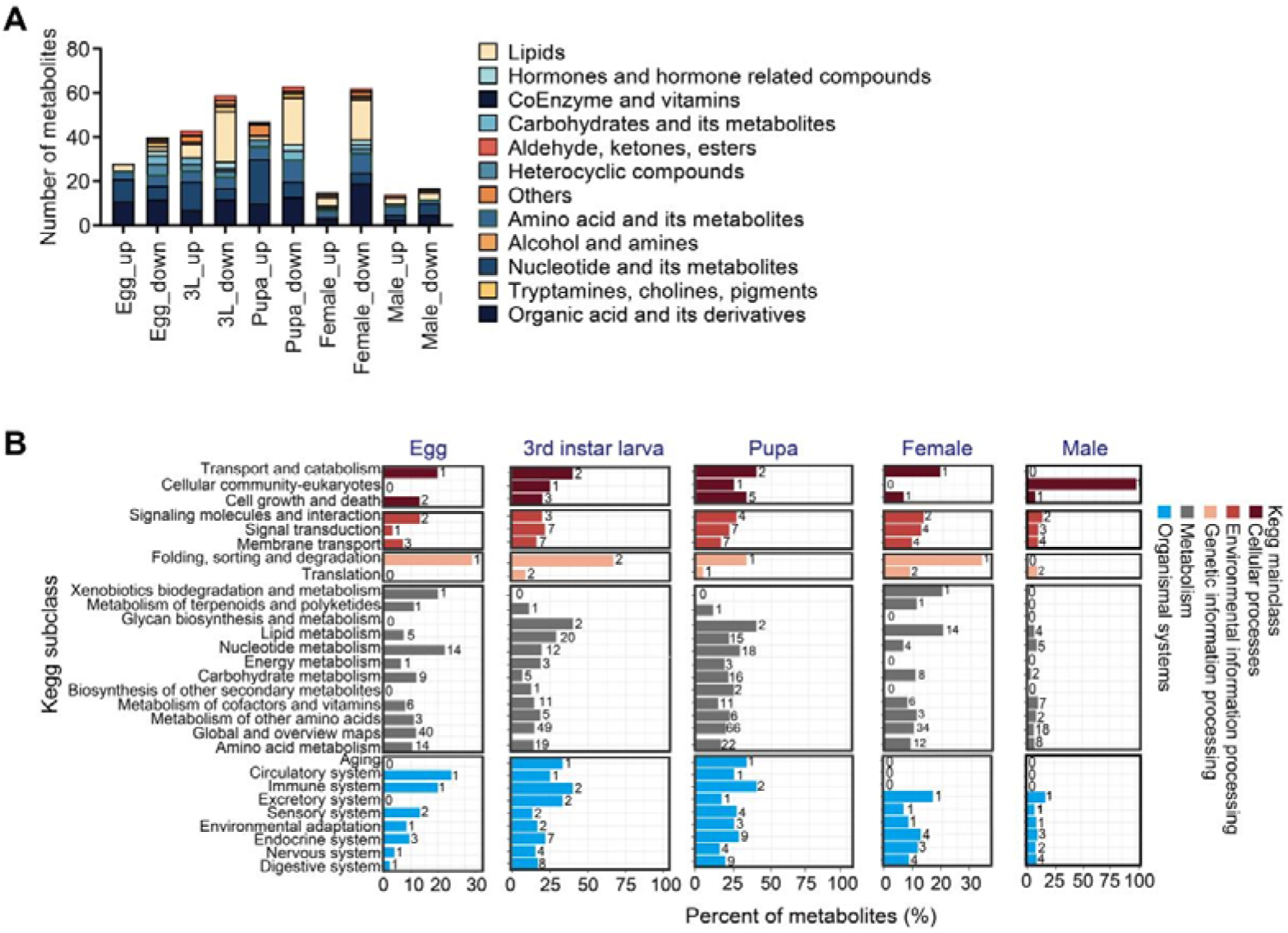
Differential metabolite classification and functional analysis in the ancestral and mutant strains across different developmental stages of *P*. *xylostella*. (A) Classification of differential metabolites between the ancestral and mutant strains at different developmental stages. (B) KEGG function classification of differential metabolites in the mutant strains compared to the ancestral strain. The vertical axis represents the names of the KEGG pathways and the horizontal axis indicates the number of differential metabolites annotated to each pathway and their proportion of the total metabolites annotated to that pathway.

**Supplementary File 1.**
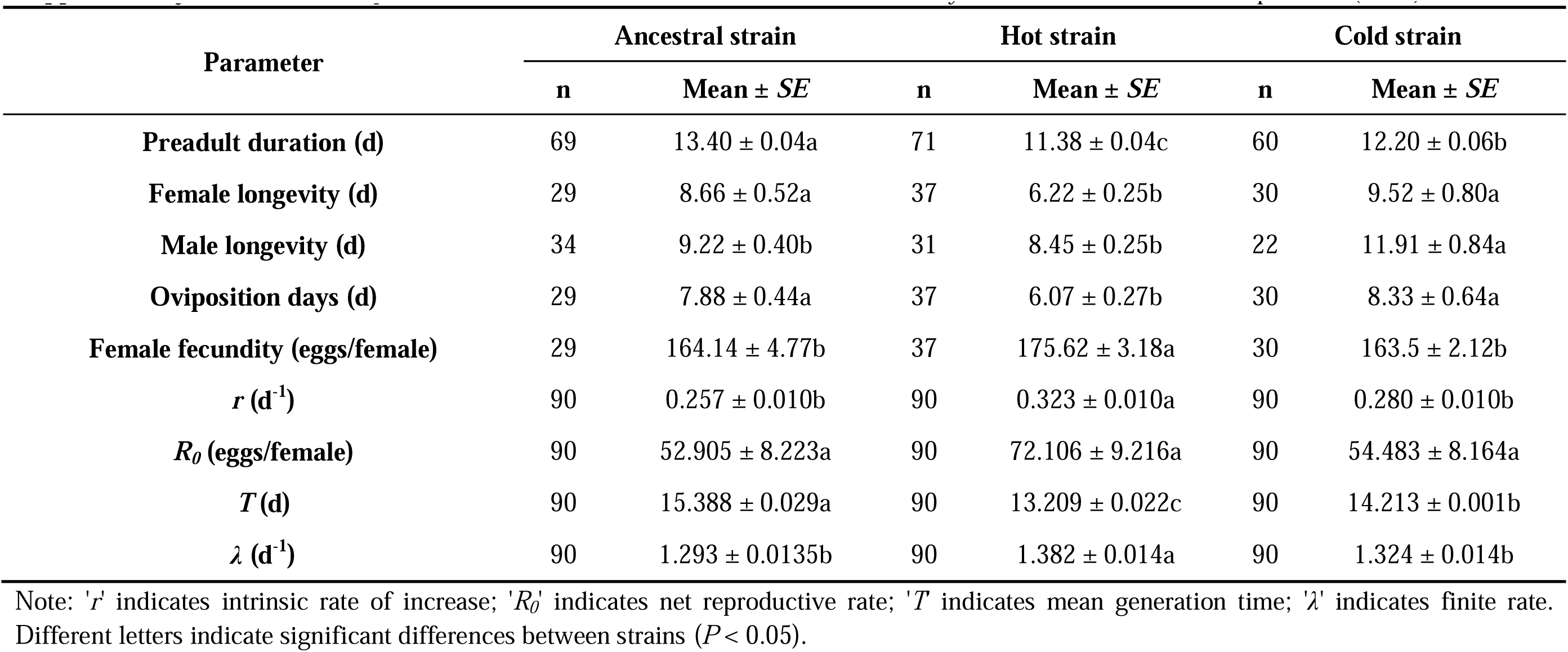
Life table parameters of the ancestral, hot and cold strains of *P. xylostella* at the favorable temperature (26°C).

| Parameter | Ancestral strain |  | Hot strain |  | Cold strain |  |
| --- | --- | --- | --- | --- | --- | --- |
| | n | Mean ± SE | n | Mean ± SE | n | Mean ± SE |
| **Preadult duration (d)** | 69 | 13.40 ± 0.04a | 71 | 11.38 ± 0.04c | 60 | 12.20 ± 0.06b |
| **Female longevity (d)** | 29 | 8.66 ± 0.52a | 37 | 6.22 ± 0.25b | 30 | 9.52 ± 0.80a |
| **Male longevity (d)** | 34 | 9.22 ± 0.40b | 31 | 8.45 ± 0.25b | 22 | 11.91 ± 0.84a |
| **Oviposition days (d)** | 29 | 7.88 ± 0.44a | 37 | 6.07 ± 0.27b | 30 | 8.33 ± 0.64a |
| **Female fecundity (eggs/female)** | 29 | 164.14 ± 4.77b | 37 | 175.62 ± 3.18a | 30 | 163.5 ± 2.12b |
| **$r$ (d<sup>-1</sup>)** | 90 | 0.257 ± 0.010b | 90 | 0.323 ± 0.010a | 90 | 0.280 ± 0.010b |
| **$R_0$ (eggs/female)** | 90 | 52.905 ± 8.223a | 90 | 72.106 ± 9.216a | 90 | 54.483 ± 8.164a |
| **$T$ (d)** | 90 | 15.388 ± 0.029a | 90 | 13.209 ± 0.022c | 90 | 14.213 ± 0.001b |
| **$\lambda$ (d<sup>-1</sup>)** | 90 | 1.293 ± 0.0135b | 90 | 1.382 ± 0.014a | 90 | 1.324 ± 0.014b |

Note: ' $r$ ' indicates intrinsic rate of increase; ' $R_0$ ' indicates net reproductive rate; ' $T$ ' indicates mean generation time; ' $\lambda$ ' indicates finite rate. Different letters indicate significant differences between strains ( $P < 0.05$ ).

**Supplementary File 2.**
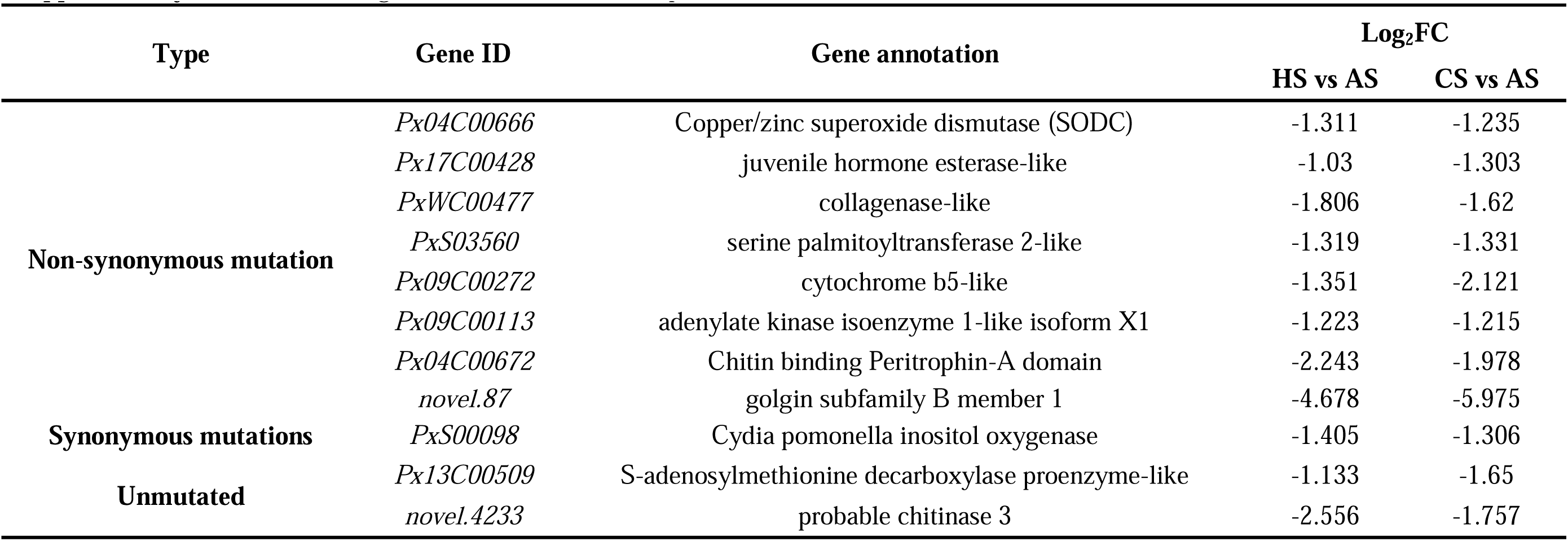
Candidate genes linked to thermal adaptation in DBM.

**Supplementary File 3.**
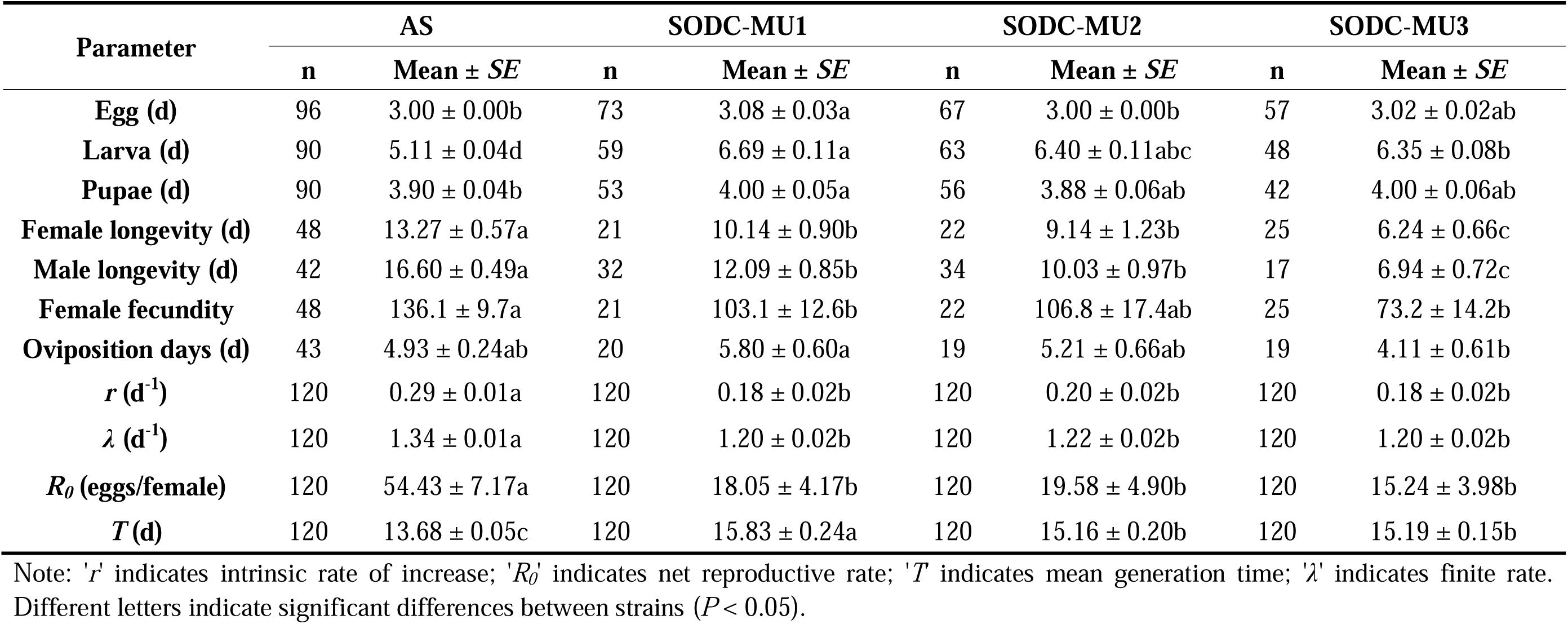
Population fitness parameters of the ancestral strain (AS) and SODC-mutant strains of DBM under the constant favorable environment (26°C).

| Parameter | AS |  | SODC-MU1 |  | SODC-MU2 |  | SODC-MU3 |  |
| --- | --- | --- | --- | --- | --- | --- | --- | --- |
| | n | Mean ± SE | n | Mean ± SE | n | Mean ± SE | n | Mean ± SE |
| **Egg (d)** | 96 | 3.00 ± 0.00b | 73 | 3.08 ± 0.03a | 67 | 3.00 ± 0.00b | 57 | 3.02 ± 0.02ab |
| **Larva (d)** | 90 | 5.11 ± 0.04d | 59 | 6.69 ± 0.11a | 63 | 6.40 ± 0.11abc | 48 | 6.35 ± 0.08b |
| **Pupae (d)** | 90 | 3.90 ± 0.04b | 53 | 4.00 ± 0.05a | 56 | 3.88 ± 0.06ab | 42 | 4.00 ± 0.06ab |
| **Female longevity (d)** | 48 | 13.27 ± 0.57a | 21 | 10.14 ± 0.90b | 22 | 9.14 ± 1.23b | 25 | 6.24 ± 0.66c |
| **Male longevity (d)** | 42 | 16.60 ± 0.49a | 32 | 12.09 ± 0.85b | 34 | 10.03 ± 0.97b | 17 | 6.94 ± 0.72c |
| **Female fecundity** | 48 | 136.1 ± 9.7a | 21 | 103.1 ± 12.6b | 22 | 106.8 ± 17.4ab | 25 | 73.2 ± 14.2b |
| **Oviposition days (d)** | 43 | 4.93 ± 0.24ab | 20 | 5.80 ± 0.60a | 19 | 5.21 ± 0.66ab | 19 | 4.11 ± 0.61b |
| **$r$ (d<sup>-1</sup>)** | 120 | 0.29 ± 0.01a | 120 | 0.18 ± 0.02b | 120 | 0.20 ± 0.02b | 120 | 0.18 ± 0.02b |
| **$\lambda$ (d<sup>-1</sup>)** | 120 | 1.34 ± 0.01a | 120 | 1.20 ± 0.02b | 120 | 1.22 ± 0.02b | 120 | 1.20 ± 0.02b |
| **$R_0$ (eggs/female)** | 120 | 54.43 ± 7.17a | 120 | 18.05 ± 4.17b | 120 | 19.58 ± 4.90b | 120 | 15.24 ± 3.98b |
| **$T$ (d)** | 120 | 13.68 ± 0.05c | 120 | 15.83 ± 0.24a | 120 | 15.16 ± 0.20b | 120 | 15.19 ± 0.15b |

Note: ' $r$ ' indicates intrinsic rate of increase; ' $R_0$ ' indicates net reproductive rate; ' $T$ ' indicates mean generation time; ' $\lambda$ ' indicates finite rate. Different letters indicate significant differences between strains ( $P < 0.05$ ).

**Supplementary File 4.** Populaion fitness parameters of the ancestral strain (AS) and mutant strains of DBM under the hot environment (32°C/27°C:12 h/12 h).

| Parameter | AS |  | SODC-MU1 |  | SODC-MU2 |  | SODC-MU3 |  |
| --- | --- | --- | --- | --- | --- | --- | --- | --- |
| | n | Mean ± SE | n | Mean ± SE | n | Mean ± SE | n | Mean ± SE |
| **Egg (d)** | 63 | 3.00 ± 0.00b | 42 | 3.29 ± 0.08a | 60 | 3.05 ± 0.03b | 63 | 3.00 ± 0.00b |
| **Larva (d)** | 57 | 5.18 ± 0.06c | 24 | 7.33 ± 0.23a | 46 | 6.91 ± 0.13a | 52 | 6.40 ± 0.09b |
| **Pupae (d)** | 57 | 3.51 ± 0.07bc | 22 | 3.91 ± 0.09a | 35 | 3.37 ± 0.08c | 44 | 3.61 ± 0.07b |
| **Female longevity (d)** | 30 | 5.83 ± 0.50b | 11 | 9.09 ± 0.67a | 20 | 5.30 ± 0.95b | 21 | 5.43 ± 0.94b |
| **Male longevity (d)** | 27 | 6.85 ± 0.51ab | 11 | 8.73 ± 1.29a | 15 | 7.67 ± 1.61ab | 23 | 4.43 ± 0.76b |
| **Female fecundity** | 30 | 62.7 ± 9.6a | 11 | 54.0 ± 11.7b | 20 | 25.8 ± 7.0c | 21 | 31.3 ± 8.7bc |
| **Oviposition days (d)** | 23 | 3.65 ± 0.29a | 11 | 4.09 ± 0.71a | 13 | 1.92 ± 0.26b | 11 | 3.09 ± 0.56ab |
| **$r$ (d<sup>-1</sup>)** | 120 | 0.20 ± 0.02a | 120 | 0.10 ± 0.03b | 120 | 0.10 ± 0.03b | 120 | 0.12 ± 0.02b |
| **$\lambda$ (d<sup>-1</sup>)** | 120 | 1.22 ± 0.02a | 120 | 1.11 ± 0.03b | 120 | 1.11 ± 0.03b | 120 | 1.12 ± 0.03b |
| **$R_0$ (eggs/female)** | 120 | 15.68 ± 3.44a | 120 | 4.95 ± 1.75b | 120 | 4.30 ± 1.43b | 120 | 5.48 ± 1.84b |
| **$T$ (d)** | 120 | 13.57 ± 0.13c | 120 | 15.91 ± 0.47a | 120 | 14.50 ± 0.46b | 120 | 14.78 ± 0.34b |

Note: ' $r$ ' indicates intrinsic rate of increase; ' $R_0$ ' indicates net reproductive rate; ' $T$ ' indicates mean generation time; ' $\lambda$ ' indicates finite rate. Different letters indicate significant differences between strains under the same temperature conditions ( $P < 0.05$ ).

**Supplementary File 5.**
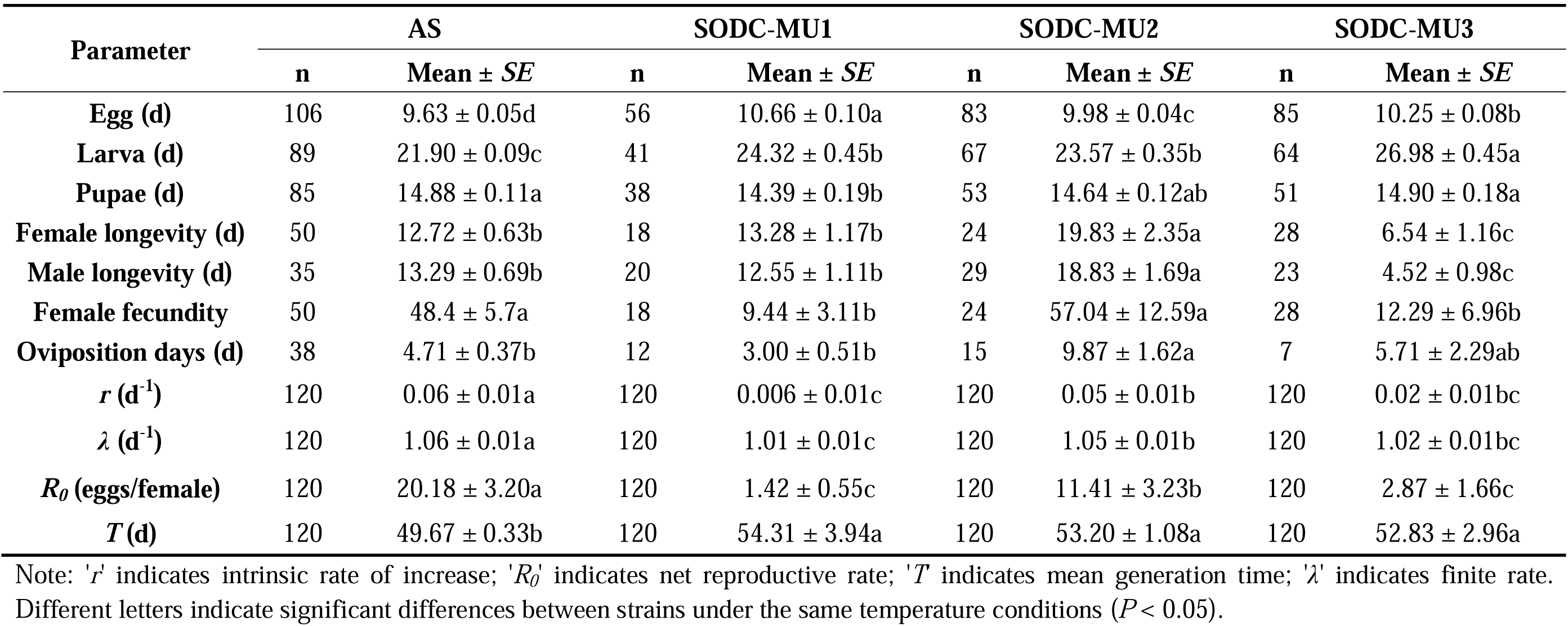
Population fitness parameters of the ancestral strain (AS) and mutant strains of DBM under the cold environment (15°C/10°C:12 h/12 h).

| Parameter | AS |  | SODC-MU1 |  | SODC-MU2 |  | SODC-MU3 |  |
| --- | --- | --- | --- | --- | --- | --- | --- | --- |
| | n | Mean ± SE | n | Mean ± SE | n | Mean ± SE | n | Mean ± SE |
| **Egg (d)** | 106 | 9.63 ± 0.05d | 56 | 10.66 ± 0.10a | 83 | 9.98 ± 0.04c | 85 | 10.25 ± 0.08b |
| **Larva (d)** | 89 | 21.90 ± 0.09c | 41 | 24.32 ± 0.45b | 67 | 23.57 ± 0.35b | 64 | 26.98 ± 0.45a |
| **Pupae (d)** | 85 | 14.88 ± 0.11a | 38 | 14.39 ± 0.19b | 53 | 14.64 ± 0.12ab | 51 | 14.90 ± 0.18a |
| **Female longevity (d)** | 50 | 12.72 ± 0.63b | 18 | 13.28 ± 1.17b | 24 | 19.83 ± 2.35a | 28 | 6.54 ± 1.16c |
| **Male longevity (d)** | 35 | 13.29 ± 0.69b | 20 | 12.55 ± 1.11b | 29 | 18.83 ± 1.69a | 23 | 4.52 ± 0.98c |
| **Female fecundity** | 50 | 48.4 ± 5.7a | 18 | 9.44 ± 3.11b | 24 | 57.04 ± 12.59a | 28 | 12.29 ± 6.96b |
| **Oviposition days (d)** | 38 | 4.71 ± 0.37b | 12 | 3.00 ± 0.51b | 15 | 9.87 ± 1.62a | 7 | 5.71 ± 2.29ab |
| **$r$ (d<sup>-1</sup>)** | 120 | 0.06 ± 0.01a | 120 | 0.006 ± 0.01c | 120 | 0.05 ± 0.01b | 120 | 0.02 ± 0.01bc |
| **$\lambda$ (d<sup>-1</sup>)** | 120 | 1.06 ± 0.01a | 120 | 1.01 ± 0.01c | 120 | 1.05 ± 0.01b | 120 | 1.02 ± 0.01bc |
| **$R_0$ (eggs/female)** | 120 | 20.18 ± 3.20a | 120 | 1.42 ± 0.55c | 120 | 11.41 ± 3.23b | 120 | 2.87 ± 1.66c |
| **$T$ (d)** | 120 | 49.67 ± 0.33b | 120 | 54.31 ± 3.94a | 120 | 53.20 ± 1.08a | 120 | 52.83 ± 2.96a |

Note: ' $r$ ' indicates intrinsic rate of increase; ' $R_0$ ' indicates net reproductive rate; ' $T$ ' indicates mean generation time; ' $\lambda$ ' indicates finite rate. Different letters indicate significant differences between strains under the same temperature conditions ( $P < 0.05$ ).

**Supplementary File 6.**
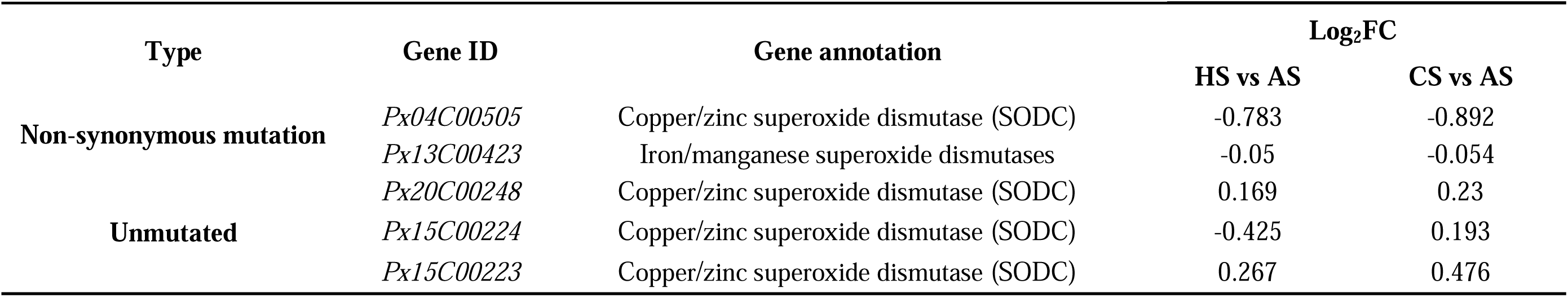
Information on the *PxSODC* homologous genes identified in the transcriptome.

**Supplementary File 7.**
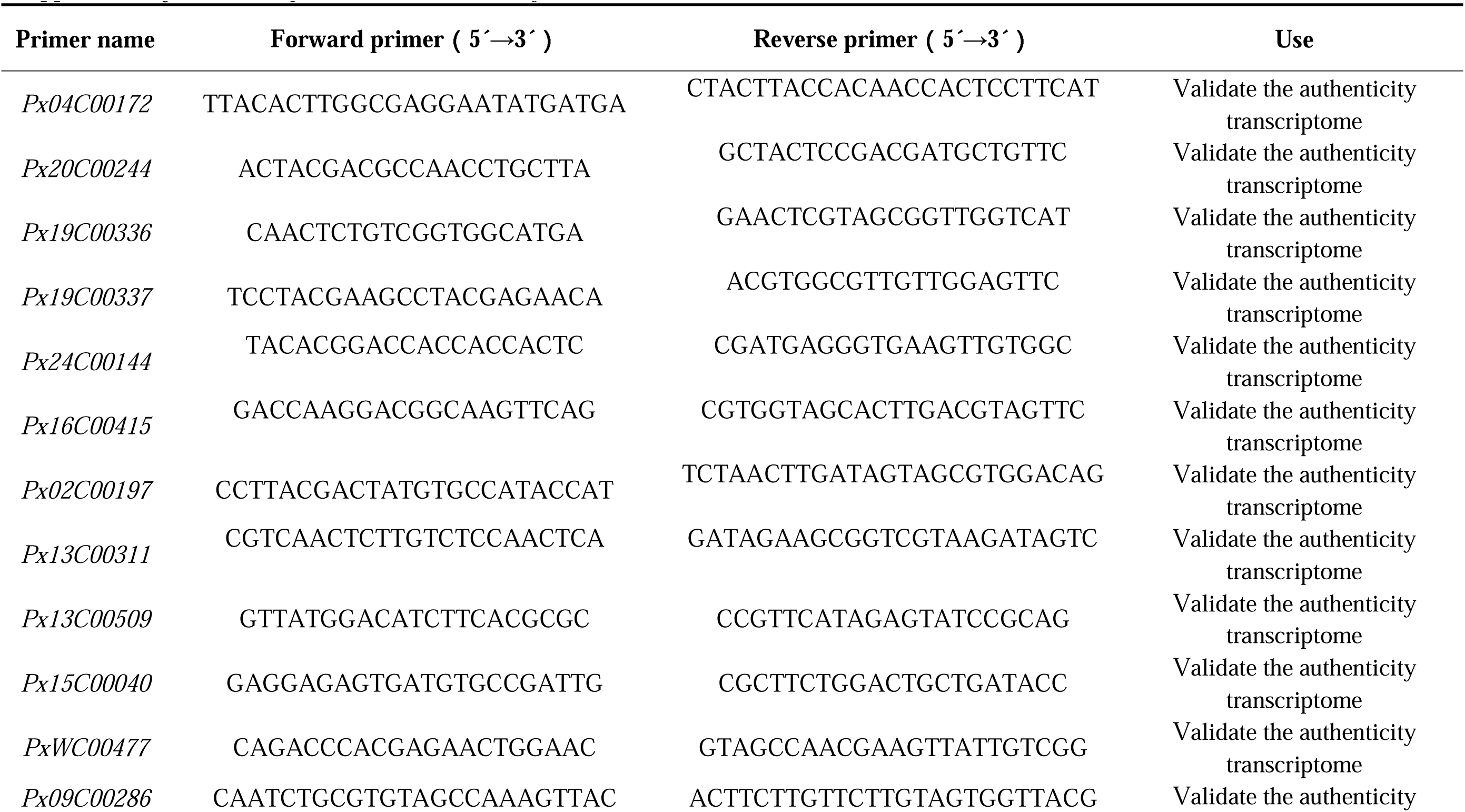
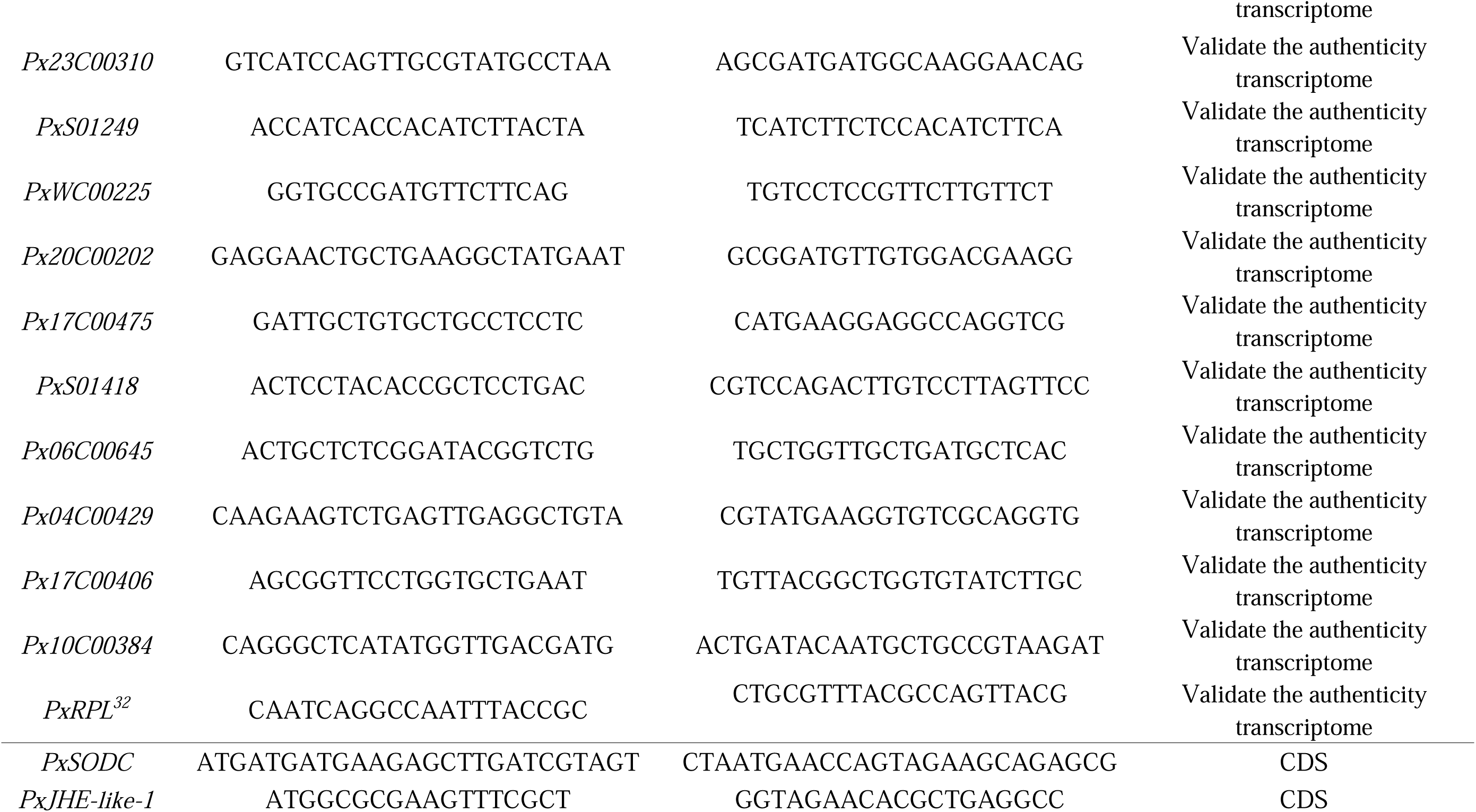
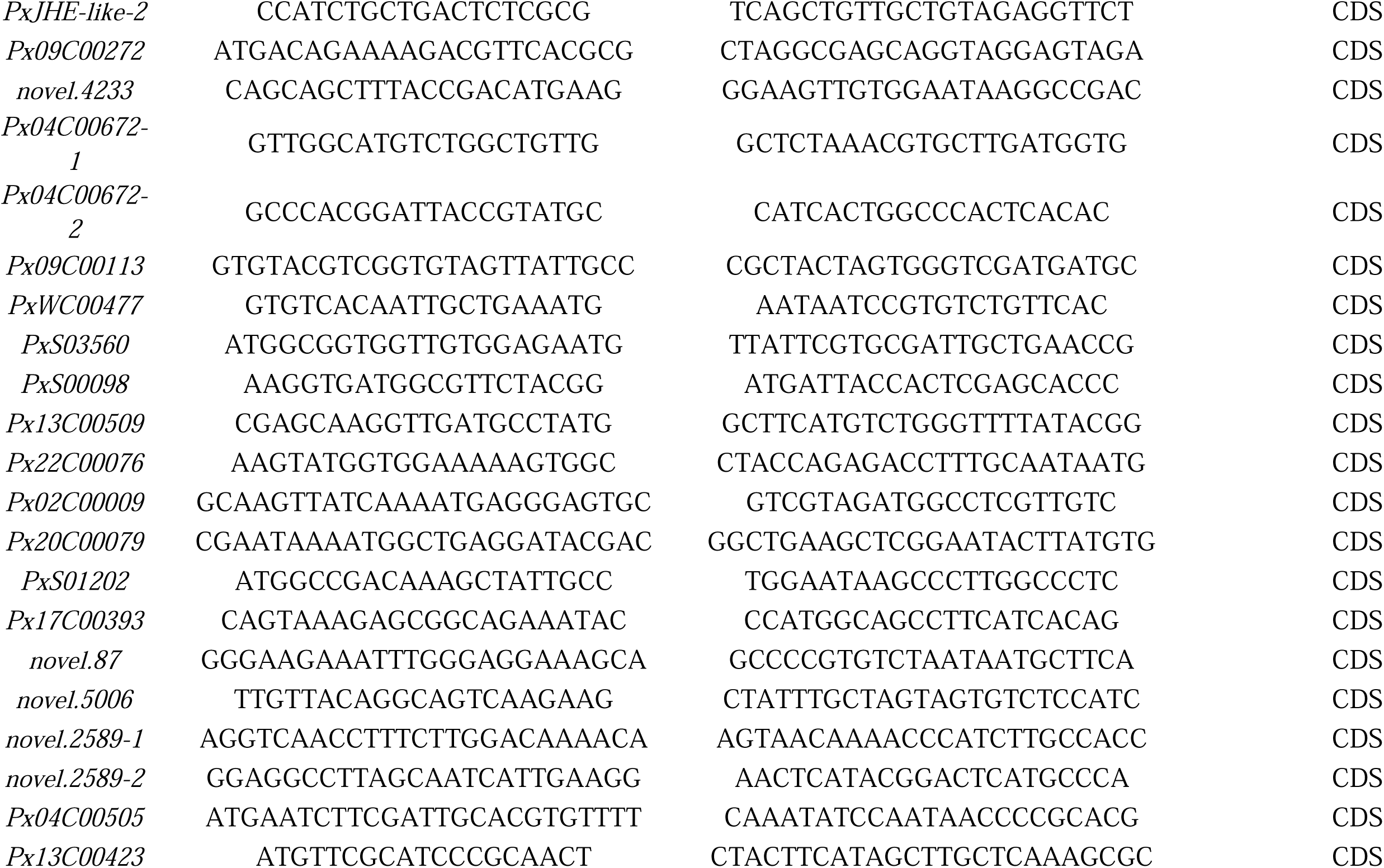
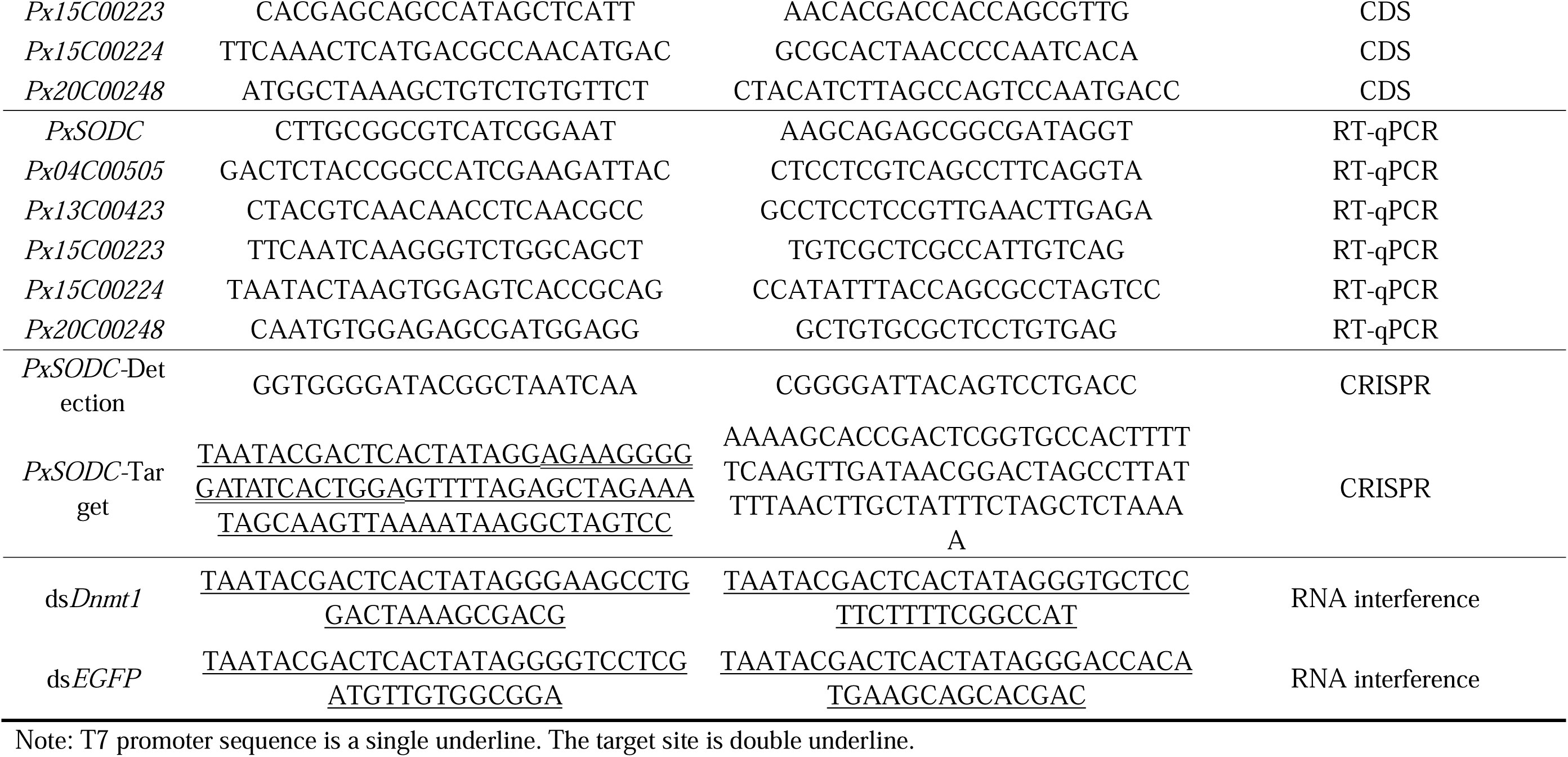
The primers used in this study.

## Notes

### Competing Interest Statement

The authors have declared no competing interest.

### Summary of Updates

This version of the manuscript is identical in scientific content to the previously revised version. The files have been reformatted to comply with eLife submission requirements for the Version of Record. Specific updates include: reorganization of the end-matter into Acknowledgements, Additional information (Funding table, Author contributions, Author ORCIDs, Competing interests), Additional files, and Data availability sections; conversion of funding information into the eLife table format; addition of source data file listings under each main figure legend; and standardization of figure supplement labels. No changes were made to the data, results, or conclusions.

## References

Bale JS, Masters GJ, Hodkinson ID, Awmack C, Bezemer TM, Brown VK, Butterfield J, Buse A, Coulson JC, Farrar J. 2002. Herbivory in global climate change research: direct effects of rising temperature on insect herbivores. Glob Change Biol 8: 1–16.

Barros-Cordeiro KB, Bao SN, Pujol-Luz JR. 2014. Intra-puparial development of the black soldier-fly, *Hermetia illucens*. J Insect Sci 14: 83.

Belfield EJ, Ding ZJ, Jamieson FJC, Visscher AM, Zheng SJ, Mithani A, Harberd NP. 2018. DNA mismatch repair preferentially protects genes from mutation. Genome Res 28: 66–74.

Benjamini Y, Hochberg Y. 1995. Controlling the false discovery rate: A practical and powerful approach to multiple testing. J R Stat Soc Ser B 57: 289–300.

Bhutani N, Burns David M, Blau Helen M. 2011. DNA demethylation dynamics. Cell 146: 866–872.

Bittner N, Hundacker J, Achotegui-Castells A, Anderbrant O, Hilker M. 2019. Defense of Scots pine against sawfly eggs (*Diprion pini*) is primed by exposure to sawfly sex pheromones. Proc Natl Acad Sci U S A 116: 24668–24675.

Block W. 1997. Cold tolerance of insects and other arthropods. Phil Trans R. Soc Lond B Biol Sci 326: 613–633.

Bremer J. 1983. Carnitine-metabolism and functions. Physiol Rev 63: 1420–1480.

Burc E, Girard-Tercieux C, Metz M, Cazaux E, Baur J, Koppik M, Rêgo A, Hart AF, Berger D. 2025. Life-history adaptation under climate warming magnifies the agricultural footprint of a cosmopolitan insect pest. Nat Commun 16: 827.

Chen Y, Liu Z, Régnière J, Vasseur L, Lin J, Huang S, Ke F, Chen S, Li J, Huang J. 2021. Large-scale genome-wide study reveals climate adaptive variability in a cosmopolitan pest. Nat Commun 12: 1–11.

Chi H. 1988. Life-table analysis incorporating both sexes and variable development rates among individuals. Env Entomol 17: 26–34.

Chi H, You M, Atlıhan R, Smith CL, Kavousi A, Özgökçe MS, Güncan A, Tuan SJ, Fu JW, Xu YY et al. 2020. Age-Stage, two-sex life table: an introduction to theory, data analysis, and application. Entomol Gen 40: 103–124.

Deutsch CA, Tewksbury JJ, Huey RB, Sheldon KS, Ghalambor CK, Haak DC, Martin PR. 2008. Impacts of climate warming on terrestrial ectotherms across latitude. Proc Natl Acad Sci U S A 105: 6668–6672.

Emre I, Kayis T, Coskun M, Dursun O, Cogun HY. 2013. Changes in antioxidative enzyme activity, glycogen, lipid, protein, and malondialdehyde content in cadmium-treated *Galleria mellonella* Larvae. Ann. Entomol. Soc Am 106: 371–377.

Furlong MJ, Wright DJ, Dosdall LM. 2013. Diamondback moth ecology and management: problems, progress, and prospects. Annu Rev Entomol 58: 517–541.

Gibert P, Debat V, Ghalambor CK. 2019. Phenotypic plasticity, global change, and the speed of adaptive evolution. Curr Opin Insect Sci 35: 34–40.

Gibson G, Barghi N, Tobler R, Nolte V, Jakšić AM, Mallard F, Otte KA, Dolezal M, Taus T, Kofler R, Schlötterer C. 2019. Genetic redundancy fuels polygenic adaptation in *Drosophila*. PLoS Biol 17: 3000128.

Halsch CA, Shapiro AM, Fordyce JA, Nice CC, Thorne JH, Waetjen DP, Forister ML. 2021. Insects and recent climate change. Proc Natl Acad Sci U S A 118: e2002543117.

Harvey JA, Heinen R, Gols R, Thakur MP. 2020. Climate changeCmediated temperature extremes and insects: From outbreaks to breakdowns. Glob Change Biol 26: 6685–6701.

Hoffmann AA, Sgro CM. 2011. Climate change and evolutionary adaptation. Nature 470: 479–485.

IPCC. 2023. Climate Change 2023: Synthesis Report. IPCC, Geneva, Switzerland.

Islam MN, Rauf A, Fahad FI, Emran TB, Mitra S, Olatunde A, Shariati MA, Rebezov M, Rengasamy KRR, Mubarak MS. 2022. Superoxide dismutase: an updated review on its health benefits and industrial applications. Crit Rev Food Sci Nutr 62: 7282–7300.

Kaiser A, Hartzendorf S, Wobschall A, Hetz SK. 2010. Modulation of cyclic CO(2) release in response to endogenous changes of metabolism during pupal development of *Zophobas rugipes* (Coleoptera: Tenebrionidae). J Insect Physiol 56: 502–512.

Langfelder P, Horvath S. 2008. WGCNA: an R package for weighted correlation network analysis. BMC Bioinform 9: 559.

Lawlor JA, Comte L, Grenouillet G, Lenoir J, Baecher JA, Bandara RMWJ, Bertrand R, Chen IC, Diamond SE, Lancaster LT et al. 2024. Mechanisms, detection and impacts of species redistributions under climate change. Nat Rev Earth Environ 5: 351–368.

Lei G, Huang J, Zhou H, Chen Y, Song J, Xie X, Vasseur L, You M, You S. 2024. Polygenic adaptation of a cosmopolitan pest to a novel thermal environment. Insect Mol Biol 33: 387–404.

Lei G, Zhou H, Chen Y, Vasseur L, Gurr GM, You M, You S. 2023. A very long-chain fatty acid enzyme gene, PxHacd2 affects the temperature adaptability of a cosmopolitan insect by altering epidermal permeability. Sci Total Environ 891: 164372.

Li J, Holford P, Beattie GAC, Wu S, He J, Tan S, Wang D, He Y, Cen Y, Nian X. 2024a. Adipokinetic hormone signaling mediates the enhanced fecundity of *Diaphorina citri* infected by ‘Candidatus Liberibacter asiaticus’. eLife 13: RP93450.

Li S, Deng B, Tian S, Guo M, Liu H, Zhao X. 2021. Metabolic and transcriptomic analyses reveal different metabolite biosynthesis profiles between leaf buds and mature leaves in *Ziziphus jujuba* mill. Food Chem 347: 129005.

Li T, Guo J, Hu G, Cao F, Su H, Shen M, Wang H, You M, Liu Y, Gurr GM, You S. 2024b. Zinc finger proteins facilitate adaptation of a global insect pest to climate change. BMC Biol 22: 303.

Li Y, Song H, Xie L, Tang X, Jiang Y, Yao Y, Peng X, Cui J, Zhou Z, Xu J. 2024c. Surviving high temperatures: The crucial role of vesicular inhibitory amino acid transporter in Asian honeybee, *Apis cerana*. Int J Biol Macromol 279: 135276.

Liu SS, Chen FZ, Zalucki MP. 2002. Development and survival of the diamondback moth (Lepidoptera: Plutellidae) at constant and alternating temperatures. Environ Entomol 31: 221–231.

Mallard F, Nolte V, Tobler R, Kapun M, Schlotterer C. 2018. A simple genetic basis of adaptation to a novel thermal environment results in complex metabolic rewiring in *Drosophila*. Genome Biol 19: 119.

McCulloch GA, Waters JM. 2022. Rapid adaptation in a fastCchanging world: Emerging insights from insect genomics. Glob Change Biol 29: 943–954.

McKean IJW, Sadler JC, Cuetos A, Frese A, Humphreys LD, Grogan G, Hoskisson PA, Burley GA. 2019. S-adenosyl methionine cofactor modifications enhance the biocatalytic repertoire of small molecule c-alkylation. Angew Chem Int Ed Engl 58: 17583–17588.

Mondola P, Damiano S, Sasso A, Santillo M. 2016. The Cu, Zn superoxide dismutase: not only a dismutase enzyme. Front Physiol 7: 594.

Ni P, Nie F, Zhong Z, Xu J, Huang N, Zhang J, Zhao H, Zou Y, Huang Y, Li J et al. 2023. DNA 5-methylcytosine detection and methylation phasing using PacBio circular consensus sequencing. Nat Commun 14: 4054.

Nobeli I, Favia AD, Thornton JM. 2009. Protein promiscuity and its implications for biotechnology. Nat Biotechnol 27: 157–167.

Outhwaite CL, McCann P, Newbold T. 2022. Agriculture and climate change are reshaping insect biodiversity worldwide. Nature 605: 97–102.

Patsch D, Schwander T, Voss M, Schaub D, Hüppi S, Eichenberger M, Stockinger P, Schelbert L, Giger S, Peccati F et al. 2024. Enriching productive mutational paths accelerates enzyme evolution. Nat Chem Biol 20: 1662–1669.

Quan PQ, Guo PL, He J, Liu XD. 2024. Heat-stress memory enhances the acclimation of a migratory insect pest to global warming. Mol Ecol 33: e17493.

Reynolds JA. 2017. Epigenetic influences on Diapause. In Advances in Insect Physiology, Vol 53 (ed. H Verlinden), pp. 115-144. Academic Press.

Rommelaere S, Boquete JP, Piton J, Kondo S, Lemaitre B. 2019. The exchangeable apolipoprotein Nplp2 sustains lipid flow and heat acclimation in *Drosophila*. Cell Rep 27: 886–899.

Schoville SD, Farrand Z, Kavanaugh DH, Veire B, Weng YM. 2024. Environmental stress responses and adaptive evolution in the alpine ground beetle *Nebria vandykei*. Biol J Linn Soc 141: 51–70.

Sheng Y, Abreu IA, Cabelli DE, Maroney MJ, Miller AF, Teixeira M, Valentine JS. 2014. Superoxide dismutases and superoxide reductases. Chem Rev 114: 3854–3918.

Sheng Y, Durazo A, Schumacher M, Gralla EB, Cascio D, Cabelli DE, Valentine JS. 2013. Tetramerization reinforces the dimer interface of MnSOD. PLoS One 8: e62446.

Sherpa S, Tutagata J, Gaude T, Laporte F, Kasai S, Ishak IH, Guo X, Shin J, Boyer S, Marcombe S et al. 2022. Genomic shifts, phenotypic clines, and fitness costs associated with cold tolerance in the Asian tiger mosquito. Mol Biol Evol 39: 104.

Stroud H, Song Q, Guan X, Chen ZJ. 2015. Dynamic roles for small RNAs and DNA methylation during ovule and fiber development in allotetraploid cotton. PLoS Genet 11: 1005724.

Tigano A, Colella JP, MacManes MD. 2020. Comparative and population genomics approaches reveal the basis of adaptation to deserts in a small rodent. Mol Ecol 29: 1300–1314.

Wang P, Shi S, Ma J, Song H, Zhang Y, Gao C, Zhao C, Zhao S, Hou L, Lopez-Baltazar J et al. 2018. Global methylome and gene expression analysis during early Peanut pod development. BMC Plant Biol 18: 352.

Wang Y, Huang Y, Xu X, Liu Z, Li J, Zhan X, Yang G, You M, You S. 2020. CRISPR/Cas9Cbased functional analysis of yellow gene in the diamondback moth, *Plutella xylostella*. Insect Sci 28: 1504–1509.

Wang Z, Receveur JP, Pu J, Cong H, Richards C, Liang M, Chung H. 2022. Desiccation resistance differences in *Drosophila* species can be largely explained by variations in cuticular hydrocarbons. eLife 11: 80859.

You M, Ke F, You S, Wu Z, Liu Q, He W, Baxter SW, Yuchi Z, Vasseur L, Gurr GM. 2020. Variation among 532 genomes unveils the origin and evolutionary history of a global insect herbivore. Nat Commun 11: 1–8.

You M, Yue Z, He W, Yang X, Yang G, Xie M, Zhan D, Baxter SW, Vasseur L, Gurr GM et al. 2013. A heterozygous moth genome provides insights into herbivory and detoxification. Nat Genet 45: 220–225.

You S, Lei G, Zhou H, Li J, Chen S, Huang J, Vasseur L, Gurr GM, You M, Chen Y. 2024. Thermal acclimation uncovers a simple genetic basis of adaptation to high temperature in a cosmopolitan pest. iScience 27: 109242.

Zhou H, Lei G, Chen Y, You M, You S. 2022. *PxTret1-like* affects the temperature adaptability of a cosmopolitan pest by altering trehalose tissue distribution. Int J Mol Sci 23: 9019.

Zhou H, Lei G, Li Y, Chen P, Liu Z, Li C, Li B. 2024. Novel regulation pathway of eclosion hormones in *Tribolium castaneum* by distinct transcription factors through the initiation of 20-hydroxyecdysone. J Biol Chem 300: 107898.

