## Supplementary File 1 for "Experimental evolution to thermal stress indicates climate resilience in a cosmopolitan arthropod"

**Supplementary File 1** Life table parameters of the ancestral, hot and cold strains of *P. xylostella* at the favorable temperature (26°C).

| **Parameter** | **Ancestral strain** | | **Hot strain** | | **Cold strain** | |
| --- | --- | --- | --- | --- | --- | --- |
|  | **n** | **Mean ± *SE*** | **n** | **Mean ± *SE*** | **n** | **Mean ± *SE*** |
| **Preadult duration (d)** | 69 | 13.40 ± 0.04a | 71 | 11.38 ± 0.04c | 60 | 12.20 ± 0.06b |
| **Female longevity (d)** | 29 | 8.66 ± 0.52a | 37 | 6.22 ± 0.25b | 30 | 9.52 ± 0.80a |
| **Male longevity (d)** | 34 | 9.22 ± 0.40b | 31 | 8.45 ± 0.25b | 22 | 11.91 ± 0.84a |
| **Oviposition days (d)** | 29 | 7.88 ± 0.44a | 37 | 6.07 ± 0.27b | 30 | 8.33 ± 0.64a |
| **Female fecundity (eggs/female)** | 29 | 164.14 ± 4.77b | 37 | 175.62 ± 3.18a | 30 | 163.5 ± 2.12b |
| ***r* (d^-1^)** | 90 | 0.257 ± 0.010b | 90 | 0.323 ± 0.010a | 90 | 0.280 ± 0.010b |
| ***R_0_* (eggs/female)** | 90 | 52.905 ± 8.223a | 90 | 72.106 ± 9.216a | 90 | 54.483 ± 8.164a |
| ***T* (d)** | 90 | 15.388 ± 0.029a | 90 | 13.209 ± 0.022c | 90 | 14.213 ± 0.001b |
| ***λ* (d^-1^)** | 90 | 1.293 ± 0.0135b | 90 | 1.382 ± 0.014a | 90 | 1.324 ± 0.014b |

Note: '*r*' indicates intrinsic rate of increase; '*R_0_*' indicates net reproductive rate; '*T*' indicates mean generation time; '*λ*' indicates finite rate. Different letters indicate significant differences between strains (*P* < 0.05).
