## Supplementary File 2 for "Experimental evolution to thermal stress indicates climate resilience in a cosmopolitan arthropod"

**Supplementary File 2** Candidate genes linked to thermal adaptation in DBM.

| **Type** | **Gene ID** | **Gene annotation** | Log_2_FC | |
| --- | --- | --- | --- | --- |
|  |  |  | **HS vs AS** | **CS vs AS** |
| **Non-synonymous mutation** | *Px04C00666* | Copper/zinc superoxide dismutase (SODC) | -1.311 | -1.235 |
|  | *Px17C00428* | juvenile hormone esterase-like | -1.03 | -1.303 |
|  | *PxWC00477* | collagenase-like | -1.806 | -1.62 |
|  | *PxS03560* | serine palmitoyltransferase 2-like | -1.319 | -1.331 |
|  | *Px09C00272* | cytochrome b5-like | -1.351 | -2.121 |
|  | *Px09C00113* | adenylate kinase isoenzyme 1-like isoform X1 | -1.223 | -1.215 |
|  | *Px04C00672* | Chitin binding Peritrophin-A domain | -2.243 | -1.978 |
|  | *novel.87* | golgin subfamily B member 1 | -4.678 | -5.975 |
| **Synonymous mutations** | *PxS00098* | Cydia pomonella inositol oxygenase | -1.405 | -1.306 |
| **Unmutated** | *Px13C00509* | S-adenosylmethionine decarboxylase proenzyme-like | -1.133 | -1.65 |
|  | *novel.4233* | probable chitinase 3 | -2.556 | -1.757 |
