## Supplementary File 3 for "Experimental evolution to thermal stress indicates climate resilience in a cosmopolitan arthropod"

**Supplementary File 3** Population fitness parameters of the ancestral strain (AS) and SODC-mutant strains of DBM under the constant favorable environment (26°C).

| **Parameter** | **AS** | | **SODC-MU1** | | **SODC-MU2** | | **SODC-MU3** | |
| --- | --- | --- | --- | --- | --- | --- | --- | --- |
|  | **n** | **Mean ± *SE*** | **n** | **Mean ± *SE*** | **n** | **Mean ± *SE*** | **n** | **Mean ± *SE*** |
| **Egg (d)** | 96 | 3.00 ± 0.00b | 73 | 3.08 ± 0.03a | 67 | 3.00 ± 0.00b | 57 | 3.02 ± 0.02ab |
| **Larva (d)** | 90 | 5.11 ± 0.04d | 59 | 6.69 ± 0.11a | 63 | 6.40 ± 0.11abc | 48 | 6.35 ± 0.08b |
| **Pupae (d)** | 90 | 3.90 ± 0.04b | 53 | 4.00 ± 0.05a | 56 | 3.88 ± 0.06ab | 42 | 4.00 ± 0.06ab |
| **Female longevity (d)** | 48 | 13.27 ± 0.57a | 21 | 10.14 ± 0.90b | 22 | 9.14 ± 1.23b | 25 | 6.24 ± 0.66c |
| **Male longevity (d)** | 42 | 16.60 ± 0.49a | 32 | 12.09 ± 0.85b | 34 | 10.03 ± 0.97b | 17 | 6.94 ± 0.72c |
| **Female fecundity** | 48 | 136.1 ± 9.7a | 21 | 103.1 ± 12.6b | 22 | 106.8 ± 17.4ab | 25 | 73.2 ± 14.2b |
| **Oviposition days (d)** | 43 | 4.93 ± 0.24ab | 20 | 5.80 ± 0.60a | 19 | 5.21 ± 0.66ab | 19 | 4.11 ± 0.61b |
| **r (d^-1^)** | 120 | 0.29 ± 0.01a | 120 | 0.18 ± 0.02b | 120 | 0.20 ± 0.02b | 120 | 0.18 ± 0.02b |
| **λ (d^-1^)** | 120 | 1.34 ± 0.01a | 120 | 1.20 ± 0.02b | 120 | 1.22 ± 0.02b | 120 | 1.20 ± 0.02b |
| **R_0_ (eggs/female)** | 120 | 54.43 ± 7.17a | 120 | 18.05 ± 4.17b | 120 | 19.58 ± 4.90b | 120 | 15.24 ± 3.98b |
| **T (d)** | 120 | 13.68 ± 0.05c | 120 | 15.83 ± 0.24a | 120 | 15.16 ± 0.20b | 120 | 15.19 ± 0.15b |

Note: '*r*' indicates intrinsic rate of increase; '*R_0_*' indicates net reproductive rate; '*T*' indicates mean generation time; '*λ*' indicates finite rate. Different letters indicate significant differences between strains (*P* < 0.05).
