## Supplementary File 4 for "Experimental evolution to thermal stress indicates climate resilience in a cosmopolitan arthropod"

**Supplementary File 4** Populaion fitness parameters of the ancestral strain (AS) and mutant strains of DBM under the hot environment (32°C/27°C:12 h/12 h).

| **Parameter** | **AS** | | **SODC-MU1** | | **SODC-MU2** | | **SODC-MU3** | |
| --- | --- | --- | --- | --- | --- | --- | --- | --- |
|  | **n** | **Mean ± *SE*** | **n** | **Mean ± *SE*** | **n** | **Mean ± *SE*** | **n** | **Mean ± *SE*** |
| **Egg (d)** | 63 | 3.00 ± 0.00b | 42 | 3.29 ± 0.08a | 60 | 3.05 ± 0.03b | 63 | 3.00 ± 0.00b |
| **Larva (d)** | 57 | 5.18 ± 0.06c | 24 | 7.33 ± 0.23a | 46 | 6.91 ± 0.13a | 52 | 6.40 ± 0.09b |
| **Pupae (d)** | 57 | 3.51 ± 0.07bc | 22 | 3.91 ± 0.09a | 35 | 3.37 ± 0.08c | 44 | 3.61 ± 0.07b |
| **Female longevity (d)** | 30 | 5.83 ± 0.50b | 11 | 9.09 ± 0.67a | 20 | 5.30 ± 0.95b | 21 | 5.43 ± 0.94b |
| **Male longevity (d)** | 27 | 6.85 ± 0.51ab | 11 | 8.73 ± 1.29a | 15 | 7.67 ± 1.61ab | 23 | 4.43 ± 0.76b |
| **Female fecundity** | 30 | 62.7 ± 9.6a | 11 | 54.0 ± 11.7b | 20 | 25.8 ± 7.0c | 21 | 31.3 ± 8.7bc |
| **Oviposition days (d)** | 23 | 3.65 ± 0.29a | 11 | 4.09 ± 0.71a | 13 | 1.92 ± 0.26b | 11 | 3.09 ± 0.56ab |
| ***r* (d^-1^)** | 120 | 0.20 ± 0.02a | 120 | 0.10 ± 0.03b | 120 | 0.10 ± 0.03b | 120 | 0.12 ± 0.02b |
| ***λ* (d^-1^)** | 120 | 1.22 ± 0.02a | 120 | 1.11 ± 0.03b | 120 | 1.11 ± 0.03b | 120 | 1.12 ± 0.03b |
| ***R_0_* (eggs/female)** | 120 | 15.68 ± 3.44a | 120 | 4.95 ± 1.75b | 120 | 4.30 ± 1.43b | 120 | 5.48 ± 1.84b |
| ***T* (d)** | 120 | 13.57 ± 0.13c | 120 | 15.91 ± 0.47a | 120 | 14.50 ± 0.46b | 120 | 14.78 ± 0.34b |

Note: '*r*' indicates intrinsic rate of increase; '*R_0_*' indicates net reproductive rate; '*T*' indicates mean generation time; '*λ*' indicates finite rate. Different letters indicate significant differences between strains under the same temperature conditions (*P* < 0.05).
