## Supplementary File 5 for "Experimental evolution to thermal stress indicates climate resilience in a cosmopolitan arthropod"

**Supplementary File 5** Population fitness parameters of the ancestral strain (AS) and mutant strains of DBM under the cold environment (15°C/10°C:12 h/12 h).

| **Parameter** | **AS** | | **SODC-MU1** | | **SODC-MU2** | | **SODC-MU3** | |
| --- | --- | --- | --- | --- | --- | --- | --- | --- |
|  | **n** | **Mean ± *SE*** | **n** | **Mean ± *SE*** | **n** | **Mean ± *SE*** | **n** | **Mean ± *SE*** |
| **Egg (d)** | 106 | 9.63 ± 0.05d | 56 | 10.66 ± 0.10a | 83 | 9.98 ± 0.04c | 85 | 10.25 ± 0.08b |
| **Larva (d)** | 89 | 21.90 ± 0.09c | 41 | 24.32 ± 0.45b | 67 | 23.57 ± 0.35b | 64 | 26.98 ± 0.45a |
| **Pupae (d)** | 85 | 14.88 ± 0.11a | 38 | 14.39 ± 0.19b | 53 | 14.64 ± 0.12ab | 51 | 14.90 ± 0.18a |
| **Female longevity (d)** | 50 | 12.72 ± 0.63b | 18 | 13.28 ± 1.17b | 24 | 19.83 ± 2.35a | 28 | 6.54 ± 1.16c |
| **Male longevity (d)** | 35 | 13.29 ± 0.69b | 20 | 12.55 ± 1.11b | 29 | 18.83 ± 1.69a | 23 | 4.52 ± 0.98c |
| **Female fecundity** | 50 | 48.4 ± 5.7a | 18 | 9.44 ± 3.11b | 24 | 57.04 ± 12.59a | 28 | 12.29 ± 6.96b |
| **Oviposition days (d)** | 38 | 4.71 ± 0.37b | 12 | 3.00 ± 0.51b | 15 | 9.87 ± 1.62a | 7 | 5.71 ± 2.29ab |
| ***r* (d^-1^)** | 120 | 0.06 ± 0.01a | 120 | 0.006 ± 0.01c | 120 | 0.05 ± 0.01b | 120 | 0.02 ± 0.01bc |
| ***λ* (d^-1^)** | 120 | 1.06 ± 0.01a | 120 | 1.01 ± 0.01c | 120 | 1.05 ± 0.01b | 120 | 1.02 ± 0.01bc |
| ***R_0_* (eggs/female)** | 120 | 20.18 ± 3.20a | 120 | 1.42 ± 0.55c | 120 | 11.41 ± 3.23b | 120 | 2.87 ± 1.66c |
| ***T* (d)** | 120 | 49.67 ± 0.33b | 120 | 54.31 ± 3.94a | 120 | 53.20 ± 1.08a | 120 | 52.83 ± 2.96a |

Note: '*r*' indicates intrinsic rate of increase; '*R_0_*' indicates net reproductive rate; '*T*' indicates mean generation time; '*λ*' indicates finite rate. Different letters indicate significant differences between strains under the same temperature conditions (*P* < 0.05).
