## Supplementary File 6 for "Experimental evolution to thermal stress indicates climate resilience in a cosmopolitan arthropod"

**Supplementary File 6** Information on the *PxSODC* homologous genes identified in the transcriptome.

| **Type** | **Gene ID** | **Gene annotation** | **Log_2_FC** | |
| --- | --- | --- | --- | --- |
|  |  |  | **HS vs AS** | **CS vs AS** |
| **Non-synonymous mutation** | *Px04C00505* | Copper/zinc superoxide dismutase (SODC) | -0.783 | -0.892 |
|  | *Px13C00423* | Iron/manganese superoxide dismutases | -0.05 | -0.054 |
| **Unmutated** | *Px20C00248* | Copper/zinc superoxide dismutase (SODC) | 0.169 | 0.23 |
|  | *Px15C00224* | Copper/zinc superoxide dismutase (SODC) | -0.425 | 0.193 |
|  | *Px15C00223* | Copper/zinc superoxide dismutase (SODC) | 0.267 | 0.476 |
