## Supplementary File 7 for "Experimental evolution to thermal stress indicates climate resilience in a cosmopolitan arthropod"

**Supplementary File 7** The primers used in this study.

| **Primer name** | **Forward primer**（**5´→3´**） | **Reverse primer**（**5´→3´**） | **Use** |
| --- | --- | --- | --- |
| *Px04C00172* | TTACACTTGGCGAGGAATATGATGA | CTACTTACCACAACCACTCCTTCAT | Validate the authenticity transcriptome |
| *Px20C00244* | ACTACGACGCCAACCTGCTTA | GCTACTCCGACGATGCTGTTC | Validate the authenticity transcriptome |
| *Px19C00336* | CAACTCTGTCGGTGGCATGA | GAACTCGTAGCGGTTGGTCAT | Validate the authenticity transcriptome |
| *Px19C00337* | TCCTACGAAGCCTACGAGAACA | ACGTGGCGTTGTTGGAGTTC | Validate the authenticity transcriptome |
| *Px24C00144* | TACACGGACCACCACCACTC | CGATGAGGGTGAAGTTGTGGC | Validate the authenticity transcriptome |
| *Px16C00415* | GACCAAGGACGGCAAGTTCAG | CGTGGTAGCACTTGACGTAGTTC | Validate the authenticity transcriptome |
| *Px02C00197* | CCTTACGACTATGTGCCATACCAT | TCTAACTTGATAGTAGCGTGGACAG | Validate the authenticity transcriptome |
| *Px13C00311* | CGTCAACTCTTGTCTCCAACTCA | GATAGAAGCGGTCGTAAGATAGTC | Validate the authenticity transcriptome |
| *Px13C00509* | GTTATGGACATCTTCACGCGC | CCGTTCATAGAGTATCCGCAG | Validate the authenticity transcriptome |
| *Px15C00040* | GAGGAGAGTGATGTGCCGATTG | CGCTTCTGGACTGCTGATACC | Validate the authenticity transcriptome |
| *PxWC00477* | CAGACCCACGAGAACTGGAAC | GTAGCCAACGAAGTTATTGTCGG | Validate the authenticity transcriptome |
| *Px09C00286* | CAATCTGCGTGTAGCCAAAGTTAC | ACTTCTTGTTCTTGTAGTGGTTACG | Validate the authenticity transcriptome |
| *Px23C00310* | GTCATCCAGTTGCGTATGCCTAA | AGCGATGATGGCAAGGAACAG | Validate the authenticity transcriptome |
| *PxS01249* | ACCATCACCACATCTTACTA | TCATCTTCTCCACATCTTCA | Validate the authenticity transcriptome |
| *PxWC00225* | GGTGCCGATGTTCTTCAG | TGTCCTCCGTTCTTGTTCT | Validate the authenticity transcriptome |
| *Px20C00202* | GAGGAACTGCTGAAGGCTATGAAT | GCGGATGTTGTGGACGAAGG | Validate the authenticity transcriptome |
| *Px17C00475* | GATTGCTGTGCTGCCTCCTC | CATGAAGGAGGCCAGGTCG | Validate the authenticity transcriptome |
| *PxS01418* | ACTCCTACACCGCTCCTGAC | CGTCCAGACTTGTCCTTAGTTCC | Validate the authenticity transcriptome |
| *Px06C00645* | ACTGCTCTCGGATACGGTCTG | TGCTGGTTGCTGATGCTCAC | Validate the authenticity transcriptome |
| *Px04C00429* | CAAGAAGTCTGAGTTGAGGCTGTA | CGTATGAAGGTGTCGCAGGTG | Validate the authenticity transcriptome |
| *Px17C00406* | AGCGGTTCCTGGTGCTGAAT | TGTTACGGCTGGTGTATCTTGC | Validate the authenticity transcriptome |
| *Px10C00384* | CAGGGCTCATATGGTTGACGATG | ACTGATACAATGCTGCCGTAAGAT | Validate the authenticity transcriptome |
| *PxRPL*^32^ | CAATCAGGCCAATTTACCGC | CTGCGTTTACGCCAGTTACG | Validate the authenticity transcriptome |
| *PxSODC* | ATGATGATGAAGAGCTTGATCGTAGT | CTAATGAACCAGTAGAAGCAGAGCG | CDS |
| *PxJHE-like-1* | ATGGCGCGAAGTTTCGCT | GGTAGAACACGCTGAGGCC | CDS |
| *PxJHE-like-2* | CCATCTGCTGACTCTCGCG | TCAGCTGTTGCTGTAGAGGTTCT | CDS |
| *Px09C00272* | ATGACAGAAAAGACGTTCACGCG | CTAGGCGAGCAGGTAGGAGTAGA | CDS |
| *novel.4233* | CAGCAGCTTTACCGACATGAAG | GGAAGTTGTGGAATAAGGCCGAC | CDS |
| *Px04C00672-1* | GTTGGCATGTCTGGCTGTTG | GCTCTAAACGTGCTTGATGGTG | CDS |
| *Px04C00672-2* | GCCCACGGATTACCGTATGC | CATCACTGGCCCACTCACAC | CDS |
| *Px09C00113* | GTGTACGTCGGTGTAGTTATTGCC | CGCTACTAGTGGGTCGATGATGC | CDS |
| *PxWC00477* | GTGTCACAATTGCTGAAATG | AATAATCCGTGTCTGTTCAC | CDS |
| *PxS03560* | ATGGCGGTGGTTGTGGAGAATG | TTATTCGTGCGATTGCTGAACCG | CDS |
| *PxS00098* | AAGGTGATGGCGTTCTACGG | ATGATTACCACTCGAGCACCC | CDS |
| *Px13C00509* | CGAGCAAGGTTGATGCCTATG | GCTTCATGTCTGGGTTTTATACGG | CDS |
| *Px22C00076* | AAGTATGGTGGAAAAAGTGGC | CTACCAGAGACCTTTGCAATAATG | CDS |
| *Px02C00009* | GCAAGTTATCAAAATGAGGGAGTGC | GTCGTAGATGGCCTCGTTGTC | CDS |
| *Px20C00079* | CGAATAAAATGGCTGAGGATACGAC | GGCTGAAGCTCGGAATACTTATGTG | CDS |
| *PxS01202* | ATGGCCGACAAAGCTATTGCC | TGGAATAAGCCCTTGGCCCTC | CDS |
| *Px17C00393* | CAGTAAAGAGCGGCAGAAATAC | CCATGGCAGCCTTCATCACAG | CDS |
| *novel.87* | GGGAAGAAATTTGGGAGGAAAGCA | GCCCCGTGTCTAATAATGCTTCA | CDS |
| *novel.5006* | TTGTTACAGGCAGTCAAGAAG | CTATTTGCTAGTAGTGTCTCCATC | CDS |
| *novel.2589-1* | AGGTCAACCTTTCTTGGACAAAACA | AGTAACAAAACCCATCTTGCCACC | CDS |
| *novel.2589-2* | GGAGGCCTTAGCAATCATTGAAGG | AACTCATACGGACTCATGCCCA | CDS |
| *Px04C00505* | ATGAATCTTCGATTGCACGTGTTTT | CAAATATCCAATAACCCCGCACG | CDS |
| *Px13C00423* | ATGTTCGCATCCCGCAACT | CTACTTCATAGCTTGCTCAAAGCGC | CDS |
| *Px15C00223* | CACGAGCAGCCATAGCTCATT | AACACGACCACCAGCGTTG | CDS |
| *Px15C00224* | TTCAAACTCATGACGCCAACATGAC | GCGCACTAACCCCAATCACA | CDS |
| *Px20C00248* | ATGGCTAAAGCTGTCTGTGTTCT | CTACATCTTAGCCAGTCCAATGACC | CDS |
| *PxSODC* | CTTGCGGCGTCATCGGAAT | AAGCAGAGCGGCGATAGGT | RT-qPCR |
| *Px04C00505* | GACTCTACCGGCCATCGAAGATTAC | CTCCTCGTCAGCCTTCAGGTA | RT-qPCR |
| *Px13C00423* | CTACGTCAACAACCTCAACGCC | GCCTCCTCCGTTGAACTTGAGA | RT-qPCR |
| *Px15C00223* | TTCAATCAAGGGTCTGGCAGCT | TGTCGCTCGCCATTGTCAG | RT-qPCR |
| *Px15C00224* | TAATACTAAGTGGAGTCACCGCAG | CCATATTTACCAGCGCCTAGTCC | RT-qPCR |
| *Px20C00248* | CAATGTGGAGAGCGATGGAGG | GCTGTGCGCTCCTGTGAG | RT-qPCR |
| *PxSODC-*Detection | GGTGGGGATACGGCTAATCAA | CGGGGATTACAGTCCTGACC | CRISPR |
| *PxSODC-*Target | TAATACGACTCACTATAGGAGAAGGGGGATATCACTGGAGTTTTAGAGCTAGAAATAGCAAGTTAAAATAAGGCTAGTCC | AAAAGCACCGACTCGGTGCCACTTTTTCAAGTTGATAACGGACTAGCCTTATTTTAACTTGCTATTTCTAGCTCTAAAA | CRISPR |
| ds*Dnmt1* | TAATACGACTCACTATAGGGAAGCCTGGACTAAAGCGACG | TAATACGACTCACTATAGGGTGCTCCTTCTTTTCGGCCAT | RNA interference |
| ds*EGFP* | TAATACGACTCACTATAGGGGTCCTCGATGTTGTGGCGGA | TAATACGACTCACTATAGGGACCACATGAAGCAGCACGAC | RNA interference |

Note: T7 promoter sequence is a single underline. The target site is double underline.
